# Prediction of biomolecule kinetics using physics-based Brownian dynamics to data-driven machine learning methods

**DOI:** 10.64898/2025.12.03.692248

**Authors:** Bin Sun, Alec Loftus, Peter Kekenes-Huskey

## Abstract

We review the Brownian dynamics (BD) modeling of biomolecular binding events, with an emphasis on enzyme-substrate interactions in cellular environments. We begin with theoretical foundations of BD and its applications to association and dissociation binding processes in both homogeneous and heterogeneous media. Next, we discuss BD in the context of continuum methods and emerging machine learning (ML) approaches toward predicting binding kinetics in vivo, with an eye toward multiscale modeling perspectives. Finally, we position BD simulations as a bridge to couple atomistic-scale models with dynamic, systems biology-based descriptions of cellular processes as a final frontier in modeling target/enzyme binding processes.

## 2 The Enzyme-substrate Interaction in Biological Systems

### 2.1 The Importance of Enzyme Kinetics

Enzymatic reactions are fundamentally important in dictating diverse biological functions; as a result, there has been an extensive focus on understanding their rates (kinetics) and molecular determinants. For example, the rapid excitability of neurons and muscle cells is in part shaped by the speed at which substrates diffuse from where they are synthesized or stored toward their cognate targets[1, 2]. Further, cell excitation, and diverse cellular responses as a whole, often arise from several enzyme-substrate and protein-protein association processes working in concert. Such coupled processes can frequently be decomposed into series of individual binding events, as we have illustrated for the multistep nucleotide degradation scheme in Figure 1, panel A. In this review, we discuss how Brownian dynamics (BD) simulations offer a powerful framework for probing the molecular mechanisms underlying these processes, providing a practical bridge between isolated binding events and their concerted behavior in biological systems.

**Figure 1:**
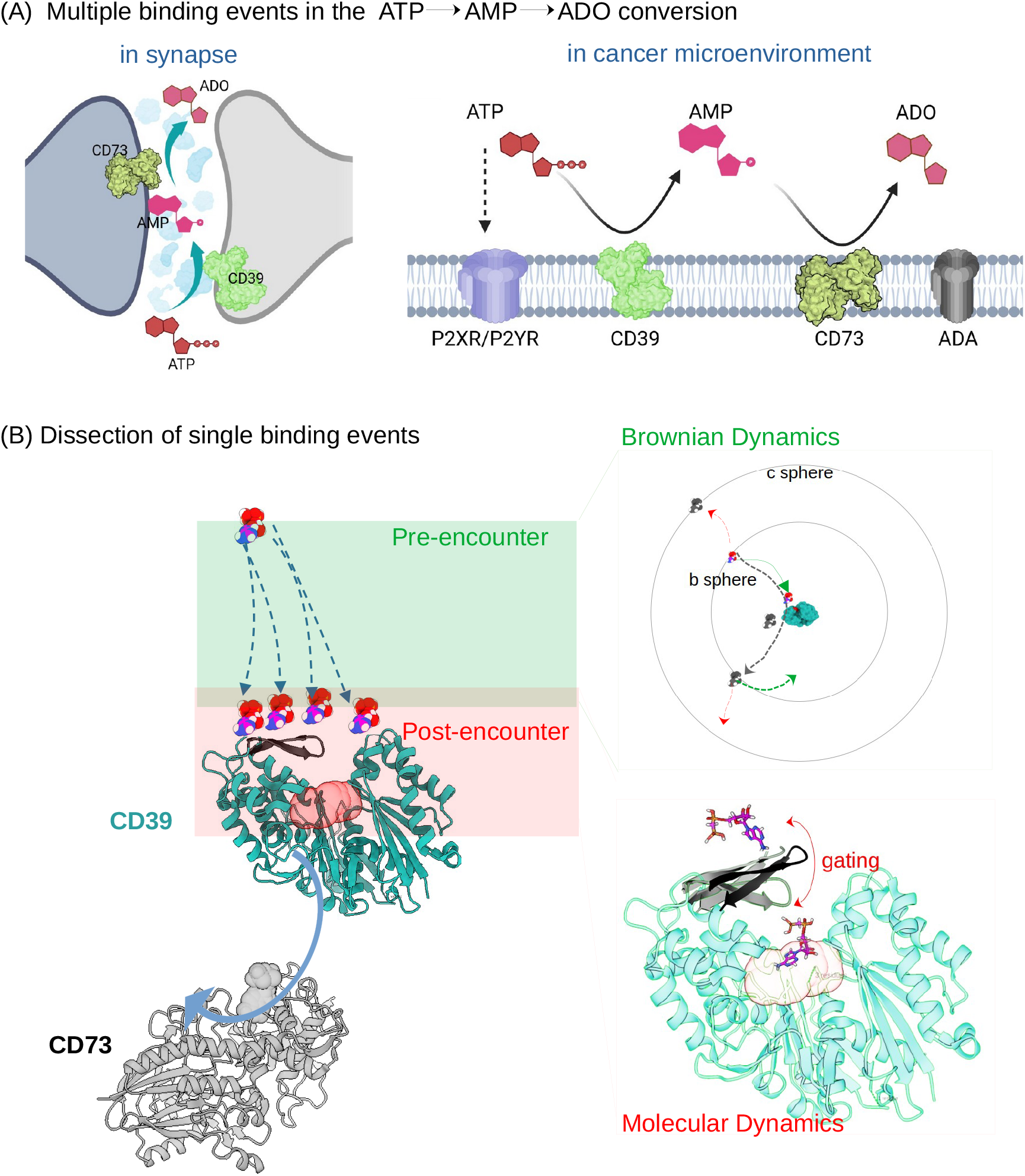
Multi-step reactions. (a) Sequential reactions or multiple binding events. The sequential conversion of ATP (or ADP) to AMP and back to ATP, catalyzed by membrane-embedded CD39 and CD73 enzymes, are important in synapse signaling [3] and cancer immunosuppression [4]. (b) Decomposition of a diffusion-limited single binding event into pre-and post-encounter stages. In this review, we discuss how Brownian dynamics simulations are well-suited for modeling the substrate diffusion to the transient-encounter surface interfacing the pre- and post-encounter regimes. BD provides computationally efficient evaluations of long-range electrostatic, solvation and hydrodynamic interaction influencing diffusion of rigid-body biomolecules toward the transient-encounter surface. In the post-encounter stage, molecular dynamics are typically better suited to capture conformational flexibility, intra-molecular dynamics (such as binding site gating), and solvation phenomena involved in molecular recognition.

### 2.2 Single Binding Events

A single ligand/protein binding event serves as the minimal motif in multi-component cellular signal transduction pathways. To simplify our discussion of these binding events, we adopt a common approach of dividing the process into two phases [5, 6]: 1) The ‘transient encounter’ during which two species approach from a distance to assume loosely bound complexes (Figure 1b, green), and 2) The post-encounter where transient encounter complexes settle into their final bound states (Figure 1b, pink). The transient encounter is a non-local process governed predominantly by long-range diffusion and electrostatics. In contrast, the ‘post-encounter’ is a local process dominated by short-range forces, conformational changes, and desolvation. A microscopic view of the interplay between these long-range and short-range interactions is essential for understanding the molecular mechanisms of binding events and how their behavior is shaped by cellular environments, such as the cell cytoplasm [7]. For the special case of rapid reactions, this entails characterizing both the rate of formation for a transient encounter complex and its transition to the final bound state.

#### 2.2.1 The Transient Encounter

The transient encounter encompasses the rapid, but generally intermediate, assembly of substrates with target enzymes [8, 9] ; we provide an example of this for nucleotide binding to the ectonucleotidase CD39 in Figure 1b. This assembly begins with weak, long-range electrostatic interactions that can bias random molecular collisions of the substrate with its target. Early descriptions of this process assumed a spherical target and a point substrate in solution that permitted analytical solutions for the corresponding association rate [10]. However, many biological binding events are not well-described by the spherical paradigm [11]. For instance, the diffusion of quinone within the mitochondrial membrane, which shuttles electrons between respiratory complexes [12, 13] (Figure 2a), is better described as a 2D binding process; fortunately, this system is also amenable to a simple model of 2D diffusion that is also analytically solvable [14]. Additional examples that deviate from the spherical paradigm but permit simple quantitative models include the association of tethered substrates with targets [15], such as folded LIM domains within disordered proteins binding to targets on muscle filament proteins [16](Figure 2b), or the association of a particle undergoing 3D free diffusion before colliding with a 2D surface [17], such as for an extracellular ligand activating a membrane-bound GPCR [18](Figure 2c). Binding events that can be described as one-dimensional, diffusion-controlled processes also arise in nature, [19], such as the diffusional sliding of a transcription factor along DNA to locate a promoter sequence, or binding events involving an ensemble of intermediate states [20]. Although simplified analytic or numerical models can be applied to many binding events found in biology, when atomistic detail or nontrivial interaction potentials are important, such approaches are typically infeasible. Here, Brownian dynamics simulations have emerged as a useful tool for capturing atomistic detail and nontrivial substrate/target interactions shaping substrate/target binding kinetics.

**Figure 2:**
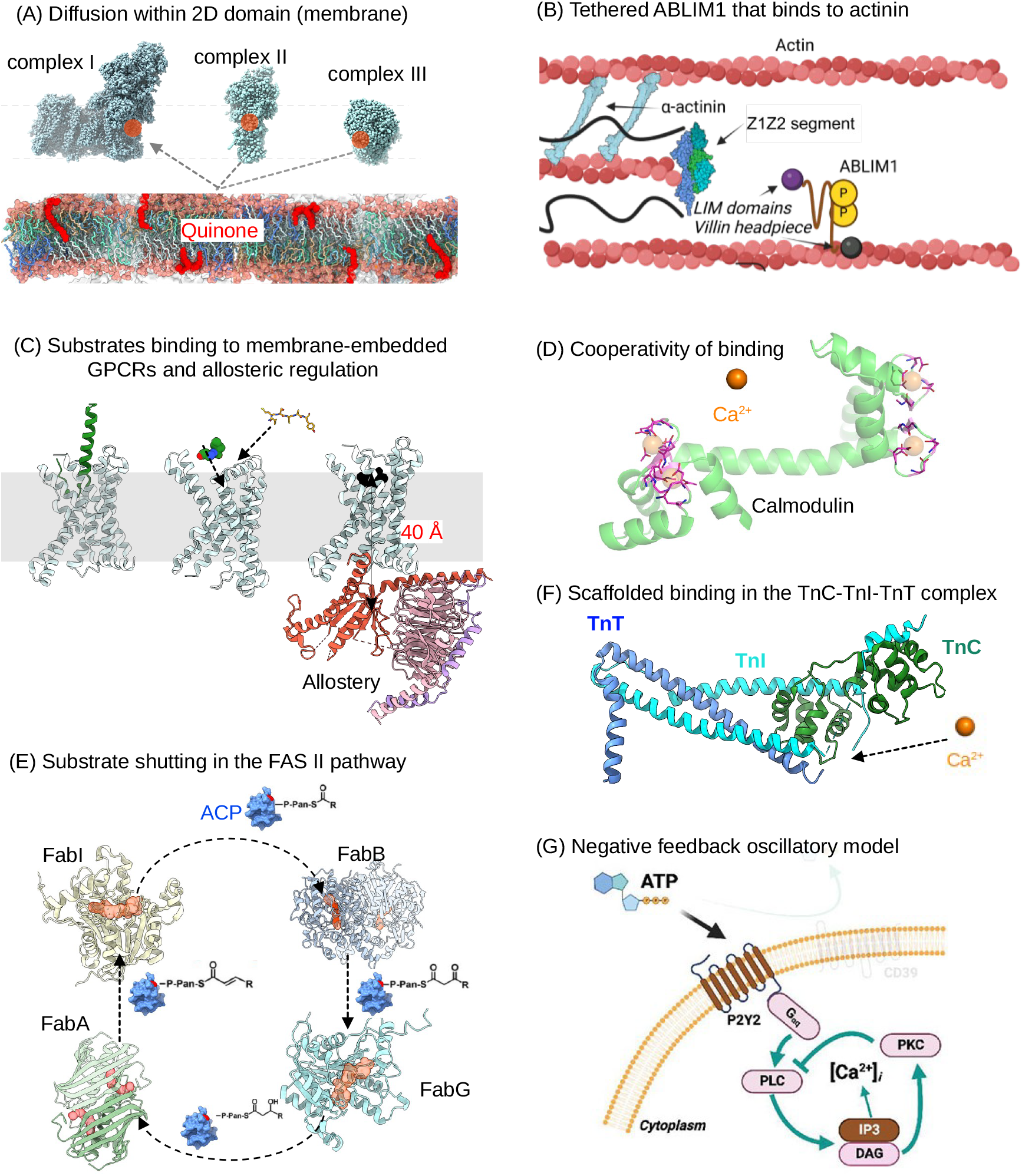
Complex environments of binding events, scales, and feedback. Although conceptual binding events often involve just two partners, these events are much more complex in biological systems due to their unique environments. (a) Substrate diffusion in an organelle membrane, such as quinone molecules diffusing within the mitochondria inner membrane to serve as charge carriers for electron transfer among respiratory complexes[12, 13]. (b) ‘Tethered’ actin-binding LIM domain protein (ABLIM) (@2025 Sun et al. originally published in [16]). (c) Extra-cellular agonists binding to cell-membrane bound GPCR receptors, which promote the allosteric binding of intracellular proteins. (d) Cooperative binding between Ca^2+^ ions and calmodulin, a protein that contains four Ca^2+^ binding EF-hands [21]. (e) Substrate shuttling between enzymes, as shown for the FAS II pathway for which the ACP protein shuttles intermediates between enzymes via diffusion [22]. (f) Scaffolded binding of TnI-TnC-TnI that is regulated by Ca^2+^ ions [23]. (g) Oscillatory, negative feedback as shown for phospholipase C (PLC), protein kinase C (PKC), and phosphatidylinositol 3,4,5-trisphosphate (PIP_3_) cycle (adapted from [24]).

#### 2.2.2 Post-encounter Binding

The post-encounter event involves the transition of transiently-bound complexes into the final, higher affinity and functional poses (Figure 1b). Intermediates from the transient-encounter complex present diverse poses differing in their propensity to form the bound state. The consolidation of loosely-bound poses with their final binding mode is driven by free energy gains from structural rearrangements that optimize chemical complementarity between a ligand and its associated protein [25], in addition to potential contributions from desolvation of hydrated reactants and even quantum chemical effects inclusive of polarization and covalent binding [26]. Short-range forces governing association can also impact the conformation ensemble by guiding conformational changes and electrostatic interactions that shift the system to its final, high-affinity bound state. It is also understood that even the reorganization of the solvent near binding interfaces can affect both the thermodynamics and kinetics of binding/unbinding [27–29]. The collective contributions of diverse mechanisms governing the shift of a transiently-bound substrate toward its post-encounter state can dramatically influence binding rates [30, 31]. Since the focus of this review is on forming the transient-encounter complex via diffusion, we refer the reader to robust discussions on this topic elsewhere [30, 32–34].

### 2.3 Multiple Binding Events

Binding events in biological systems can entail multiple reactants, binding sites, or feedback mechanisms. Allostery and cooperativity (Figure 2d-e) are two important factors for characterizing processes involving multiple reactants or binding sites. Allostery generally describes the binding of substrate to a distinct site on a target that in turns affects the binding of a different molecule elsewhere [35]. Such allosteric behavior is commonplace among GPCRs. For these transmembrane receptors, an agonist’s binding to their extracellular domains ultimately drives G-protein binding on the intracellular side at distances nearly 40 Å in magnitude [36]. Cooperativity in turn refers to when the binding of one substrate molecule either facilitates or disfavors the binding of a subsequent substrate of the same type [37]. A prototypical example of cooperative binding can be found in the binding of Ca^2+^ to calmodulin, of which the latter is a protein comprising four Ca^2+^ binding sites called EF-hands [21] (Figure 2d), each with different tendencies to bind the cation.

Another consideration in processes involving multiple binding events includes substrate channeling (Figure 2e), whereby intermediates produced by one enzyme are guided for further processing by a subsequent enzyme, often through their spatial co-localization [22]. For instance, in vitro reconstitution experiments have demonstrated how transient complexes, such as the human 8-oxoguanine glycosylase 1 (OGG1) and human AP endonuclease 1 (APE1) complex, align consecutive steps in base excision repair [38]. Related to this is scaffolding (Figure 2f), whereby substrates are colocalized to a protein or other biomolecule to favor a reaction or pathway through proximity and often through allosteric means. Here, the association of Ca^2+^-bound TnC with a segment of TnI is facilitated by their joint tethering to actin filaments [23], is a canonical example. Signaling cascades as a whole are complex and can be strongly coupled. One such example is the coupled conversion of ATP into AMP and ultimately adenosine, which factors into cancer immunosuppression [4] (Figure 1a). Many of these cascades are subject to feedback control, whereby reaction products can enhance or suppress enzymes involved in their synthesis, as exemplified by the phospholipase C (PLC), protein kinase C (PKC), and phosphatidylinositol 3,4,5-trisphosphate (PIP_3_) pathways in Figure 2g [39].

### 2.4 Kinetic Dysregulation in Disease

The precise biological control of substrate binding kinetics is of fundamental importance in human health and disease. In fact, many diseases can be caused by, or exacerbated by, enzymes whose reaction rates are altered relative to their physiological levels. For instance, dysfunctional enzymes important to development can disrupt the precise timing of cell division, maturation, and differentiation, leading to congenital defects and pediatric cancers [40, 41]. Similarly, a mismatch in reaction rates can be at the root of metabolic diseases. A classic example is phenylketonuria (PKU), where mutations in the phenylalanine hydroxylase (PAH) enzyme lead to impaired catalytic [42] conversion of phenylalanine to tyrosine, leading to the accumulation of phenylalanine and neurotoxic byproducts. As another example, the precise timing of calcium binding to cardiac myofilaments is critical for forceful and energetically-efficient muscle contraction. Certain mutations, such as the A8V mutation in cardiac troponin C (TnC), alter kinetics by increasing TnC’s calcium binding sensitivity [43]. This enhanced binding impairs cardiac muscle’s ability to relax, which ultimately leads to diastolic dysfunction. Similarly, impaired binding kinetics of proteins involved in neurotransmission can contribute to neurological diseases [44]. Similarly, in pharmacological contexts, drug design and delivery can be ineffective if the substrates do not engage their targets for sufficient lengths of time [45].

### 2.5 The Role of Modeling with Brownian Dynamics

Given the complexity and importance of the substrate/target binding events, computational modeling is an indispensable tool for systematically evaluating the many physicochemical factors that can influence such processes, synthesizing experimental data probing their effects, and generating novel, mechanistic hypotheses. In this review, we position Brownian dynamics as uniquely well-suited for modeling complex, biomolecular processes and their kinetics in molecular detail. Toward this end, we formalize in Section 3 the fundamental physical relationships underlying the binding scenarios outlined above.

## 3 The Molecular Physics of Binding Events

This section delves into the fundamental physical principles modeled by Brownian dynamics that govern ligand-protein interactions. These principles provide a quantitative basis for our discussion of computational approaches for simulating binding events in Section 5.1. We also briefly introduce complementary experimental methods for evaluating ligand binding thermodynamics and kinetics.

### 3.1 Mass Action and Free Energy of Binding (Δ*G*)

We begin with the thermodynamic foundations of the equilibrium between bound and unbound species. We start with the principle of mass action and free energy of binding (Δ*G*). The fundamental relationship between substrates and products in a reversible reaction is given by:

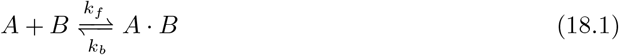

Here, B represents an apo enzyme that forms a holoenzyme complex (A · B) upon binding substrate A, with *k*_*f*_ and *k*_*b*_ representing the forward (association) and backward (dissociation) rate constants, respectively (see Figure 3a). At equilibrium, the dissociation constant (*K*_*D*_) is defined as:

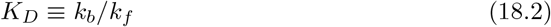

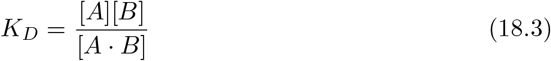

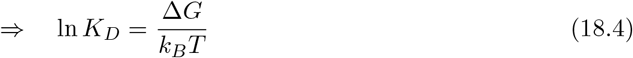

**Figure 3:**
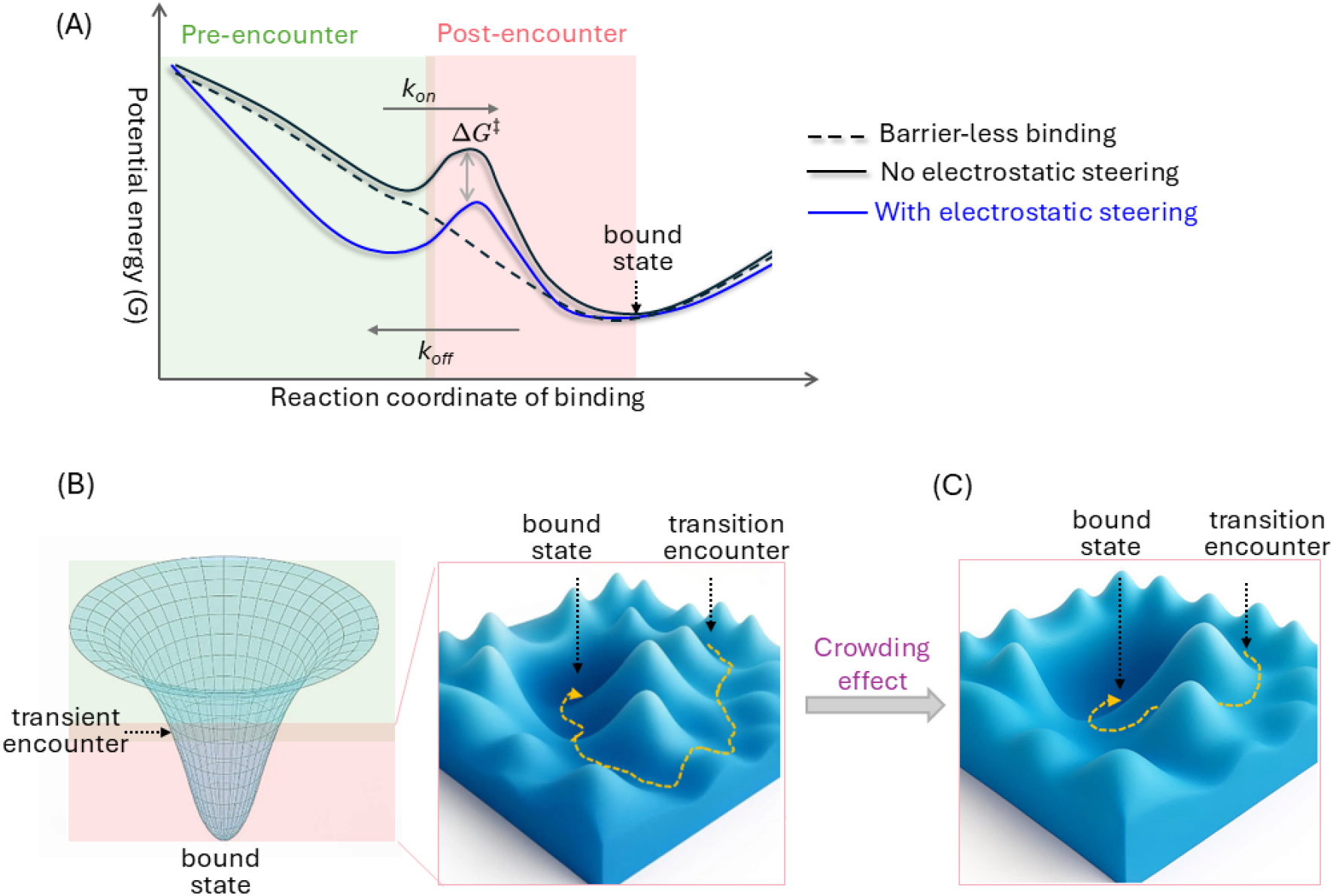
Energy landscape view of biomolecular binding events. (a) The overall potential energy of a complex decreases throughout the binding process, from the pre-to post-encounter stages, until the bound state is reached. The thermodynamic favorability of forming the bound state from isolated components determines the event’s spontaneity. However, the binding rates (*k*_*on*_ and *k*_*off*_) are determined by the heights of the energy barriers between the bound and unbound state, rather than the thermodynamic stability of the final complex. During the pre-encounter stage, biomolecules are often guided by long-range electrostatic interactions that can accelerate encounter complex formation. Here, the association rate (*k*_*on*_) dictates the rate of a substrate’s arrival in the local energy minimum representing the encounter complex, while the intrinsic rate (*k*_*int*_) dictates the subsequent crossing of the main activation barrier to reach the final bound state. (b) Binding follows a funneled energy landscape, with the native bound state at the global minimum. The intrinsic flexibility of biomolecules is associated with a rough binding funnel, featuring numerous metastable intermediate states between the initial encounter and the final bound state. In a realistic cellular environment, macromolecular crowding can be conceptualized as altering the distribution and stability of these intermediate states, thereby modulating the overall binding kinetics.

The last line reflects the relationship between the standard Gibbs free energy of binding (Δ*G*) and the dissociation constant.

We next outline key thermodynamic contributions to the Δ*G* associated with Eq. 18.1. As shown in Eq. 18.2, the binding equilibrium constant (*K*_*D*_) determines the equilibrium distribution between the bound and unbound states of a reaction; its values commonly range from high affinity picomolar (pM) constants to low affinity millimolar (mM) values. While Δ*G* indicates the spontaneity of a reaction, the *k*_*f*_ and *k*_*b*_ rates determine its feasibility on biological timescales (e.g., whether the system is in rapid equilibrium or presents long-lived, non-equilibrium configurations). As a state function, Δ*G* depends only on the initial and final states, but not the specific reaction pathway. This thermodynamic quantity reflects the sum of all energetic interactions among the reactants, products, and the surrounding solvent [46]. The change in free energy upon binding, Δ*G* = Δ*H* − *T*Δ*S*, can be decomposed into its enthalpic (Δ*H*) and entropic (Δ*S*) components. Partition functions provide a quantitative basis for these terms [46], which enumerate interactions not only between the binding partners (e.g., enzyme and substrate) but also with the solvent [47]. Solvent contributions are key considerations in describing binding thermodynamics. The hydrophobic effect for instance drives the assembly of nonpolar surfaces to minimize energetically unfavorable interactions with water. Underlying this phenomenon is the release of water molecules bound to the reactants or target binding site that can contribute substantial gains in entropy upon binding [48]. The complexity of these interactions motivates the use of simulation techniques like molecular dynamics (MD) simulations, which can extensively sample ensembles of thermodynamically-accessible states and their associated energies. It is also important to note that the local concentrations of the ligand [A] and target [B] used in Eq. 18.1 may considerably deviate from their bulk values, which can locally drive product formation relative to the surrounding medium. This behavior arises in contexts such as substrate channeling or tethering.

### 3.2 Association and Dissociation

For systems out of equilibrium, it can be meaningful to track the time-dependent concentration changes of the reactants and products. Given the process described in Eq. 18.1, we can write the following reaction scheme:

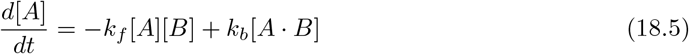

where *k*_*b*_ represents the dissociation rate, and *k*_*f*_ is the forward rate constant. When considering a system of diffusible substrates, the forward rate is commonly decomposed into an initial rate of transient encounter (*k*_*on*_) and a rate for forming the final complex (*k*_*int*_) (see Figure 3top). The overall forward rate constant (*k*_*f*_) is then given by [49, 50]:

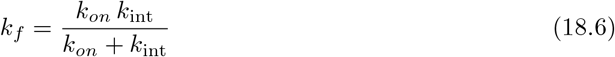

where *k*_*on*_ and *k*_*int*_ represents the diffusional encounter and intrinsic reaction rates, respectively.

The overall rate of a reaction is determined by its slowest step, which leads to two distinct kinetic regimes. For first is the reaction-limited regime, for which the intrinsic chemical or conformational step is much slower than the diffusional encounter rate (*k*_*on*_ >> *k*_*int*_) and hence the overall rate is dominated by the intrinsic rate (*k*_*f*_ → *k*_*int*_) per Eq. 18.6. The second is the diffusion-limited regime, for which the intrinsic rate is much faster than the association rate (*k*_*int*_ >> *k*_*on*_); here, the overall rate approaches the diffusional encounter rate (*k*_*f*_ → *k*_*on*_). A formal definition is discussed in detail elsewhere [50].

#### 3.2.1 Factors Governing *k*_*on*_: Diffusion-limited

Many biological reactions are limited by the rate of diffusional search of a substrate for its target. For these cases, *k*_*on*_ implicitly reflects the solute’s exploration of a potential energy landscape to form an encounter complex (see Figure 3). In the diffusion-limited regime, these rates typically range from 10^5^ to 10^9^ M^−1^s^−1^ [51], whereas in reaction-limited processes the intrinsic catalytic rates (*k*_*int*_) can vary from milliseconds to seconds [52].

Physical factors across a hierarchy of spatial scales, from long-range diffusion to local interactions, collectively shape substrate/target association. At large distances, the diffusion coefficient (*D*) helps determine the rate of substrate/target assembly. This coefficient reflects the contribution of random collisions with the solvent that both propel and dampen molecular diffusion through friction. Hydrodynamic interactions arising from the displacement of solvent by large solutes like proteins can also impact association rates relative to small molecules [53].. In addition, the dimensionality of the problem domain is also an important factor for determining collision timescales, as searches constrained to lower (1D or 2D) dimensions often proceed more efficiently than in 3D [54].

In parallel, long-range mean-field electrostatic interactions have the capacity to steer molecules toward assembly, often increasing rates by orders of magnitude [51] relative to their basal rates (without electrostatic contributions). The barnase-barstar interaction [55] is a classic example of this phenomenon, where electrostatic steering drives binding rates to 10^8^–10^9^ M^−1^s^−1^ [55]. Similar electrostatic acceleration is observed in intrinsically disordered proteins, such as the CaMBR (Calmodulin-binding region) of calcineurin binding to calmodulin, even where conformational gating of the binding interface can limit access to the final binding configuration [56]. Following the initial diffusion encounter, the system transitions toward a more tightly associated state where short-range forces become dominant (Figure 4, bottom).

**Figure 4:**
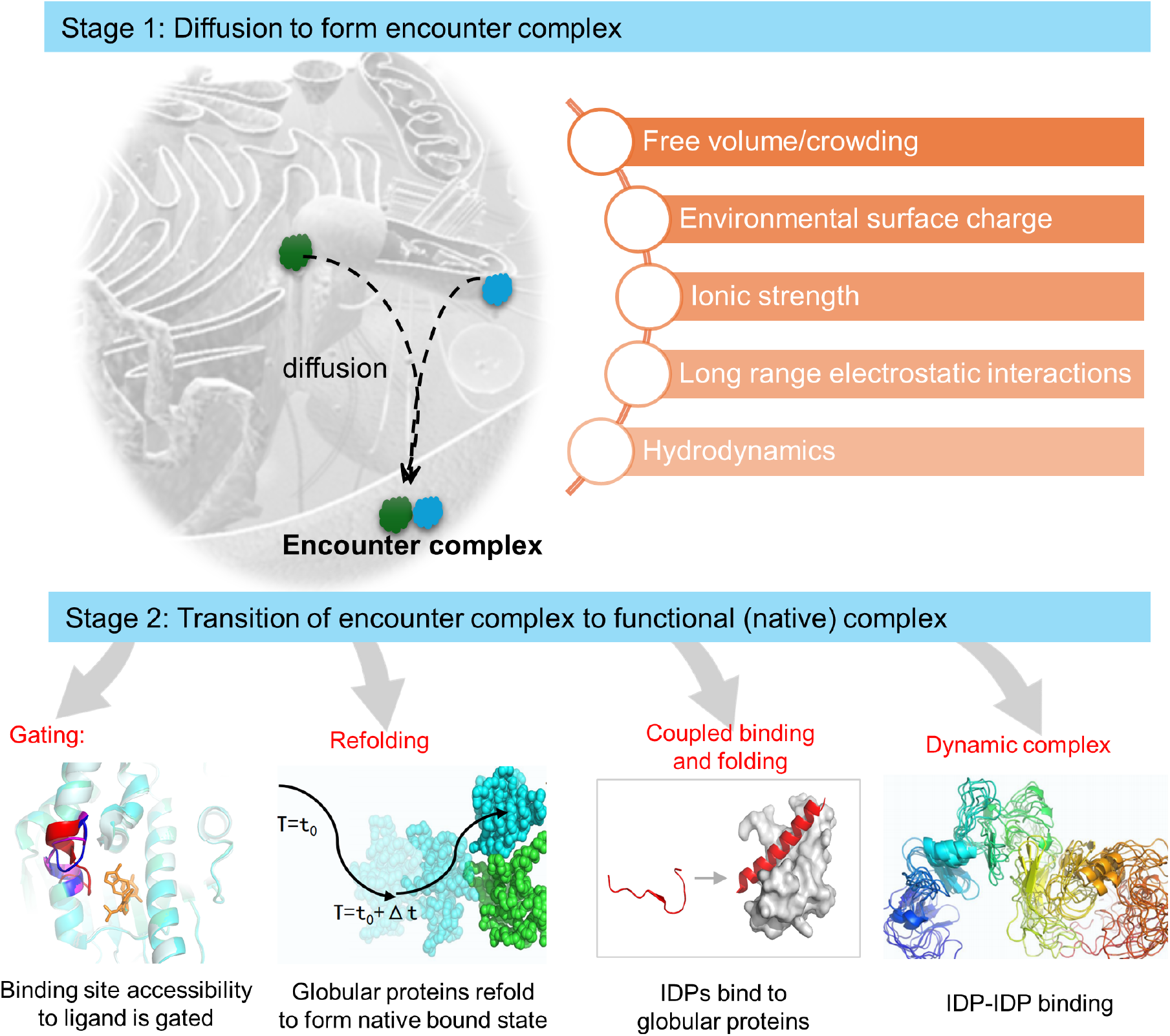
Two-stage ligand/protein binding. Biomolecule binding is frequently described as a two-stage process (Figure 3). The first stage involves the formation of an encounter complex through diffusion, often guided by long-range electrostatic interactions, with typical rates of 10^5^ to 10^6^ M^−1^s^−1^. The second stage entails the transition from this encounter complex to the final binding pose, a process whose rate is determined by an assortment of physicochemical factors. Some binding events are shaped by conformation gating, as exemplified by inhibitor-HSP90 binding [57]. In this system, the binding site’s accessibility to ligands is controlled by the time-dependent conformation shifts of a ‘lid’ that transitions between a loop and an *α*-helix [57]. The refolding of a transient complex into the final bound state is depicted here for p53-ASSP protein-protein binding, where simulations revealed encounter complexes that exhibited structural fluctuations relative to their initial contact poses as evidence of refolding [58]. Coupled binding and folding is a paradigm of substrate/target recognition mechanisms between intrinsically disordered proteins (IDPs) and globular target, whereby an IDP can adopt secondary structure upon forming the final bound state [59]. Binding may also take the form of fuzzy complexes, typically seen in IDP-IDP binding, such as the H1–ProT*α* complex [60]; for such events, changes in conformational entropy play a key role in determining the binding free energy. Modern structural biology and AI offer compelling possibilities for probing transient states involved in these binding processes. For instance, preliminary models of encounter geometries can be generated from programs such as AlphaFold-Multimer [61], and then refined with BD simulations. In parallel, generative AI models could be trained with emergent data from time-resolved X-ray or NMR to capture transient structural states.

#### 3.2.2 Factors Governing Association Kinetics

Reaction-limited systems are determined by the timing of a substrate’s chemical conversion into a final product. The rates of which are shaped by events occurring between the transient encounter of a substrate with its target and the substrate’s eventual recognition at the binding site. [20] These contributions are reflected in the *k*_*int*_ term of Eq. 18.6 that shapes the overall forward rate. An important consideration for post-encounter events is the energetic cost of conformational changes a protein and its substrate each undergo to reach the final bound state [62]. One such contribution is the desolvation process stemming from the water molecules squeezed out from the interfaced formed between substrate/target molecular surfaces that stabilize a bound complex. Configurational entropy increases as hydrating waters are released from the active site, which can compensate for entropic losses associated with confining both the ligand and target to their bound conformations [63]. In some cases, complex dewetting behavior manifests between solvated molecules, comprising transient transitions between wet and dry states of interacting molecular interfaces; these transitions can introduce long-lived correlations in otherwise ‘memoryless’ random binding events [64]. In a similar vein, ‘dangling’ hydrogen bonds at hydrophobic solute–water interfaces [65] have been suggested to lubricate conformational changes during binding.

#### 3.2.3 Factors Governing *k*_*off*_

While the association rate often decides the timing of the initial binding event, the dissociation rate, or *k*_*off*_, determines the persistence of the bound state and is implicated as a critical factor for biological function.[45]. In fact, recent findings suggest in vivo drug efficacy correlates strongly with the dissociation rate (*k*_*off*_) [66]. For a binding event without a subsequent chemical reaction, the association (*k*_*on*_) and dissociation (*k*_*off*_) rates fully determine both the kinetics and the thermodynamic equilibrium of the system. In the event that a binding event is diffusion-limited, variations in intrinsic dissociation rates can disproportionately influence binding free energies. Dissociation rates are determined by the most prominent energy barriers separating the bound and unbound states. These energy barriers arise from thermodynamic contributions ranging from the enthalpic cost of breaking strong, close-range nonbonded interactions to the entropic penalty of solvent reorganization and changes in substrate/target mobility during dissociation.

## 4 Experimental Approaches for Characterizing Thermodynamics and Kinetics of Binding

### 4.1 Assays for Slow Reactions or Steady-state Conditions

A number of experimental techniques are used to measure binding thermodynamics and kinetics, which can be used to inform and validate theoretical models. While a comprehensive survey of equilibrium and kinetic assays is beyond the scope of this review, we highlight several commonly-used methods, but defer the reader to excellent reviews on these topics and best practices [67, 68]

### 4.2 Isothermal Titration Calorimetry (ITC)

Among the core experimental techniques used to measure binding thermodynamics and determine thermodynamic quantities like Δ*G* and Δ*H*, isothermal titration calorimetry (ITC) is a gold standard [69]. ITC directly measures the heat released or absorbed upon binding [69], through incremental additions of ligand. Hence, ITC is an ideal tool for capturing the thermodynamics of binding resulting from the formation of the post-encounter complex from isolated components. ITC can also be applied to monitor allostery and cooperativity that may contribute to these binding mechanisms. In these complex cases, fitting the ITC data requires extra caution, as different binding models can produce equally good fits to the same dataset [70]. For such systems, variable-c ITC has been applied, which extends conventional ITC by titrating ligands across a range of protein concentrations. This approach was utilized by Bonin et al. [71], for example, to quantify cooperative binding of dUMP to thymidylate synthase (TS).

In addition to ITC, fluorescence-based techniques discussed later in Section 4.3.2.1 can also be used to infer cooperative and allosteric binding mechanisms [72, 73]. For instance, by utilizing the intrinsic tryptophan fluorescence of the glucokinase regulatory protein (GKRP), Zelent et al. revealed cooperative behavior exhibited by several allosteric ligands [74].

Classical experimental techniques, such as colorimetric assays and Western blotting, are appropriate for measuring slow reaction-limited processes. Colorimetric assays monitor color changes as a reaction progresses and remain a foundational method for high-throughput screening [75]. Early quantitative measurements of biochemical processes often focused on enzyme-catalyzed reactions [76]. Much of this work was performed using spectrophotometers that measure absorbance changes as a reaction proceeds. These methods are limited to tracking relatively slow processes, making them most appropriate for characterizing steady-state or reaction-limited kinetics [76]. Western blotting is a complementary technique rooted in measuring protein concentrations by antibodies. Western blots lend themselves to kinetic studies when done as time dependent assays, e.g. by monitoring the change in a protein’s concentration after a reagent is applied [77]; unfortunately, these assays are generally labor-intensive, low throughput, and can be very qualitative in accuracy.

Kinetic ITC can also be used to estimate rate constants for enzyme-inhibitor interactions [78], but are also limited to slow reactions.

### 4.3 Techniques for Rapid Kinetics and Conformational Dynamics

#### 4.3.1 Fluid-phase and Surface-based Kinetics

##### 4.3.1.1 Stopped-flow for Rapid Kinetics

Estimation of rapid, diffusion-limited rates requires specialized experimental techniques capable of capturing events on millisecond and shorter timescales. Among these, stopped-flow spectroscopy is a gold standard for rapid binding kinetics, with a key advantage being its short dead time, e.g. the period of time before reliable measurements can be made, is on the order of milliseconds. This rapid dead time makes it suitable for measuring fast *k*_*on*_values [79], such as those associated with diffusion-limited reactions [80]. One of the earliest stopped-flow apparatuses was developed by Chance and Gibson [81] in the late 1960s. Over the following decades, the technique has been used in a variety of applications, including capturing fast pre-steady-state kinetics and resolving transient intermediates [82]. In addition to stopped-flow, Surface Plasmon Resonance (SPR) is often used for kinetics (*k*_*on*_, *k*_*off*_), including those influenced by diffusion [83], which we discuss below.

Label-free techniques like SPR and BLI are widely used for the real-time study of biomolecular interactions and the estimation of kinetic parameters. These methods immobilize a binding partner on a sensor surface while the other partner is flowed over it, during which binding events can be monitored without the need for fluorescent or radioactive labels. SPR works by detecting a change in refractive index at a metal-film interface [84]; in the dissociation phase, where the bound ligand dissociates from the immobilized chip, the decayed SPR signal yields the corresponding dissociation rate [85].In contrast, BLI measures the interference pattern of light reflected from two surfaces [86], with the shift being proportional to the number of bound molecules. BLI is less sensitive to bulk refractive index changes compared to SPR [86]. For both approaches, data are plotted to give the binding response versus, which can highlight the association, saturation, and dissociation phases of the interaction [86]. Accordingly, kinetic parameters such as *k*_*on*_, *k*_*off*_, and the equilibrium dissociation constant (*K*_*D*_) are extracted by fitting reaction models to these phases.

#### 4.3.2 Structural Dynamics and Label-Based Approaches

##### 4.3.2.1 Fluorescence-based Methods

The discovery of Green Fluorescent Protein (GFP) has been transformative for cell biology, enabling the observation of dynamic molecular processes in vivo [87]. This discovery enabled many new techniques that could simultaneously follow proteins in subcellular detail and in real time. Methods based on labeling of proteins with fluorescent probes, including genetically-encoded domains based on GFPs, are now commonplace for in vivo studies.

Among these, Förster Resonance Energy Transfer (FRET) is a popular technique based on introducing a fluorescent donor probe and acceptor probe to a protein substrate and target, respectively, via genetic or chemical means. If the distance between the paired probes varies over a 1 to 10 nm range, the efficiency of fluorescence energy transfer from the donor to the acceptor can be measured via the acceptor’s emission spectrum [88]. Hence, FRET enables time-dependent measurements of protein association and dissociation [89], as well as conformational changes induced by PTMs or substrate binding to form functional complexes.

FRET has also been adopted to measure drug-target binding kinetics, typically through competitive assays using unlabeled drugs that displace a fluorescent probe bound to the target [90]. As an example, high-throughput kinetic probe competition assays based on FRET have been used to survey drug kinetics across broad classes of protein kinases [91]. In addition, single-molecule fluorescence resonance energy transfer (smFRET) is particularly well suited for probing biomolecular binding events, given its high sensitivity to probe distances [73]. Applications of smFRET include studies revealing conformation changes that occur upon association [92], such as complexin binding to the SNARE complex [93].

Fluorescent probe labeling techniques are also commonly used to measure biomolecule diffusion in heterogeneous environments, with green fluorescent protein (GFP) being widely used for proteins. For example, GFP-labeling of G3BP1 enabled studies of liquid–liquid phase separation (LLPS) in cells, including the characterization of its diffusion behavior both before phase separation and within the resulting condensates [94]. This GFP labeling strategy for studying protein dynamics within condensates has been broadly used, with applications to bacterial cells [95] and nuclei where transcription factors can form condensates [96]. For smaller molecules, fluorescent dyes are often used to label metabolites from enzymatic process, such as ATP, to characterize their diffusion in cellular environments. For instance, by labeling ATP with a fluorescent moiety, Vendelin and colleagues showed that ATP diffusion is hindered by the compact filamentous sarcomeres of rat cardiomyocytes [97]. Fluorescent dyes can also be used into tandem with genetically-encoded fluorophores, such as for measuring cargo-transport in the cytoplasm. The aforementioned G3BP1 LLPS study elegantly applied this tandem strategy, by using quantum dots that were co-loaded with GFP-labeled G3BP1, enabling simultaneous characterization of crowder diffusion (via the quantum dots) and LLPS formation dynamics (via the GFP label) [94].

#### 4.3.3 Nuclear Magnetic Resonance (NMR) Spectroscopy

As a complementary approach, NMR spectroscopy provides a unique window into atomic-level details of kinetic processes across a broad range of timescales. NMR explicitly probes motion and exchange processes spanning picosecond to second timescales [98]. This affords measurements of molecular structure and conformation dynamics at atomic resolution [99]. Examples include following bond rotations on the ps–ns scale via relaxation measurements and the application of relaxation dispersion techniques to characterize conformation exchange on the *µ*s–ms timescale [98]. Two-dimensional exchange spectroscopy (2D EXSY) experiments capture slower exchange rates, such as dissociation processes that occur at the second timescale [98].

NMR is almost unrivaled at dissecting complex kinetic mechanisms in atomistic detail, including conformational gating [98]. For example, the KIX domain was studied via NMR in the formation of binary complexes with c-Myb, as well as ternary complexes with c-Myb and MLL. This information provided a structural basis of positive cooperative binding of these ligands to KIX [100]. NMR may also be complemented with Hydrogen-Deuterium Exchange Mass Spectrometry (HDX-MS) to identify allosteric conformational changes upon binding [101]. Binding rates can also be assessed via NMR by tracking changes in residue-specific relaxation rates before and after ligand binding [102]. A key advantage of these NMR-based methods is their ability to characterize weak binding events that typically exhibit fast dissociation rates [102]. Lastly, NMR as well as smFRET can resolve protein conformational states and their dynamics upon ligand binding, towards identifying induced-fit and conformational-selection pathways [103, 104].

### 4.4 Structure determination

Atomistic-level structural data are often instrumental for interpreting data from kinetics experiments and devising new molecular mechanisms. Classically, X-ray crystallography has been the standard for structural interpretation of binding data, such as revealing the cooperative binding between transcription factors and DNA [105]. Other techniques have more recently emerged as complements to X-ray crystallography, including NMR, cryoelectron microscopy, and ab initio structure prediction via tools like AlphaFold and RoseTTAFold [61, 106]. Recent reviews provide a broad overview of the current state of the art in this rapidly evolving area [107, 108].

## 5 Modeling Diffusion-limited Kinetics with Brownian Dynamics

MD and BD are the most common simulation approaches for modeling substrate/target binding using detailed molecular structures. Briefly, MD provides all-atom resolution of binding events, but is generally too computationally intensive to model those involving diffusional encounters; on the other end, analytic mathematical models lend themselves to diffusion-limited reaction modeling, but lack explicit molecular details. BD simulations serve as an intermediate to the two extremes, by treating the solvent as a continuum (implicit solvent), in contrast to explicit solvent models used for all-atom simulations. This efficiency makes it practical to simulate thousands of trajectories (Figure 1b), enabling the robust statistical sampling required to study kinetic processes. BD also provides a flexible framework for accounting for a variety of physicochemical factors that influence interactions within and between biomolecules, from solvent contributions to electrostatic potentials. Beyond kinetics, its utility extends into equilibrium properties as well, such as predicting ensembles of flexible systems like intrinsically disordered proteins. We therefore posit that Brownian dynamics is an ideal framework for balancing the detailed resolution of molecular dynamics with the computational efficiency of continuum models to examine binding processes such as those provided in Figure 1. To demonstrate the utility of BD, it is helpful to first establish its fundamental physical basis, and then compare it to other simulation techniques used for probing transient encounters and coupled binding events. Therefore, in this section we discuss a variety of paradigms in the modeling of kinetics and where BD is appropriate.

### 5.1 Fundamentals of the Brownian Method

The mathematical foundation of BD lies in the Langevin equation, which describes the stochastic motion of a solute subject to friction and an external potential:

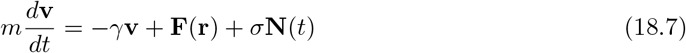

Here, the inertial term 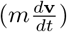 is balanced by the frictional drag arising from the solute’s interaction with the solvent (− *γ***v**), forces from molecular interactions (**F**(**r**)), and a random force (amplitude *σ*) from solvent collisions (**N**(*t*)). The friction term can be estimated from the Stokes-Einstein equation and can include hydrodynamic effects when appropriate. The force term, **F**(**r**), can encompass long-range interactions such as an electrostatic potential or local interactions arising from van der Waals forces. Oftentimes these long-range interactions are implicitly represented by a potential of mean force (PMF). A PMF is an effective free energy landscape that averages out degrees of freedom orthogonal to the reaction coordinate, including the solvent degrees of freedom [46].

The fundamental assumption of BD is that for a particle in a viscous solvent, the system is in the high-friction limit, for which the frictional drag term dominates the inertial term. This allows the inertial term to be considered negligible 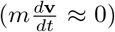, which yields the over-damped Langevin equation frequently used for BD simulations:

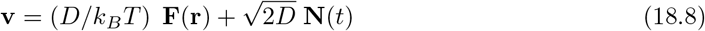

where we have used the Einstein relation:

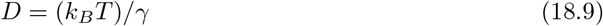

and *D* = *σ*^2^/(2*γ*^2^) to express the equation in terms of the diffusion coefficient. The friction (and thereby diffusion) coefficient fundamentally arises from the fluctuation-dissipation theorem (Eq. 18.10) that relates the random kicks the solvent imparts on the solute to the macroscopic frictional drag it experiences.

In aqueous media, force fluctuations arising from the random movements of water molecules transfer momentum to the surroundings, resulting in energy dissipation via friction. This behavior can be linked to the diffusion coefficient by the fluctuation-dissipation theorem, such as by using the Green-Kubo relationship. The Green-Kubo relation [109] is a formalism that provides the friction coefficient from the time integral of the random force autocorrelation function:

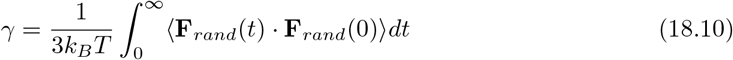

where **F**_*rand*_(*t*) corresponds to the instantaneous random force, parameterized as *σ***N**(*t*) in Eq. 18.7 [109]. The diffusion coefficient, *D*, can then be determined by the Einstein-Smoluchowski relation (Eq. 18.9) from the friction coefficient. Later in Section 5.2 we will show how *k*_*on*_ is positively correlated with increasing *D*. Beyond dissipation through random solvent collisions, the translational and rotational diffusion of larger solutes, such as proteins, can be impacted by significant intramolecular hydrodynamic interactions. To characterize these, it can be useful to consider the autocorrelation functions of a solute’s position or velocity are used in lieu of Eq. 18.10 [110, 111]:

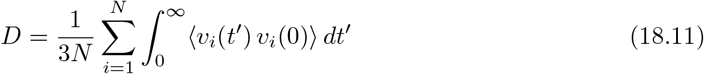

where *v*_*i*_ signifies the velocity of particle i at time *t*^′^ or 0 and N is the number of particles. This provides a basis for extended formulations of Eq. 18.8 such as those from Ermak and McCammon for capturing translational and rotational degrees of freedom inclusive of hydrodynamic interactions [112]; alternatively, approximations for hydrodynamic interactions may suffice, such as those due to Rotne-Prager [113].

### 5.2 Transient Encounter: BD for Homogeneous Media

#### 5.2.1 Modeling Rigid Solutes

Early treatments of enzyme kinetics involved model systems (often spherical), for which association rates captured a substrate’s diffusion to a reactive site on the enzyme surface. We overview important kinetic frameworks arising from this simple system, beginning with the Smoluchowski model; this model describes diffusion toward a spherical enzyme with an absorbing surface as the active site. From there, we discuss Kramers’ theory to address the concept of an energy barrier omitted from early Smoluchowski descriptions. We additionally present more modern and rigorous ways of analyzing reaction kinetics, including committor analysis, Markov models, and survival probability.

##### 5.2.1.1 Kinetic Models

###### 5.2.1.1.1 The Spherical Model

The study of diffusion-limited reactions often begins with the simplest possible case: a spherical target. This perspective provides an easy to compute framework for understanding diffusion-limited reactions, from which refinements can be developed to account for more realistic protein shapes, electrostatic interactions, and reactive surface patches. We start with the Debye-Smoluchowski model [114], which describes a particle diffusing toward and reacting with a spherical enzyme governed by a centrosymmetric interaction potential. This is usually framed (in steady state) as a Laplace problem of the form:

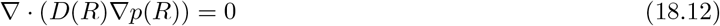

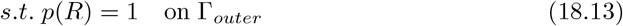

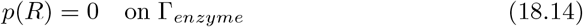

These equations describe the relationship between a particle reactant’s concentration *p*(*R*) and its diffusion coefficient *D*(*R*) at a given radius *R*. The system is contained within two boundaries: Γ_*outer*_ (the outer boundary) where *p*(*R*) = 1, and Γ_*enzyme*_ (the enzyme surface) where *p*(*R*) = 0. This base association rate can be obtained through analytical or numerical solutions of a reaction-diffusion equation, given suitable boundary conditions:

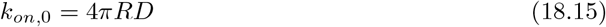

When a potential of mean force is important to the association event, the Smoluchowski equation is often used [115]:

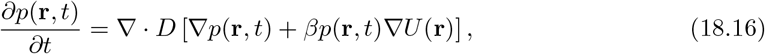

where we characterize the particle concentration with respect to position, **r**, and time, *t*. Here, *D* is the diffusion coefficient, *β* ≡ (*k*_*B*_*T*)^−1^ is the inverse thermal energy, *U*(**r**) is a drift term from the interaction potential of mean force, and *p*(**r**, *t*) is normalized: ∫ *p*(**r**, *t*) *d***r** = 1. Although it is beyond the scope of this review, it can be shown that an ensemble of particles undergoing Brownian motion subject to *F*(*r*) = −∇*U*(*r*) can be expressed as a continuum diffusion equation via Eq. 18.16 under specific conditions [116].

Electrostatic interactions are typically the basis of PMFs far from the transient encounter complex, given their slow decay as a function of distance. Here, the Poisson-Boltzmann equation, which relates charge distribution, ionic strength, and boundary conditions (such as molecular surfaces) to the electric potential is a standard model [117]:

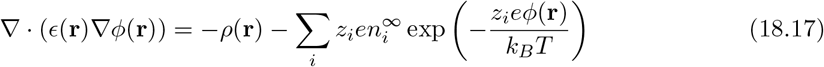

In this equation, the term within the summation reflects the bulk solution’s ionic strength and the potentials decay away from the charged surface. The resulting potential is multiplied by a substrate’s charge to yield the electrostatic energy, *U* = *qϕ*. This equation is commonly evaluated with tools like DelPhi [118] and APBS [119]. To quantify their effect on *k*_*on*_, an additional term to the Debye model can be shown [120] to give [114, 121]:

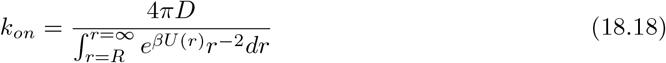

In this expression, the lower integration limit *R* represents the collision distance, defined as the radius of closest encounter between the two interacting molecules. More generally, *k*_*on*_ can be determined from the total flux to a reactive surface [120]:

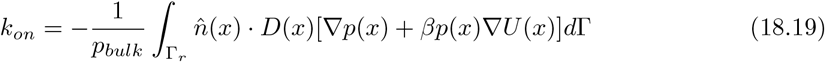

where Γ_*r*_ is the reactive region on the enzyme surface (Γ), 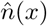 is the surface normal along Γ, *x* ∈ Γ and *p*(*x*) is the steady-state solution to Eq. 18.16. We note that the interaction potential can have significant impacts on both *k*_*on*_ and *k*_*off*_, as a potential gradient that favors association can result in a deeper energy well that impedes the escape of bound substrates, which would reduce *k*_*off*_. A limitation of these approaches is that they are usually restricted to idealized geometries.

##### 5.2.1.2 Brownian Dynamics

Brownian dynamics is a prominent approach for streamlining the calculation of diffusion-limited association kinetics (*k*_*on*_), especially for atomistically-detailed proteins [6, 122, 123]. BD simulations rely on coarse-grained representations of the reactants to model their diffusional approach from a dissociated configuration to form the transient encounter ensemble [122]. The corresponding association rate is proportional to the number of successful collisions, usually by defining a reaction criterion based on pairwise interactions between the substrate and target. Given the coarse-grained representations, BD simulations can model the evolution of particles over timescales spanning nanoseconds to microseconds. This is precisely the temporal regime for monitoring fast diffusive processes and binding events that would otherwise be difficult to capture with atomistic MD simulations. In these simulations, the interaction forces between molecules can be defined arbitrarily, though in practice they are often derived from an electrostatic potential via the Poisson-Boltzmann equation [123].

The Northrup-Allison-McCammon (NAM) algorithm [6] is among the most well-known schemes for computing association rates from BD simulations (Figure 5). These simulations typically start from experimentally-determined structures (e.g., X-ray crystallography, NMR spectroscopy, or cryo-EM) or from predicted models generated by docking, homology modeling, or ab initio methods. This approach couples short-range BD simulations with analytic estimates for diffusion far from the active site, enabling efficient estimation of association rates. NAM simulations begin by evolving many BD trajectories, starting with molecules randomly placed on a “*b*-sphere” of radius *b* around a second, fixed molecule. Each diffusive trajectory is simulated using Langevin-like equations of motion until the molecules either diffuse away to an outer “*c*-sphere” or meet the predefined binding criteria. The overall rate is determined in part from the fraction of trajectories that successfully reach the bound state. A key parameter *k*(*R*), the diffusion rate at separation *R* within the *b*-*c* layer is also used to describe the centrosymmetric, far field contribution that abides the Smoluchowski expression in Eq. 18.15.

**Figure 5:**
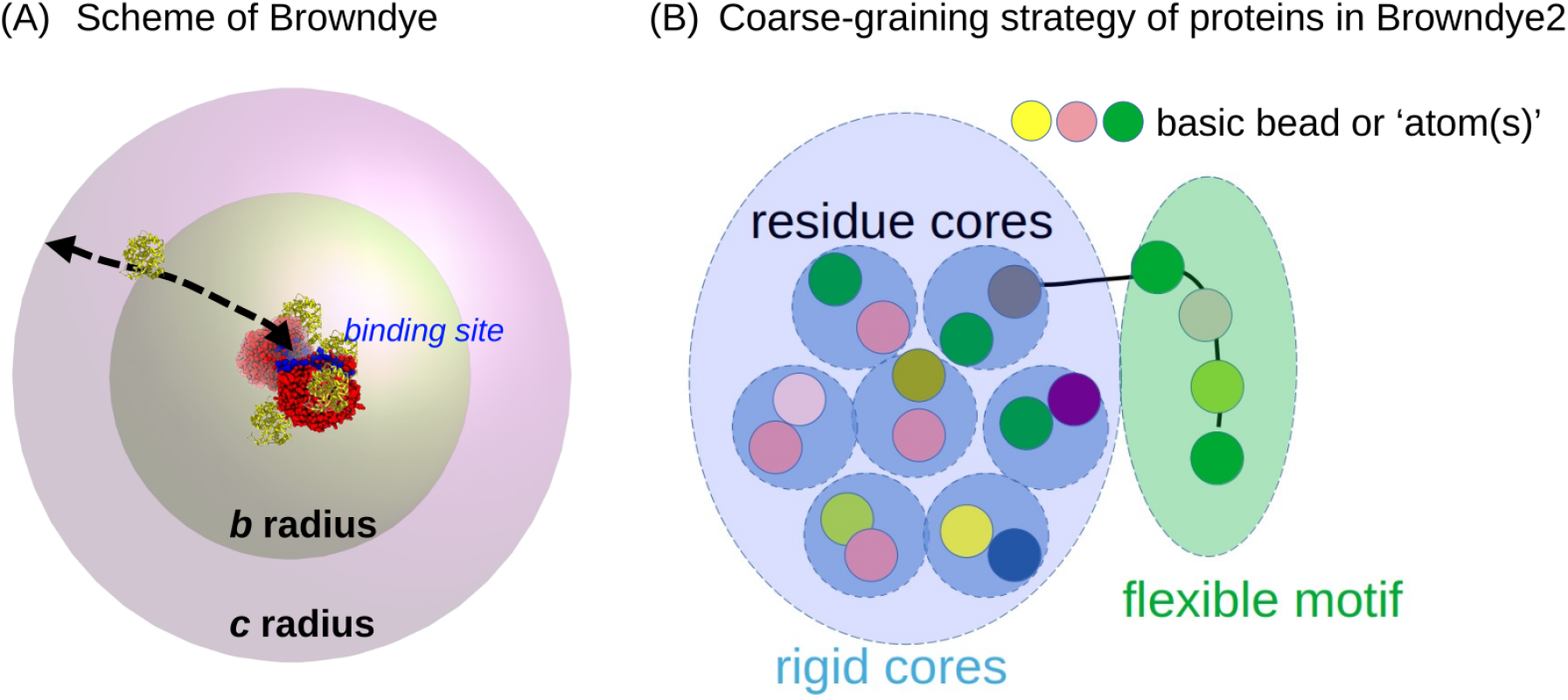
Northrup-Allison-McCammon simulation framework. The NAM framework [6], implemented in the software Browndye and Browndye2 [122], uses BD to calculate *k*_*on*_for protein-protein association. (a) The *b*-sphere and *c*-sphere that mark the starting position of the binders as well as the binder’s “escape” boundary. By setting reaction criteria that constrain the ligand to bind to a pre-defined binding site, the effects of conformation selection can be accounted for in the calculated *k*_*on*_ via Browndye. (b) Browndye2 utilizes a special coarse-graining strategy to achieve efficient simulation of diffusional encounter statistics, as well as to account for conformational flexibility during binding by treating parts of the binders as flexible chains/motifs.

An early example from Northrup et al. [124] demonstrated through BD simulations of protein-protein association involving reactive patches that the viscosity of the solvent traps the transient encounter complex. This kinetic entrapment accelerates the formation of the functional complex by allowing more opportunities for re-collision between the proteins [124]. This solvent effect helps explain the fast measured association rate constants (*k*_*on*_) of 0.5–5 × 10^6^ M^−1^s^−1^ for protein-protein associations that have restrictive geometric requirements [124]. Similarly, Gabdoulline et al. used BD to predict the association rates for five globular protein-protein pairs with varying degrees of intermolecular electrostatic steering, showing that BD can accurately predict *k*_*on*_ values that align with experiment [125]. More recent BD packages commonly reported in the literature include UHBD[126], SDA7[127], Browndye[122], BD-BOX[128], ReaDDy2[129], GeomBD3[130], as well as hybrids that combine BD and MD (e.g., SEEKR[131]). These packages differ by their specific target applications, the level of approximation they use, and the features they prioritize. For example, Browndye is specialized for predicting electrostatically-steered binding rates, while ReaDDy2 is designed for reaction-diffusion networks. An excellent review summarizing these BD-based packages is provided by [32].

While BD is computationally more efficient than MD, it can still present burdensome computational expenses, especially for large or highly flexible systems. To address this computational cost, variants of the NAM algorithm, such as weighted-ensemble BD [132] and biased-BD [133], have been developed. These methods accelerate the sampling of binding events, either by preferentially exploring trajectories that move toward the bound state or by applying a biasing force to speed up the process.

Although the NAM algorithm and BD as a whole are primarily used for computing association rates, there are a few examples of BD being applied to evaluate *k*_*off*_ . These cases typically apply to events where dissociation occurs through a “barrier-less” diffusive escape, such as using BD to simulate microtubule disassembly [134]. However, many biological systems have nontrivial and sometimes multiple transient encounter states, or significant barriers bridging the transient and post-encounter states, which would generally challenge conventionally BD approaches. To address more general kinetic modeling of biological systems, we now introduce a few frameworks for estimating kinetic rates that complement or extend BD simulation approaches.

###### 5.2.1.2.1 Kramers’ Theory

Extending on the models for the diffusional encounter, we now introduce Kramers theory to estimate the rate of escape of a bound particle over a potential energy barrier (see Figure 3). A model developed by Eyring in the 1930s called Transition State Theory (TST) provided an estimate of the escape rate for a particle 1) equilibrated in a potential well representing the bound state with a curvature of *ω*_*a*_ and 2) confined by a barrier of amplitude Δ*G*^‡^ [135] (see Figure 3a). Kramers extended this idea by assuming that the particle additionally undergoes Brownian motion with viscous drag, while the barrier assumes a parabolic PMF. Under those assumptions, it was shown that

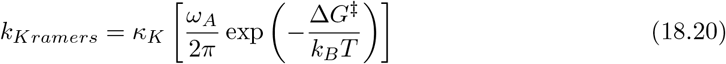

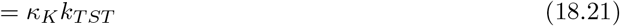

where the short-form of Kramers transmission coefficient *κ* represents the influence of friction on the observed escape rate [136] In the high-friction limit most appropriate for biological systems, *κ* → *ω*_*b*_*/γ*, where *ω*_*b*_ reflects the particle’s oscillations at the barrier. As the friction *γ* increases, *k*_*Kramers*_ is reduced relative to the rate from TST (*k*_*TST*_). In the low-friction limit, *κ* → 1, upon which *k*_*Kramers*_ → *k*_*TST*_. While the model was originally formulated for escape (dissociation) processes, it can also be extended to association phenomena when an effective free-energy barrier governs the encounter [137]. A comprehensive review of this theory is available for a further elaboration of these concepts [136].

The Kramers approach differs from the Brownian dynamics framework as the potential energy surface (PES) governing the diffusive encounter is replaced with a single barrier between bound and unbound states. It can, however, be used in conjunction with BD if the most significant barrier along the reaction coordinate sits between the diffusional encounter and bound states. This perspective was used by Berezhkovskii et al. for BD simulations of ligands targeting buried binding sites, as the corresponding free energy barrier to binding entry could be represented by a simple one-dimensional potential of mean force [138]. Although this study assumed a constant diffusion constant along the reaction trajectory, refinements that include a spatially-varying, local diffusion coefficient along the PMF can further improve rate estimates [139].

##### 5.2.1.3 Committor Probabilities

The reaction path in Figure 3a is a simplification of a binding process that is amenable to treatments like Kramers’ theory. A more representative ligand/protein association landscape for biomolecules is represented in Figure 3b, which includes a yellow reaction coordinate linking the transitional encounter and bound state. This tortuous path traverses saddle representing barriers to association that limit direct application of simple models like Kramers’ theory. In this situation, committor probabilities provide a more general mechanism to both identify transition state barriers along a reaction path and to accommodate free energy barriers of arbitrary curvature. Importantly, the use of commitors enables the estimation of barrier crossings obtained from molecular simulations. Rather than relying on a predefined reaction coordinate, the committor probability (*p*_*B*_(**x**)) quantifies the likelihood that a trajectory initiated from a configuration (**x**) will reach the product state *B* before returning to the reactant state *A*:

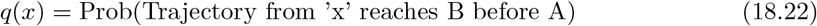

where we use *q*(*A*) and *q*(*B*) to signify the unbound and bound states, respectively. The transition-state surface is then defined by configurations where the committor equals one half (*q*(*x*) = 0.5), signifying that the system is equally likely to commit to either state. The equilibrium probability density (*ρ*(*x*)) and the probability flux (*D*(*x*) ∇*q*(*x*)) are then evaluated along the ‘isocommittor’ surface where *q* = 0.5 to estimate transition rates for A to B. This is done by integrating along the isocommittor surface and normalization by the probability of being in *A*:

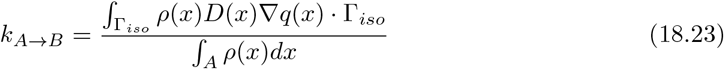

where Γ_*iso*_ is the isosurface along which *q* = 0.5. The reverse process can be estimated by leveraging a detailed balance criterion requiring the forward flux (*k*_*A→B*_*π*_*A*_) to equal the reverse flux at equilibrium [140]:

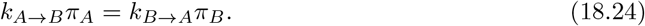

Here (*π*_*A*_) and (*π*_*B*_) are the equilibrium populations of states *A* and *B*, and (*k*_*A*_ *→* _*B*_) and (*k*_*B*_ *→* _*A*_) are the corresponding transition rates. The committor framework forms the theoretical basis for trajectory-based methods such as milestoning [141].

###### 5.2.1.3.1 Markovian Framework

Markov state models (MSMs) provide an alternative approach for predicting binding kinetics, by relying on a predefined set of microstates representing fully-bound, fully-dissociated, and intermediate states. These strategies discretize the state space into regions that permit transitions from between system states, independent of their origin, thereby linking long-lived and functionally important conformational states of proteins. Ideally, the discretization provides a suitable representation of a reaction pathway that encompasses biologically relevant states (e.g. fully-dissociated and bound conformations).

The Markov state modeling approach provides a transition probability matrix, **T**(*τ*), for evolving a given microstate population, P(t) at time t, by a lag time *τ* to obtain the updated probabilities, P(*t* + *τ*):

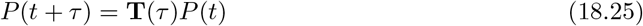

The resulting system satisfies conditions such as detailed balance between states *P*_*i*_ and *P*_*j*_ when at equilibrium:

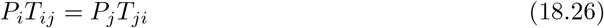

and assumes that transitions from a given *t* to *t* + *τ* are independent of the preceding steps.

Given a Markovian model that encompasses functionally-relevant states of a system, steady-state and time-dependent probabilities for a microstate configuration can be readily determined. Relaxation rates for a microstate configuration to its equilibrium distribution can also be estimated from the eigenvalues, *λ*_*i*_, of **T**(*τ*), using *k* = − *τ* ^−1^ ln *λ*_*i*_.

One important quantity obtained from Markov state models is the mean first passage time (MFPT), which can be used to describe the kinetics of a binding event. In this context, the time, *τ*_*MFPT*_, for the system to progress from an arbitrary configuration to an ‘absorbing’ microstate, *P*_*i*_, is evaluated by setting to 0 all probabilities for transitions leaving *P*_*i*_, e.g. *T*_*ij*_ = 0, ; ∀*j* ≠ *i* [142]. This yields a different fundamental matrix, from which *τ*_*MFPT*_ can be obtained for any starting state to the absorbing state. When the absorbing state corresponds to a bound configuration, the resulting MFPT can provide rate constants via:

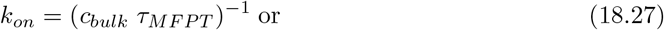

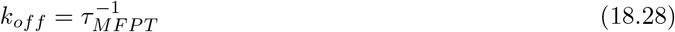

where *c*_*bulk*_ reflects the ligand concentration for association rate equation; the latter equation for the dissociation rate is used when the absorbing state corresponds to an unbound configuration.

While a single simulation may be limited to tens of microseconds, Markov State Models (MSMs) [143] and related frameworks for dimensionality reduction and deep learning, such as VAMP [144], VAMPnets [145], and TICA [146, 147], have enabled the synthesis of extensive MD data from independent simulations into kinetic models. In effect, this strategy provides a basis for coupling short MD trajectories to estimate long-timescale kinetics. A conceptually similar approach is used in the multi-scale method SEEKR [131] (discussed in Section 6.3.1.2).

###### 5.2.1.3.2 Survival Probability

There are many examples, such as in the gating of ion channels, where a comprehensive definition of microstates as part of an MSM is infeasible. Instead, transition probabilities for canonical states can be determined, such as from patch clamp experiments of single channel currents reflecting a channel’s open versus closed states [148]. Formally, the survival probability, *s*(*t*), reflects the likelihood that a particle remains in a specific state over a given time interval, *t*. [149]. For the simple example of a dissociating complex, the survival probability can be represented as a single exponential decay, for which the integral over *t* = 0 to *t* = ∞ provides an estimate of that decay rate via [149].

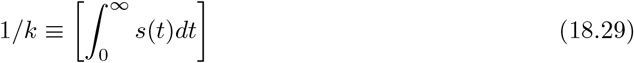

The survival probability has the advantage of being simple to evaluate from molecular simulations, as it essentially entails evaluating the duration of a complex’s association over a large ensemble of simulation. In the event that dissociation is well-described as a simple two-state process with mono-exponential decay, Eq. 18.29 also provides the “residence time” of the ligand as:

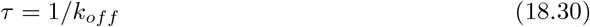

which is frequently used as a metric for drug efficacy [150].

##### 5.2.1.4 Beyond the Spherical Cow: Anisotropic Reactivity and Dimensionality

###### 5.2.1.4.1 Effects of Dimensionality (2D vs. 3D)

Naturally, most biological binding events are expected to be topologically different from the symmetric system assumed in simple spherical models [11]. Even the dimensionality of the system, such as a 2D plane or a bilayer as shown in (Figure 2c), versus an open, 3D system, can impact the rate estimate or whether it converges. For instance, the diffusion of quinone within the mitochondrial membrane to shuttle electrons between respiratory complexes [12, 13] (Figure 2a) is amenable to analytically-solvable 2D diffusion models under strict boundary conditions [14]. This concept was neatly framed for a molecule adsorbing onto a surface, after which its diffusion within the plane of the surface enhanced its sampling of the domain [151]. This enhancement was due to the molecule’s effective confinement by dimensionality reduction, i.e. the 2D diffusion of the particle within a plasma membrane, which increased the re-encounter probability and thereby, its association with transmembrane targets [152, 153] (Figure 2d). It is worth noting that the 2D diffusion rate in eukaryotic membranes is sensitive to factors including the lipid phase, cholesterol content, and presence of obstacles like other proteins [154].

Relatedly, receptor tethering to or embedding within the plasma membrane is another mechanism for confinement that can enhance diffusion-limited association, as seen with the TNF*α* ligand and its cognate receptor [155]. Chang and colleagues used BD simulations to investigate the binding of complementary ssDNA strands to ssDNA molecules immobilized on a surface, providing insights into the kinetics of surface-tethered biosensors [156]. The binding of endogenous, extracellular ligands to GPCRs on the plasma membrane represents another classic situation involving extracellular ligands diffusing in 3D to react on a planar membrane, often with a finite-sized reactive site [17] (Figure 2e). One-dimensional diffusion-controlled binding events are also frequently encountered [19], such as the search a transcription factor performs while sliding along DNA to locate its specific promoter sequence, or binding events involving an ensemble of intermediate states [20]. Adapting BD simulations for these paradigms can be as simple as imposing different boundary conditions (e.g., spherical vs. planar) or using alternative coordinate systems, such as cylindrical coordinates for filaments. However, these non-Cartesian coordinate systems are often limited to idealized geometries.

Modern continuum methods and Brownian dynamics simulations with rigid bodies can be used to account for complex protein shapes and electrostatic interactions. For BD, this can be as arbitrary as defining a boundary representative of a protein target’s geometry. Along these lines, continuum methods, namely partial differential equation-based models of diffusion, commonly use the finite element method to account for the surfaces of complex target geometries [157]. Such continuum approaches define specific reactive patches on a protein or cell surface as boundary conditions for evaluating diffusion-limited association rates.

#### 5.2.2 Modeling Flexible Solutes

Other types of binding events include ligands binding to deep pockets within a target protein or flexible, intrinsically disordered regions (IDRs) binding to globular protein partners. For these systems, the conformational flexibility of the associating proteins can be the deciding factors for the association rate constants. More generally, enzymes undergoing conformational changes during substrate recognition or gating will present binding rates that differ from treatments assuming rigid binding partners. Mechanistically, depending on timing of those conformation changes, the association can fall into a conformational selection regime, an induced fit regime, or somewhere in between these two extremes [20]. Conformational gating and selection refer to mechanisms where dynamic regions of a protein transiently expose binding clefts or ligands adopt target-compatible conformations that can modulate both *k*_*on*_ and *k*_*off*_ [51]. Induced fit corresponds to when either substrate and target undergo conformation changes after association. These binding mechanisms have distinct impacts on the association rate.

Accurate predictions of binding kinetics for flexible reactants require accounting for the rates of the binding pocket’s conformational gating and the intramolecular conformation sampling of the substrate, like an IDR. For example, the HIV-1 protease [158] samples distinct conformations where the binding pocket flips between occluded and accessible configurations, which renders binding as a conformational selection process. In contrast, induced fit is evident in the binding of p53 to its cognate DNA motif, as DNA binding induces a loop conformation change in p53 that gives a kinetic advantage to cognate over nonspecific DNA binding [159]. IDR binding on the other hand is often a mixture of conformational selection and induced fit. Specifically, IDRs can retain residual structure (partially folded elements) while remaining highly flexible. Some reports indicate that this residual structure accelerates the coupled binding and folding of IDRs to their targets [160], while others have reported no significant effect [161]. These examples therefore highlight the case-dependent, complex mechanisms governing their association events.

##### 5.2.2.1 BD solutions for Kinetics Prediction of Flexible Binding Events

While early BD studies used simple descriptions of flexibility of the active site to estimate binding rates [162, 163], BD is largely ignorant of important atomistic details that can significantly impact the kinetics; one such example is found in the impact of trapped solvent on loosely-associated proteins, which needs to be squeezed out for the transition to the ‘native complex’ from the post encounter stage (stage 2 in Figure 4). In light of which, BD simulations of binding events involving flexible molecules benefit from including a wide variety of structures and configurations from experiment or MD simulations to serve as inputs to BD approaches.

Toward this end, extensions of the NAM framework of BD have incorporated conformational flexibility in kinetics characterization. One example using the BDflex method [164], which considered a spherical protein with a buried pocket that was divided into an exterior and interior space. The ligand’s diffusion in the exterior region and its encounter rates at the exterior-interior border were simulated by conventional BD simulations. Concurrently, the interior region of the pocket was assumed to switch between a reactive and an unreactive configuration with pre-defined rates. The dividing surface between the exterior and interior region was treated by a radiative boundary condition to interface the substrate flux from BD with the interior region [164]. This simplified model was sufficient to capturing the impact of pocket flexibility on predicted association rates.

###### 5.2.2.1.0.1 Prediction of Complex Ensembles from BD simulations

BD and related simulation methods can also be used to generate equilibrium conformation ensembles for highly flexible proteins in isolation, which can then be used as starting configurations or representative rigid-body states in more complete simulations of binding. This is particularly important for proteins exhibiting conformational gating or intrinsic disorder, which can impact association pathways and transient encounter complexes. While most applications in this regard tend to rely on atomistic-resolution simulations of flexible proteins or IDRs, progress has been made with coarse-grained (CG) models that can reduce the computational expense [165]. For example, one study performed extensive coarse-grained Langevin dynamics simulations of IDPs, by restricting the model to three energy terms (steric repulsion, attractive hydrophobic interaction, and electrostatics), which yielded IDP conformations that agreed well with FRET-measured residue-residue distance data [166]. Another study used COFFDROP, which is a simulation approach based on an implicit-solvent force field [167] that derives nonbonded and bonded CG potentials from all-atom MD simulations of two-residue fragments. When coupled with Brownian dynamics, this force field succeeded in recapturing the hydrodynamic radii of eight IDRs, including an A*β* fragment, ProT*α*, and *α*-synuclein [167]. In a related work, internal coordinates of chained particle positions were propagated using Brownian dynamics [168] to study the aggregation dynamics of *α*-synuclein. For that study, the *α*-synuclein protein was represented as either a sphere with layered interaction surfaces representing the disordered state or a spherocylinder with an attractive surface representing the folded *β*-sheet state. In addition, a Brownian dynamics model for IDRs based on bead-rod chain dynamics, inclusive of hydrodynamic interactions and excluded-volume effects, has also been developed [169]. This model was used to simulate the internal dynamics of synthetic (GS)_*n*_ IDRs ranging from 11 to 40 residues.

##### 5.2.2.2 Comparison of Kinetics Obtained from Pure MD Approaches

While BD is most appropriate for modeling the loose assembly of substrate/target complexes, MD-based methods remain indispensable for computing the final assembly of the complex, especially when protein flexibility, solvent organization, or ions play critical roles [32]. As discussed in detail by Wang et al. [170], physics-based methods include brute-force MD, MD with Markov state modeling, and Biased MD (such as LiGaMD [171]); these methods typically require extensive simulations [59, 172]. In recent years, unbiased MD simulation trajectories spanning tens to hundreds of microseconds have been made possible via specialized computer architectures; the Anton supercomputer, as an example, has been used to identify small-molecule binding pathways for GPCRs [172] and protein kinases [173]. By explicitly simulating the substrates’ approaches toward their targets in atomistic detail, the Anton simulations circumvented the need for prior knowledge of the transient encounter complexes that are required for BD approaches.

Nevertheless, pure brute-force MD simulations are not ideal for predicting binding kinetics, as they require sampling a sufficient number of binding and unbinding events, which are often rare on typical simulation timescales. Biased sampling strategies improve upon conventional MD simulations by focusing trajectories that are most likely to lead to successful substrate/target binding. Many enhanced sampling methodologies are reported in the literature, which tend to use either biased potentials (umbrella sampling, metadynamics etc.) or weighted ensemble methods [66]. Representative methods include Ligand GaMD (LiGaMD) [171], random acceleration MD [174], metadynamics [175], and WExplore [176], which have been used to calculate *k*_*on*_ and *k*_*off*_ from detailed binding/unbinding pathways, as summarized in [177]. Other recent developments in this area include ModBind, which afforded the estimation of *k*_*off*_ values ranging from 10^0^ to 10^−6^ s^−1^ [178]. This approach entailed running simulations at an elevated temperature (e.g., 1000 K) to accelerate unbinding, followed by reweighting the trajectories to recover the conformational distribution at the physiological temperature [178]. As computational resources grow and enhanced sampling techniques mature, predicting kinetics (*k*_*on*_ and *k*_*off*_) for ligands binding to medium-sized proteins like trypsin and Src kinase on microsecond timescales using biased sampling is becoming increasingly feasible [66].

### 5.3 Transient Encounter: BD for Heterogeneous Media

#### 5.3.1 Factors in Heterogeneous Media

Heterogeneous media encompass mixed solutes and solvents and are characterized by physicochemical properties such as free-volume fraction, electrostatic potential, and the spatial distribution of enzymes, all of which can influence reaction kinetics. In turn, the kinetics of binding in heterogeneous media can be influenced by all of these factors, as observed under conditions of macromolecular crowding within the cell cytosol, which can slow diffusion, but also trap reactive intermediates to accelerate reactions, depending on the crowders’ physicochemical properties. Local attractive potentials can further steer substrates or drive co-localization of enzymes in cellular compartments or LLPS condensates (discussed below) to concentrate reactants and thereby improve reaction efficiency [3] (see also Figure 7). Together, these factors dictate the local ligand concentration and, consequently, the timescale and amplitude of events contributing to cell signaling cascades (see Eq. 18.5). For example, by monitoring the diffusion trajectories of proteins that form stress granules (SGs), Zhang et al. [94] showed that under SG formation stimuli, proteins have a reduced diffusion coefficients (*D*) compared to normal conditions. Under both conditions, evaluating the mean-squared displacement versus time (MSD ∝ Δ*t*) relationship showed that while proteins exhibit a broad range of motion types (e.g., normal vs. confined diffusion), the predominant motion type remains Brownian [94].

##### 5.3.1.1 Crowding

Protein diffusion in the cytosol often occurs under crowded conditions, which can lead to reduced diffusion and in many cases, anomalous diffusion [183]. The effects of crowding in cells are complex, as reviewed by Zhou, Rivas, and Minton [184]: crowding can hinder association by increasing viscosity or imposing physical barriers, but also promote association by modulating the available volume fraction [185].. Excluded volume effects can also promote association by reducing the translational entropy to be gained when reactants dissociate [186]. The net impact of crowding on kinetics is a balance between these competing effects [184].

Mixtures also present complex crowding effects depending on the physicochemical properties of the solutes themselves as well as their interactions with one another. Hence, crowding impacts not just diffusion but also the conformational landscapes of enzymes and the transition states of a reaction. This can arise from the excluded volume shifting the conformational ensemble toward compact states that may alter binding site accessibility. In other words, excluded volume effects can confine reactants, which can increase the reaction rate, though the crowding can also slow overall diffusion, suppressing both *k*_*on*_ and *k*_*off*_[184]. For a thorough discussion of simulation with solutes in crowded environments, we refer the reader to Zhou et al. [184].

##### 5.3.1.2 Liquid-liquid Phase Separation

The presence of different phases, such as in liquid-liquid phase separation (LLPS) (see Figure 7), also influences binding kinetics. For instance, reactions can occur in distinct organelles or phase-separated entities like LLPS droplets, as exemplified by eukaryotic cells’ use of compartments, including mitochondria and membraneless germline P granules to localize reactions [187] (Figure 2b). For LLPS, reaction rates are uniquely shaped by the condensate’s composition, which can increase local concentrations of favorably partitioned species, exclude competing species, and alter the diffusion behavior of a substrate within the condensate compared to its boundary [188, 189]. Additional considerations include how the physicochemical properties within the condensates themselves can alter catalysis. For example, changes in solvent viscosity and substrate/active site solvation properties can directly modulate enzyme catalytic rates and specificity [190]. In addition, the condensate environment can influence the nature of the conformational ensembles, which in turn could impact enzyme kinetics, such as that reported for the MORC3 and DDX proteins, where phase separation activates their enzymatic activities [191, 192].

#### 5.3.2 Modeling Reactions in Heterogeneous Media

##### 5.3.2.1 Crowded Environments

Crowded environments motivate using effective diffusion coefficients to implicitly represent the influence of crowders on solute diffusion. The Maxwell-Garnett relationship is a simple approximation that can be used to quantify excluded volume effects [193] that modulate diffusivity as a function of the free volume fraction, *ϕ* ∈ [0, 1] e.g.

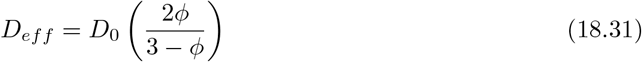

for spherical inclusions much bigger than the solute [193]. These coefficients can be numerically estimated from explicit MD or BD simulations by computing positional or velocity autocorrelation functions (Eq. 18.11) for diffusing crowders [110], such as through the Green-Kubo relationships discussed earlier.

There are numerous studies that have used Brownian dynamics simulations to investigate protein-protein association rates within crowded environments (see depictions in Figure 6d.) In one study, simple hard spheres were used for a suspension of crowders through which the Brownian motion of two proteins was simulated with pre-calculated diffusion coefficients [194]. In essence, crowding effects were accounted for by a simple model of hard-sphere repulsion between the proteins and crowders. With this simplified model, the authors showed that the reduced free volume due to the crowders accelerated association rates by up to two-fold compared to the uncrowded case. Brownian dynamics can additionally simulate crowding [184] by including explicit crowder molecules as mobile or fixed entities. Under some conditions, the solute’s interactions with crowders or the crowders’ mobility can give rise to anomalous diffusion [195].

**Figure 6:**
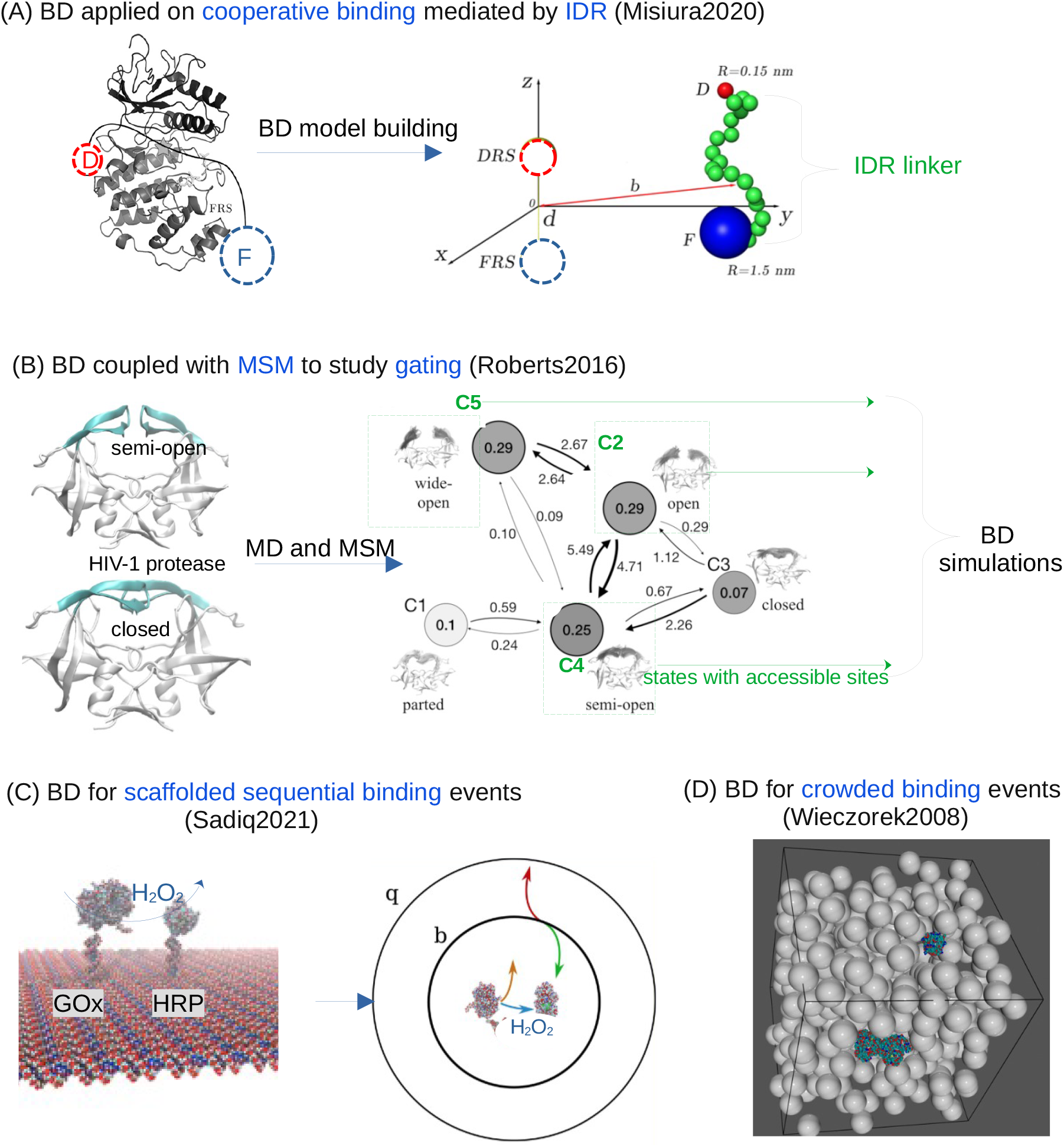
Brownian-type motions dominate LLPS dynamics. (a) Experimental observation via imaging. Imaging techniques are used to capture experimental data on LLPS dynamics, such as the diffusion trajectories of individual molecules [94] or time-dependent changes in droplet volume [179]. (b) Modeling with Brownian motion-induced coalescence (BMC). The BMC model [180] serves as a fundamental framework for explaining LLPS dynamics in living cells, including the formation of nuclear bodies [179, 181] and transcription factor condensates [96]. Brownian motion provides a foundational model upon which additional potentials (e.g., for collision or dissociation) and modeling techniques (such as lattice models) can be integrated to explain case-specific LLPS data [182]

**Figure 7:**
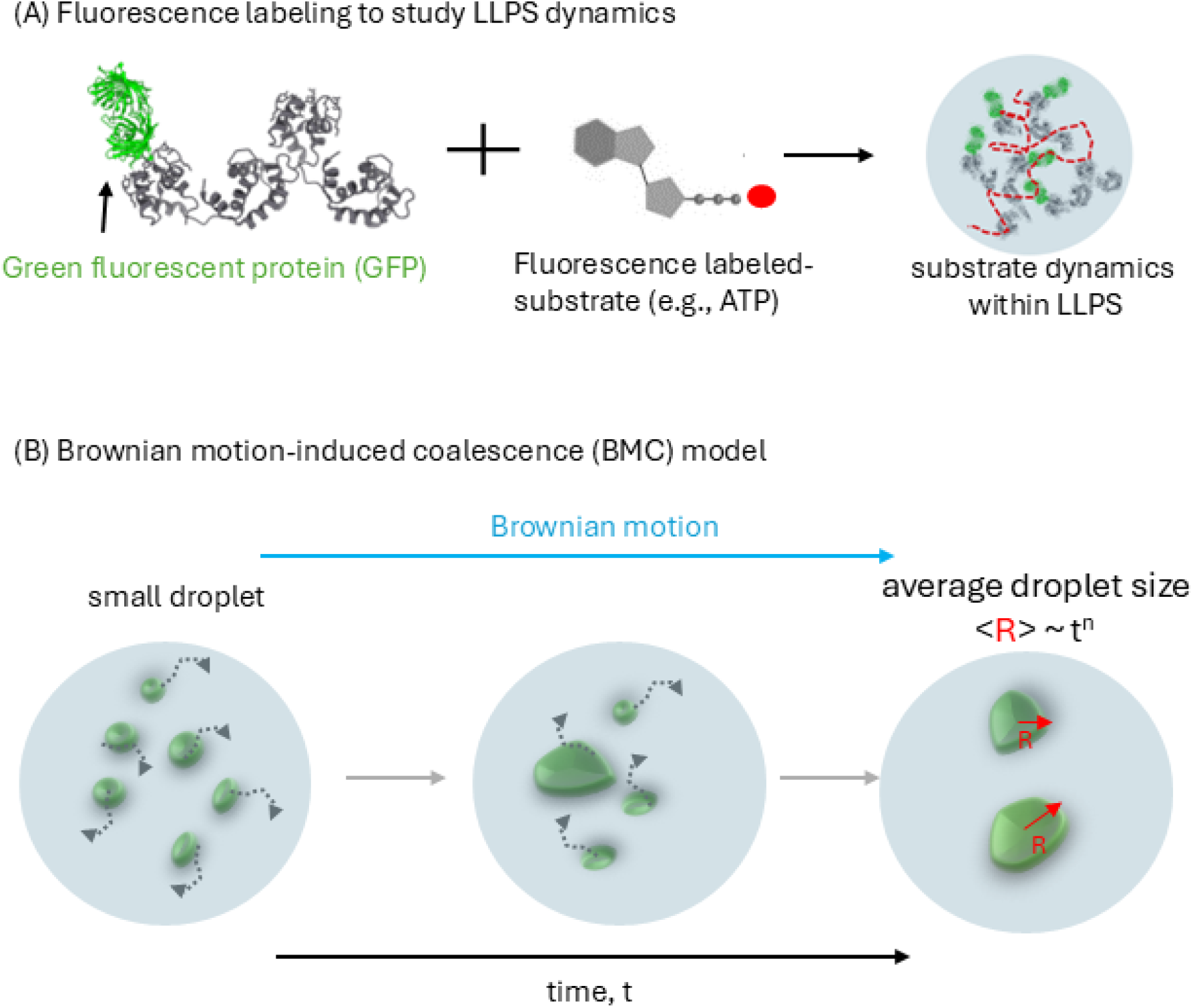
Exemplary BD applications for calculating binding rates under biologically complex conditions. (a) Multi-site binding involving the ERK2 enzyme. ERK2 possesses two binding sites for the two domains of the EtsΔ138 protein, which are connected by an IDR linker. BD simulations by Misiura et al. demonstrated that this IDR tethering accelerates association rates by 3-to 4-fold [196]. Adapted with permission from [196]. (b) Hybrid BD/MD/MSM approach for ligand binding to HIV-1 protease. Sadiq et al. coupled BD with MD and MSM to accurately calculate binding rates [158]. MD and MSM first identified the binding-site-accessible microstates of the ligand, which were then used as starting points for rigid-body BD simulations. The final binding rate incorporates both the BD-simulated diffusion rate and the conversion rates between binding-site-accessible and inaccessible microstates. Adapted with permission from [158]. (c) BD simulation of scaffolded sequential binding. BD has been applied to study sequential reactions, such as the transfer of H_2_O_2_ between the GOx and HRP enzyme pair, where H_2_O_2_ is a product of GOx and a substrate for HRP. Simulations elucidated the impact of enzyme separation distance on the H_2_O_2_ transfer rate [197]. Adapted with permission from [197]. (d) BD for protein association in crowded environments. BD simulations have been used to investigate how macromolecular crowding affects protein-protein association rates. For example, Wieczorek et al. modeled crowders as hard spheres and simulated the diffusion of two proteins using pre-calculated diffusion coefficients to study this effect [194]. Adapted with permission from [194].

Classical BD simulations typically assume an instantaneous decay of the velocity autocorrelation (VAC) function, which neglects memory effects that can give rise to anomalous diffusion. To accurately model such conditions, generalized Langevin dynamics frameworks that explicitly consider memory effects can be used. Furthermore, the presence of weak, nonspecific interactions between solutes and crowders adds another layer of complexity; these interactions generally slow diffusion or lead to off-target binding, though some configurations have been shown to enhance transport. The development of realistic, large-scale simulations of the cytoplasm, represents a significant step toward capturing those influences; simulations from Feig and colleagues for instance have provided atomistically-detailed representations of the crowded cellular interior relative to simpler models using explicit crowder spheres [198]. In a similar vein, the self-assembly of IDPs driving LLPS (such as ABLIM [199]) have been detailed using recent Langevin [200] and BD approaches [201].

Diffusion in crowded environments can also be modeled via “homogenization” approaches. Homogenization theory provides a strategy to compute effective transport parameters, such as a diffusion coefficient, that is valid for a large problem domain, based on solving a related problem on a smaller domain. [202, 203] (see Figure 8). In practice, the homogenized diffusion equation (Eq. 18.12) is solved in a representative volume element that reflects the crowder composition of the larger domain (Figure 9). The solution to that homogenization formulation yields an effective diffusion parameter that captures the influence of crowders on solute diffusion over the larger domain, without having to explicitly model the individual crowders. Previous studies have used this approach to model the impact of atomistic-resolution crowders on continuum-scale diffusion coefficients [204, 205], as well as for the diffusion of ATP based on a near-atomistic-resolution representation of the sarcomere [206, 207]; these estimates were later shown to agree well with ATP diffusion measurements in skeletal muscle cells [208].

**Figure 8:**
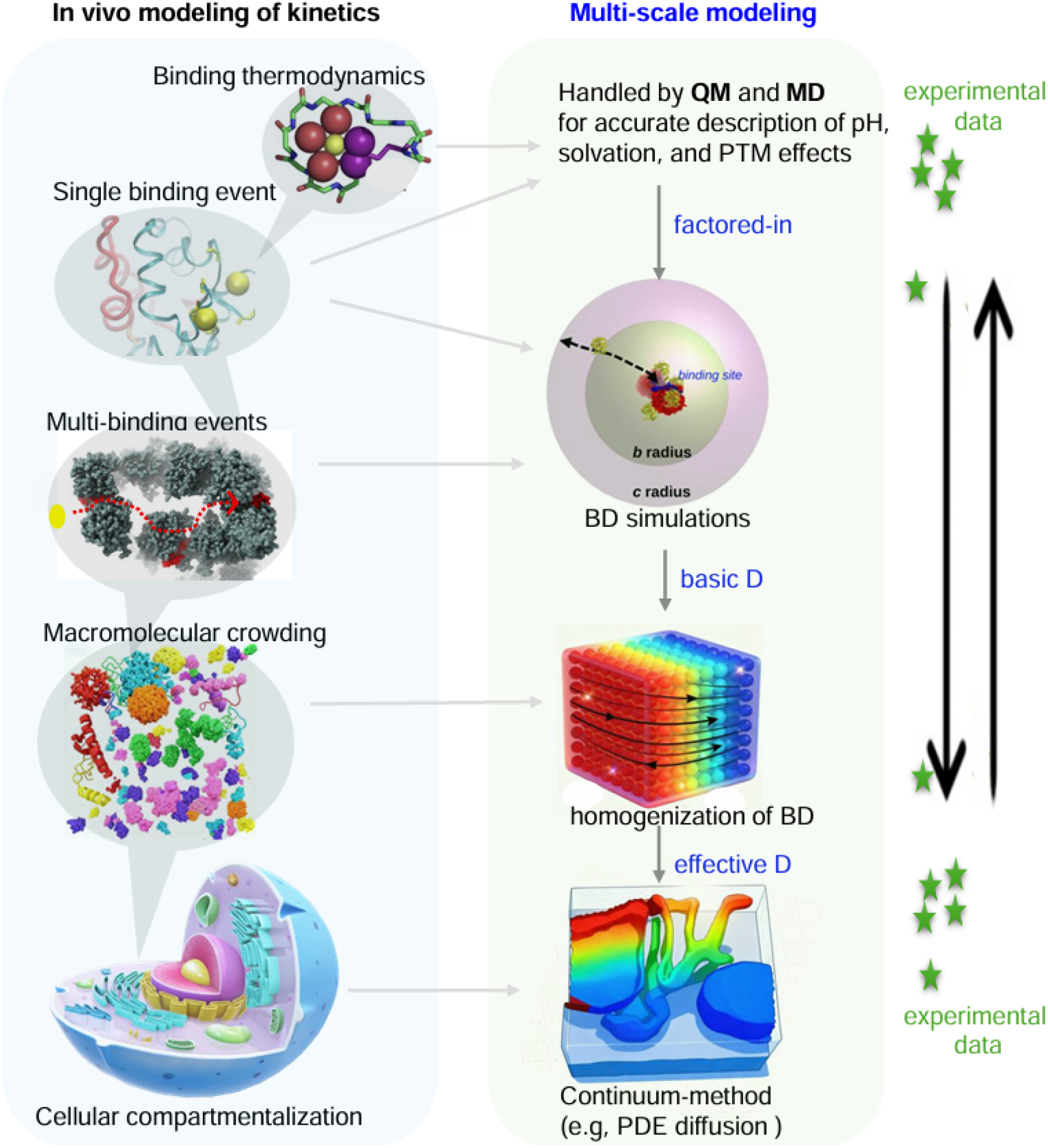
A bidirectional multi-scale modeling framework for predicting in vivo kinetics. In vivo binding is a complex process governed by multi-scale events, including cellular compartmentalization, macromolecular crowding, and coupled reactions like metabolic channeling. Furthermore, binding kinetics are modulated by local physicochemical factors such as ionic strength, pH, and solvation effects. To achieve comprehensive insights, a multi-scale computational framework is essential. We propose that such a framework can be formed by integrating existing computational methods across different scales, centered around Brownian dynamics (BD). The BD technique occupies a central position in the computational tool spectrum: at one extreme, QM/MM provides atomic-level detail but is limited to small scales; at the other, continuum models simulate cellular-level effective diffusion phenomena. Thus, BD can utilize QM/MM outputs for refined diffusion coefficient estimation, while BD results can serve as input for homogenization and continuum methods. Finally, we propose that such a multi-scale modeling framework should be bidirectional. This is because computational insights require experimental calibration, and because experimental data itself spans multiple scales, the entire framework must operate bidirectionally (e.g., bottom-up and top-down) to ensure consistency and utility.

**Figure 9:**
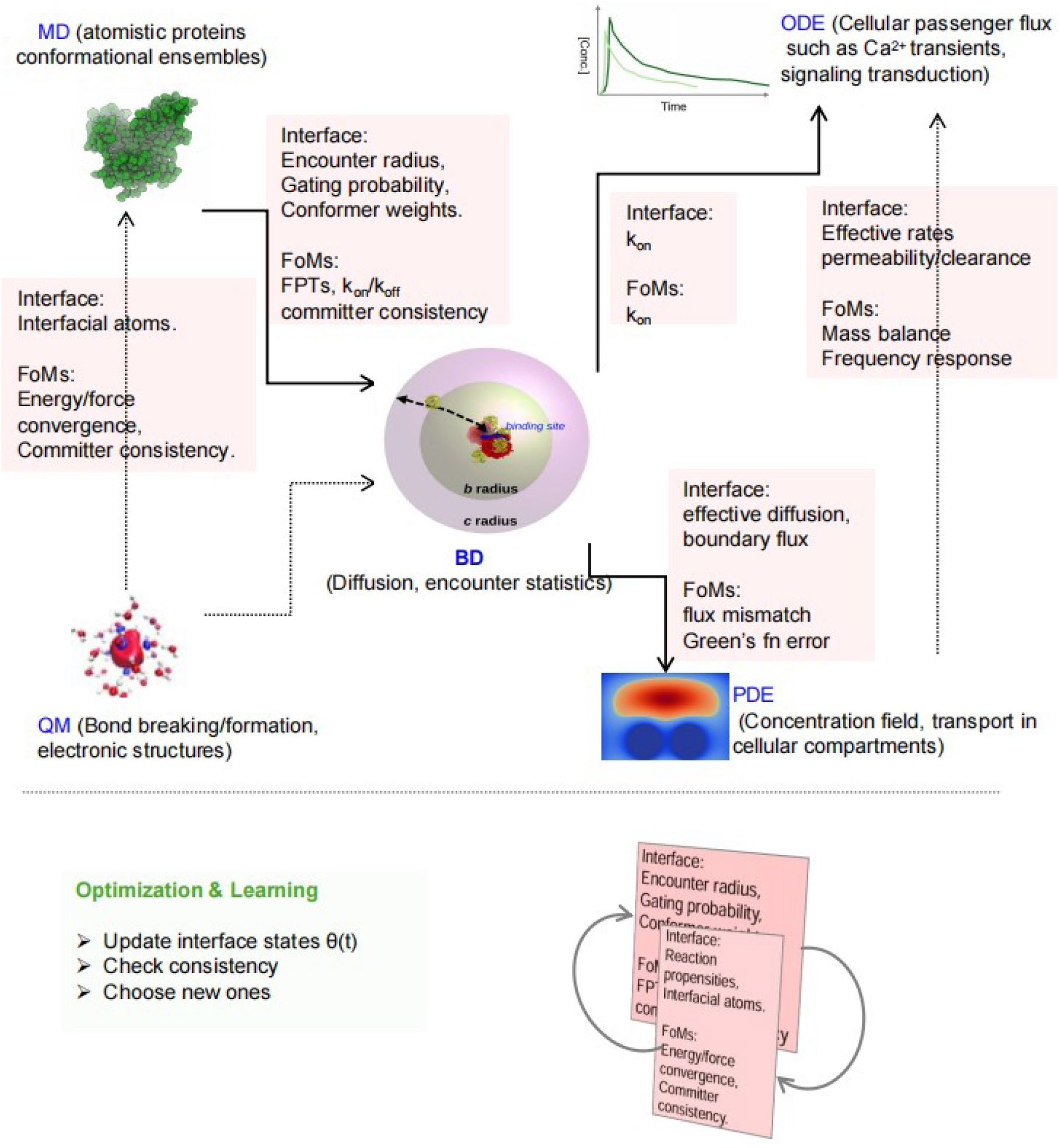
Multiscale coupling of biomolecular binding models. Schematic of a bidirectional modeling framework linking quantum mechanics (QM), molecular dynamics (MD), Brownian dynamics (BD), and continuum descriptions (PDE/ODE) of cellular transport and signaling. BD occupies a central mesoscopic role, translating atomistic structure and energetics (from QM/MD) into effective kinetic parameters (*k*_*on*_, *k*_*off*_) and fluxes that inform continuum transport and cellular response models. Interface variables and figures of merit (FoMs) at each scale enable consistency checks and provide targets for optimization, supporting a closed-loop, multiscale learning framework for predicting and controlling biomolecular binding kinetics in vivo.

Interestingly, the contributions of crowders can also be reflected implicitly in PMFs used by models for predicting the association rate (Eq. 18.19 as an example). This is advantageous for homogenization approaches, as it goes beyond reflecting the crowders’ shape by incorporating physical factors like long range electrostatics through effective potentials. Further, by changing the form of the Smoluchowski equation (Eq. 18.19) to resemble a standard diffusion equation, arbitrary mean-field potentials, such as PMFs from atomistic simulations [209], can be reflected in an effective diffusion coefficient [204, 210] appropriate for large-scale simulations using BD methods or continuum approaches.

##### 5.3.2.2 LLPS Modeling

BD is a powerful tool for capturing the co-assembly of solutes into a phase separate from their solvent (Figure 7). As a recent example, Zhang et al. [182] simulated LLPS dynamics by assuming each protein underwent free Brownian motion within a lattice. This allowed individual proteins to collide and form condensates, or dissociate from them, with probabilities determined by energy barriers for the events. Both the individual proteins and the resulting condensates were treated as Brownian particles, and by assuming the condensate’s diffusion coefficient changed dynamically with its size, the BD-Lattice model successfully simulated LLPS dynamics in a manner consistent with experimental data.

### 5.4 Post-encounter: Hybrid BD/MD Approaches

#### 5.4.1 Modeling the Post-encounter Event from Hybrid BD/MD Approaches

While Brownian dynamics excels at modeling the initial diffusional encounter, simulating the transition to the final bound complex often requires atomistic details best captured by molecular dynamics (MD) [211]. Consequently, modern frameworks have been developed to couple BD with MD-derived information in order to bridge the gap between the encounter complex and the native bound state [32].

##### 5.4.1.1 Incorporating MD Data for Barriered or Gated Events

To resolve binding mechanisms beyond simple diffusion, BD is frequently complemented with atomistic simulations that account for energy barriers and conformational gating. Here, the Berezhkovskii et al. [138] approach based on using model PMFs for the post-encounter event, and incorporating MD-derived position-dependent diffusion coefficients per Hinczewski et al. [139], are prime examples. For kinetic effects like conformational gating, BD is often coupled with Markov State Models (MSMs) of a target protein [158, 212]. If the timescale of the gating motion is slower than that of diffusion, the gate serves as a kinetic filter, reducing the association rate below the diffusion limit and conferring substrate selectivity based on internal dynamics [213]. Applications of these coupled approaches range from early models using simple gating factors [50] to complex systems including the gated binding of drugs to HIV-1 protease [158, 214], the conformation-selective binding of intrinsically disordered calcineurin to calmodulin [56], and the association of BRCA1-BRCT peptides [215].

##### 5.4.1.2 Directly Coupling MD and BD via Milestoning

A robust alternative for directly coupling BD with MD simulations is through milestoning theory, as implemented in the SEEKR framework [131, 216, 217]. In this approach, MD provides high-resolution details of binding near the target site (the “post-encounter” region), while BD calculates the diffusional flux to the transient encounter boundary. The macroscopic on-rate is then estimated by combining the MD-derived probability that a solute commits to binding (*P*_*react*_) with the BD-derived diffusion rate to the boundary 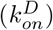:

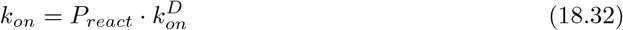

This framework was further advanced in subsequent code releases [131, 217].

### 5.5 Multiple Binding Events & Systems: BD Treatments

We now discuss the application of BD in the context of multiple, sequential binding events, first within a single complex and then involving multiple complexes.

#### 5.5.1 Allostery for a Single Complex

We assume that once a substrate arrives at the transient-encounter hypersurface, its fate is to either diffuse away or transition to its final binding mode. In this position, short-range non-bonded interactions between the substrate and target enzyme are optimized, oftentimes through structural rearrangements of the enzyme and the substrate. Although many aspects of this process are reviewed extensively in the literature [218, 219], we briefly discuss two phenomena intrinsic to post-encounter binding in the context of the transient encounter: allostery and cooperativity.

While allostery and cooperativity are conventionally associated with systems in thermodynamic equilibrium, they can also significantly influence binding kinetics. Prominent examples of allosteric behavior that could impact kinetics can be found among members of the GPCR superfamily, as their intracellular G protein binding regions are usually unavailable until the cognate ligand binds the receptors’ extracellular domains. The rapidity with which the structural changes at the extracellular side propagate to the G protein binding sites, relative to the residence time (Eq. 18.30) of a cognate ligand, can influence which G-protein mediated downstream signaling pathway is activated [220].

Extending on the earlier allosteric example, the pre-assembly of a GPCR with its cognate G protein can in turn affect the affinity for its endogenous ligand [221]. This is an example of “scaffolding”, whereby target proteins are components of larger macromolecular assemblies that tune substrate affinities and diffusional searches. In cardiac myofilaments, for instance, troponin C’s calcium binding kinetics are significantly altered when it is assembled into the larger troponin complex, and again when assembled with other components of the thin filament [222]. From an equilibrium perspective, a scaffold could impact affinity by constraining the available conformational states of a protein, disfavoring a low-affinity state, and thereby increasing the relative population of the high-affinity state. From a kinetic perspective, the precise mechanism is less clear. Ancillary units, for instance, could alter the energy barrier for substrate binding, perhaps by destabilizing the unbound state, which would increase the rate constant according to Kramers’ theory (Eq. 18.20). Alternatively, if ligand binding is followed by a subsequent binding event elsewhere, the binding of one component could shift the population distribution [223] and thereby decreasing the dissociation of the first ligand, per Eq. 18.26.

##### 5.5.1.1 Computational and Analytic Approaches

Cooperative binding thermodynamics are often described by Hill equations of the form

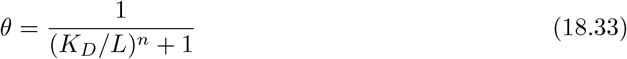

where *θ* is the fraction of protein binding sites occupied by ligands, and [*L*] and *K*_*D*_ are the ligand concentration and dissociation constant, respectively. By fitting the Hill equation to experimentally measured ligand-protein binding curves, the Hill coefficient *n* can be estimated, for which positive cooperativity is reflected for *n >* 1 and negative cooperativity for *n <* 1.

Another popular analytical model for studying cooperativity and allostery is the Monod–Wyman–Changeux (MWC) model [224]. This model assumes that allosteric proteins are oligomers comprising identical subunits, akin to a scaffold, with each subunit able to exist in at least two conformational states. A small ligand can bind differentially to either of those defined states, upon which the equilibrium population of the subunit’s conformational states can redistribute to favor those with higher ligand affinity. A key assumption of the MWC model is that all subunits must undergo concerted shifts to alternative conformation states. Recent applications of the MWC model include explaining the positive cooperative binding between allosteric and orthosteric ligands in GPCRs [225].

Beyond these analytic models, MD simulation provides a numerical approach for revealing allosteric mechanisms in proteins [226]. The primary advantage of MD for deciphering allostery and cooperative binding is its ability to provide atomistic descriptions of conformational dynamics upon ligand binding, including those potentially arising from allosteric or orthosteric sites. Furthermore, MD can uncover communication pathways connecting these multiple sites, thereby revealing hidden intermediate states whose stability is sensitive to ligand binding [226]. Where MD generally falls short is in the sampling of a sufficient number of binding and unbinding events to capture transitions between markedly different conformational states. For this reason, while Brownian dynamics (BD) is not traditionally used for insights beyond the initial diffusional encounter, it holds immediate potential for predicting binding kinetics to hidden allosteric sites that are unveiled by a bound ligand. In this capacity, BD could serve as a cornerstone for estimating *K*_*D*_ for analytic models like Eq. 18.33.

#### 5.5.2 Signaling Cascades and Substrate Channeling

Cellular responses to stimuli often involve sequential reactions and dynamic enzyme assemblies, which can be modeled computationally. Cells typically respond to extracellular stimuli using a variety of messengers, including ions, ligands, small proteins, or lipids, such as in the binding of an extracellular ligand to a plasma membrane receptor, culminating in downstream responses like gene transcription (see Figure 1, top). These responses to stimuli can enlist a series of sequential reactions, wherein a reactant A is converted to B, reactant B is converted to product C, and so forth. In many reaction networks, the responses are subject to negative or positive feedback regulation, or for consensus building among multiple signals [227, 228]. Under certain conditions, this network topology can give rise to oscillatory changes in substrate concentrations [229].

The kinetics of sequential reactions are shaped by intrinsic catalytic properties of an enzyme, its configuration within macromolecular assemblies (e.g., scaffolds), and properties of its surrounding environment [230]. In some cases, the efficiency of sequential reactions are controlled by substrate channeling (or shuttling), whereby coupled reactions occur within the same enzyme or via dynamically assembled enzyme complexes (such as in Figure 6). Substrate shuttling can arise when a substrate is localized to an active site on a target, such as via electrostatic steering [51, 231] effects that can accelerate *k*_*on*_. In tandem, steric confinement of a substrate can prevent its diffusion away from the binding site and in effect, increase the probability of reaction, as was elegantly demonstrated for amino acids and quinones in the PutA peripheral membrane flavoenzyme [232]. While this confinement tends to decrease the apparent *k*_*on*_[3], it can improve the efficiency of reactions by decreasing competition with off-target substrates.

In the context of multienzyme complexes, the proximity of enzymes, often coupled with favorable electrostatic interactions between the enzymes and reaction intermediates, prevents significant diffusion of a given intermediate away from the reactive centers [233], thereby accelerating overall reaction rates [234]. However, substrate channeling does not always entail enhancing diffusion-limited rates, as the reaction may still be reaction-limited [235]. In such cases, channeling prevents reactive intermediates from diffusing from the reaction complex or be consumed by competing enzymes [236, 237]. This selectivity can be tuned by controlling the distances between reactant sites and the strengths of electrostatic interactions. For instance, enzymes can dynamically assemble to mediate metabolic pathways, such as for glycolysis and oxidative phosphorylation [235], thereby controlling metabolite flux at branch points.

Prime examples of substrate channeling are found among the many enzymes involved in cellular metabolism. For the fatty acid synthase (FAS) as one example, substrate channeling occurs between multiple enzymatic domains, via its Acyl Carrier Protein (ACP) domain that covalently binds intermediates and shuttles its cargo [22] (Figure 2a). Similarly, the mitochondrial electron transport chain is another prime example, for which quinone molecules shuttle electrons between large, membrane-embedded complexes [12, 13]. Additional examples include DHFR [233] and glycolysis enzymes [238].

##### 5.5.2.1 Computational Approaches

###### 5.5.2.1.1 Kinetic Monte Carlo and Agent-based Models

Kinetic Monte Carlo (KMC) is a stochastic simulation method that has been effectively applied to simulate substrate channeling efficiency. For example, this approach was utilized to probe the conversion of glucose to 6-phosphogluconolactone by hexokinase and glucose-6-phosphate dehydrogenase. The study modeled an engineered system where the two enzymes were tethered by a charged linker, which served as a physical track guiding substrate transfer [239].This algorithm evaluates the total escape rate from a given state *i* as *k*_*tot*_ = ∑_*m*_ *k*_*i,m*_, where *k*_*i,m*_ is the transition rate from state *i* to any accessible state *m*. The simulation proceeds by drawing a random number *r* ∈ (0, 1] to select the next state *j*^*′*^, provided it satisfies the condition:

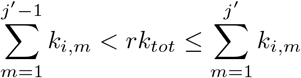

Following the transition, the current simulation time is advanced in time by increments drawn from an exponential distribution with a mean of 1*/k*_*tot*_ [240]. This procedure is repeated for the duration of the simulation.

Agent-based models (ABMs), which model the states of individual molecules (agents) as they react with their environment and with other agents, have also been used to study metabolic channeling in virtual cellular environments [241]. In a study by Klann et al., the conversion of glucose to lactate through four sequential enzymatic reactions was simulated using an ABM [241]. The model considered different spatial arrangements of the four enzymes, either in a linear complex or as a randomly distributed complex, and described substrate dynamics using predetermined effective diffusion constants and reaction rates with individual enzymes. Using this ABM framework, they demonstrated that the final product formation rate is influenced by the enzymes’ spatial distribution, which can be optimized to maximize the production rate [241].

###### 5.5.2.1.1.1 PDE-based Frameworks

The strategies described in this review so far largely apply to particle-centric systems. However, considering ensembles of particles permits the use of continuous representations, namely partial differential equations, to evaluate kinetic parameters. We therefore introduce reaction–diffusion equations (Eq. 18.35), which are continuous, PDE-based representations that are commonly used to model biomolecular events occurring over larger spatial scales. A reaction-diffusion system can be used in detailed environments consisting of explicit protein geometries or long-range interactions that would otherwise be intractable to solve analytically. A reaction diffusion system for two species, A and *A* · *B*, that undergo the equilibrium relationship in Eq. 18.1 can be expressed as:

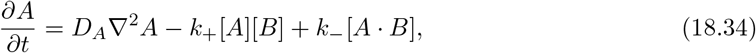

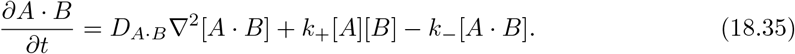

As with Eq. 18.16, the *k*_*on*_ of species *A* binding to its target (*B*) can be determined by solving for the steady-state of these expressions, then using the steady-state probability to compute the flux along a reactive boundary via Eq. 18.19 to yield the association rate. Eq. 18.34 of course can be extended to incorporate mean-field interactions and in the event that the protein target B and its complex with A are stationary, the diffusion term for 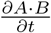 can be dropped. Such a framework has been applied to understand how the alignment of reactive centers and characteristics of associated electrostatic potentials can drive shuttling rates and efficiencies for charged substrates [242].

###### 5.5.2.1.1.2 BD Simulations of Sequential Substrate Channeling

BD has also been used to study substrate channeling and substrate transfer in parallel with experimental approaches. BD models substrate channeling within dynamic enzyme assemblies by simulating the diffusion of an intermediate product generated at the active site of one enzyme to its reactive site on a target enzyme. This approach allows for the calculation of transfer efficiency as a function of inter-enzyme distance and electrostatic steering, as discussed for PDE-based frameworks. For example, horseradish peroxidase (HRP) and glucose oxidase (GOx) form an enzyme pair, for which the product of the first (H_2_O_2_) serves as the substrate for the second. By immobilizing these enzymes on an artificial scaffold like DNA origami, the impact of inter-enzyme separation on transfer rates can be measured experimentally [243], and modeled by BD [197].

BD simulations have also been applied to other substrate channeling phenomena. For the bi-functional dihydrofolate reductase-thymidylate synthase (DHFR-TS) system of P. falciparum, as an example, the negatively-charged dihydrofolate molecule synthesized at the TS site diffuses to the DHFR site via electrostatic steering. BD simulations of this process revealed the efficiency afforded through the inter-domain substrate transfer and successfully identified key charged residues comprising an electrostatic channel [233, 244]. Additionally, BD has been used to model substrate channeling in a metabolon formed by malate dehydrogenase (MDH) and citrate synthase (CS) in the TCA cycle. These simulations estimated the transfer efficiency of oxaloacetate (OAA) from MDH to CS in the presence of electrostatic steering [238].

### 5.6 Limitations of BD Approaches

Several limitations continue to hinder the robust utilization of simulations for predicting binding kinetics. The kinetics of enzymatic processes represent a fundamentally more complex problem than the prediction of static binding poses and equilibrium affinities. Enzymatic reactions that occur in vivo involve multiple substrates, essential cofactors, and a sequence of elementary steps, making it significantly more difficult to measure experimentally and computationally predict overall reaction kinetics [245]. This difficulty is compounded by the computational expense of detailed kinetic simulations, especially when solvents are treated explicitly. In addition to computational expense, accuracy can be limited by the quality of the force field parameters used, particularly for transition metals and nonstandard ligands [246, 247]. For these reasons, computational studies are often biased toward small molecule-protein binding events, given that there are more experimental kinetic data available compared to protein-protein or protein-peptide interactions.

In addition, one of the key limitations of BD is that explicit solvent effects can be difficult to capture, especially close to the solute. For instance, implicit solvent models do not capture the discrete hydration shells and water-mediated hydrogen bond networks that can be critical determinants in final stages of molecular recognition and binding pocket desolvation[248]. Creating accurate implicit solvent models that can capture long spatial and temporal correlations could benefit from novel network architectures, such as graph neural networks, to learn these behaviors from explicit solvent data [249].

## 6 The Next Frontier

### 6.1 Introduction

Sections 3– 5 demonstrated how binding kinetics for simple, dilute solutes can be determined using analytic, numerical, and computational methods. However, predictions based on these idealized in vitro conditions are unlikely to translate directly to in vivo contexts, as they neglect the complexity of the cellular environment. The cellular environment is dynamic and crowded, within which metabolic fluxes, spatial gradients, and local chemical conditions can shape when and how binding events occur. Because simulating this level of complexity is impossible with any single computational method, multi-resolution strategies are likely necessary to approach this frontier. We propose that Brownian Dynamics can serve as a central, unifying interface for connecting models across these disparate scales (Figure 8).

We proceed by outlining several research areas essential for modeling the in vivo cellular environment, beginning with processes that involve sequential binding events (Section 6.1.1). In Figure 8, we depict prominent factors, from individual molecule encounters to macroscopic cellular compartmentalization, that can collectively influence in vivo kinetics. These factors are relevant over distinct temporal and spatial scales, necessitating a multi-scale computational framework for a complete description of binding events in vivo. By integrating multi-event modeling with experimental data collected at multiple scales, emergent biological insights are possible.

The complex binding events occurring in vivo can be categorized into three regimes based on spatial scale. 1) Single binding events: The focus here is on transient encounters, determining how quickly a binding event occurs and how stable the resulting complex is. At this scale, factors influencing binding rates, such as pH, ionic strength, and post-translational modifications, as well as post-encounter effects such as conformational flexibility, can be addressed by integrating QM, MD, and BD techniques (Figure 8). 2) Multiple binding events: For processes like metabolic channeling, BD is especially suitable. Its strength in simulating diffusion-controlled encounters makes it ideal for modeling sequences of binding events where the product of one reaction becomes the substrate for the next. 3) Cellular scale: At this level, macromolecular crowding and cellular compartmentalization create confined functional domains, whose reduced effective volumes can significantly alter the thermodynamics and kinetics of binding events relative to bulk solutions. Homogenization techniques and partial differential equation (PDE)-based models, are highly effective for capturing these phenomena.

These three scales are integrated in Section 6.3 (Figure 9), where we discuss multiscale modeling for connecting biological processes across spatial and temporal scales to capture emergent system behaviors. The section closes with a discussion on leveraging artificial intelligence (AI) techniques to enhance BD-based modeling (Section 6.4) and highlights representative applications in critical areas like drug design (Section 6.5).

#### 6.1.1 The Challenge of in vivo: Cellular Environments

Enzyme kinetics are frequently studied in isolated, well-controlled in vitro systems. Analogously, most computational applications assume simplified conditions for interacting targets and substrates exist relative to the cellular environments in which they normally occur. In contrast, the cellular environment is enriched with complex and interdependent processes, inclusive of competing substrates, allosteric modulators, and nonspecific interactions. Moreover, it is an important point to recognize that cells operate far from equilibrium, rendering the kinetics of a process just as important as its thermodynamics. For instance, variations in membrane potential, ionic composition, pH, and macromolecular crowding can all alter the rate of transient encounter complex formation and the composition of the resulting ensemble [250]. Further influencing these factors are covalent, post-translational modifications like oxidation, protonation, and phosphorylation, all of which can strongly affect the stability of post-encounter binding configurations [235]. Accounting for these complex in vivo effects is challenging, limiting insight into enzyme function within the cellular environment. We therefore discuss the role that BD could play in accommodating the impacts of these effects on enzyme kinetics.

#### 6.1.2 The Need for Bidirectional, Self-consistent Multiscale Models with Shared Figures of Merit

Simulating substrate/target binding kinetics in vivo requires a blend of mathematical and computational frameworks for phenomena intrinsic to different spatial and temporal scales . Existing modeling approaches are already positioned to integrate factors including macromolecular crowding, subcellular compartmentalization, metabolite channeling, and mechanical forces, albeit usually within a single length or timescale or in some cases, by successively passing information from microscopic (e.g. atomistic) to macroscopic (cells, tissue) scales. We argue that the next leap in predictive power requires a paradigm shift away from simple one-way upscaling towards frameworks that enforce self-consistency between scales and enable information to flow bidirectionally between scales (Figure 8). By treating molecular, macromolecular, and cellular processes as coupled, we can begin to understand how small perturbations within one scale propagate to others, and can sometimes manifest emergent behaviors, such as metabolic oscillations [251]. Bidirectional, self-consistent modeling while conceptually simple brings about significant challenges. These challenges include determining the important scales relevant to a given in vivo biological process and establishing robust biological parameters, or figures of merit, predicted from each coupled scale. Establishing appropriate figures of merit and requiring their self-consistency between coupled scales is a likely antidote to runaway propagation of error that can plague multiscale approaches. We propose that BD can play a central, bridging role for linking the scales and biological information deemed relevant (Figure 9).

### 6.2 In vivo Modeling in Stages

#### 6.2.1 Transient Encounter in vivo

##### 6.2.1.1 Cellular Changes Impacting the Transient Encounter

As discussed previously, a core assumption in Brownian dynamics simulations is that macroscopic factors like ionic strength and pH are constant. However, cells are dynamic, where fluctuating metabolic demands or even cell phenotype can impact the propensity for biochemical reactions to occur. For example, the transient encounter rate between interacting species can be affected by the extent of macromolecular crowding, impacting their collision probability; this is particularly evident when reactants are sequestered to specific cellular compartments or phases, such as lipid rafts in the plasma membrane. Such localization effects can greatly impact apparent association kinetics.

Electrostatic interactions and conformational changes influence binding kinetics. Comparing Eq. 18.15 and Eq. 18.18 reveals that electrostatic attraction can accelerate binding kinetics beyond the basal rate by modifying the reaction potential energy surface (PES) (Figure 3). For instance, a pH change altering a protein’s protonation state can shift its net charge, thereby modulating the binding of charged ligands over significant distances. Similarly, covalent post-translational modifications like phosphorylation can increase a protein’s net negative charge and steric interactions. Charge changes due to these modifications can have particularly strong effects on the structure of disordered regions in proteins, depending on the intrinsic charge of the site’s neighboring residues [16], which in turn impact binding site availability. Langevin (flexible solutes) and Brownian (rigid solutes) dynamics are well-suited to treat long-range electrostatic interactions between a target and its substrate; by combining these simulation tools for instance with PDEs that account for how dynamic changes in pH and surface charge alter the electrostatic environment, a more holistic model of binding kinetics is possible [252].

The study of diffusion-limited reactions in complex biological environments, particularly those involving tethered substrates or targets, requires a nuanced approach. To illustrate this, we provide a representative example: the diffusion of a tethered, intrinsically disordered region of the myofilament protein Troponin I, toward a hydrophobic cleft in its target protein Troponin C that is unveiled when the latter binds calcium (Figure 2f) [253]. Typically, studies have modeled tethered components in idealized environments that neglect macromolecular crowding and associated electrostatic effects; accounting for those factors may yield differences in protein disorder relative to idealized simulations that could significantly alter binding phenomena (Figure 2e). Even in the cardiac sarcomere, this TnI/TnC example is just one of many protein/protein interactions involving intrinsically disordered regions and tethers (reviewed in [254]), for which crowding effects could have significant impacts on their association kinetics [184]. Towards capturing these effects, recent frameworks have simulated tethered components either explicitly, using flexible linkers bridging folded domains, or implicitly, by analytically estimating the local concentration of tethered binding elements via polymer models [255].

#### 6.2.2 Post-encounter in the Context of Cell State

Just as the cellular environment influences the initial diffusional encounter, it can also profoundly affect the subsequent transition to the final bound state. Cellular signaling pathways, for instance, can trigger PTMs such as phosphorylation, acetylation, and glycosylation that in turn may impact the rate and mechanism of bound-state formation, through altering a protein’s charge distribution, hydrophobicity, or conformational preferences [256, 257]. These shifts reshape the local energy landscape of the encounter, modifying the stability of transient intermediates and transition states associated with the final bound state. In addition, the physical properties of the local environment, like membrane surface charges and spatial constraints in the cell, can impact how long the ligand remains near the protein following the initial encounter [258]. Together, these biochemical and structural influences bias the conformational selection process and alter the rate at which the final bound state forms; where BD simulations are used, these effects are important considerations when describing the transient encounters between diffusing substrates. Trivially, such simulations could explicitly model each protein’s PTM configuration as a distinct ensemble of conformations, though such a brute force approach would become intractable for more than a few modulated PTM sites.

#### 6.2.3 Multiple Binding Events in vivo

A holistic view of cell signaling entails considering multiple, sequential binding events. This is again exemplified in our example of the troponin proteins, where the diffusion-influenced binding of calcium, Ca^2+^, to troponin C ultimately culminates in the turnover of myosin/actin cross-bridges as the sarcomere contracts. Modeling sequential binding events presents significant challenges. The first challenge is to accurately reflect the kinetics of the individual “reactions” within the cellular environment in isolation, often while constraining their rates based on established data from in vitro or in vivo experiments [259]. A second challenge is that most studies only consider a few reactive centers in sufficient explicit (e.g. atomistic) detail due to computational or modeling limitations.

Beyond these, other key challenges include determining free parameters for states of a system that are hidden from direct measurements, such as the gating of ion channels. Often, several kinetic or even structural models are compatible with observed data, that is, they are not unique and can permit a wide range of parameter values [260]. Genetic algorithms and Bayesian inference [261, 262] have been used with success to identify and parameterize best-fit models to collected data. In addition, there may be thermodynamic or kinetic constraints on steps in concerted binding events, such as the free energy of ATP hydrolysis that sets the catalytic activity of ATPases [263–265]. Toward this end, parameter lumping, inspired by Crampin and others, can reduce the total number of states in the system and thereby reduce computational expenses of modeling by merging those steps undergoing rapid equilibrium into macro states that co-evolve with comparatively slower states [263, 264]. Other constraints include those for ensuring that compartmented or spatially-explicit models yield spatially-averaged values that are temporally consistent with whole-cell measurements [266, 267].

Generally, a compromise must be made between the accuracy of representing proteins (e.g., the detail of the molecular surface or its conformational dynamics) and the number of reactive centers. This trade-off is apparent even when modeling systems comprising only a few reactive centers, such as in the assemblies of oxidative phosphorylation complexes portrayed in Figure 2A. In this complex, the kinetics of the respective complexes’ dynamics, variations in their relative positions, and the electro-diffusion of the many reactants and products yield a staggering number of possible configurations, all of which can individually influence reaction kinetics and efficiency. Here, BD approaches offer computational savings by modeling a single substrate’s reaction with targets while assuming all others are in steady state. This approach is similar to how continuum reaction-diffusion models handle multiple species, in essence by assuming steady-state conditions for all reactions represented by the system of equations. Relatedly, when a process undergoes frequent binding events in time or in space, they can be amenable to reduced representations such as ordinary differential equations (ODEs) to follow their probabilities instead of tracking individual events. Such reduced representations are useful for describing complex biological circuits involving signal amplification or feedback (Figure 2d).

It is also important to reflect the contributions of cellular compartments and different phases that are in proximity. Complexes like the electron transport chain in oxidative phosphorylation (Figure 2a) or chromatin complexes (Figure 7) for instance involve substrates that can partition into different phases. This is further exemplified in the aggregation of LIM proteins undergoing liquid-liquid phase separation (LLPS) [199]. In some cases, the individual phases can be modeled as independent, homogeneous systems, but their coupling still requires careful consideration of interfacial terms that could impact solute partitioning. Similarly, cells are rife with processes involving substrates that must traverse interfaces like lipid bilayers, such as material transport through aquaporins and nuclear pore complexes.

### 6.3 Multiscale Modeling of in vivo Systems

In the preceding section, we highlighted a number of complications that arise in simulating reactions in vivo. Here we highlight key scales that may be considered for the multiscale modeling of a typical in vivo binding event, and how BD may be one tool to link those scales to interpret and predict binding kinetics. At the macroscopic level of cell signaling, continuum ODE or PDE models can be used to describe reacting substrates by their concentrations, either across an entire cell or within specific subcellular regions. At the intermediate scale, coarse-grained models can be used to model a substrate’s diffusional encounter at a single or small number of reactive centers. At the microscopic scale, molecular dynamics simulations can be used to reflect important conformational motions the substrate and target undergo during a successful binding event. At the finest spatial scale relevant for typical biological processes, the catalytic conversion of a substrate into a product is a quantum mechanical phenomenon involving the rearrangement of electrons and nuclei. Therefore, a single in vivo binding event encompasses a vast range of scales, each governed by distinct physical considerations.

Computationally bridging these distinct physical scales is key to modeling in vivo biological events. Although robust methods exist for individual scales, developing computationally-efficient frameworks to connect them remains a central challenge in the modeling community. This process involves defining appropriate fine- and coarse-grained models for the scales deemed important and establishing the rules for how information, such as parameter or state values, is passed between them. The reaction–diffusion equation in Eq. 18.16 provides an excellent illustration of a multi-scale modeling approach for coupling microscopic and macroscopic scales. Namely, the equation captures the continuum level diffusion of a given substrate, subject to its concentration gradient (∇^2^*p*) and in response to a potential of mean force (∇*U*), of which the latter could be derived from MD, BD or other particle-based modeling techniques.

We go beyond the reaction-diffusion description of Eq. 18.16 and propose BD as a simulation paradigm uniquely well-suited to broaden multiscale modeling, by serving as the mesoscopic bridge between microscopic, atomistic-resolution systems and macroscopic, continuum-level descriptions. Already, we have discussed how BD is routinely interfaced with molecular dynamics simulations, much in the same way quantum mechanics modeling has long been combined with molecular dynamics simulations (QM/MM) to model enzyme chemistry [268]. Yet, at the other end of the spectrum, where macroscale cellular processes are commonly described using reaction-diffusion systems, there is an excellent opportunity to make use of BD methodologies, but strategies to do so are comparatively less developed. This gives rise to a key frontier in furthering multiscale modeling: linking particle-based MD and BD simulations with continuum models, which we believe is necessary for quantitative probes of cellular processes in molecular level detail. Therefore, we will next illustrate how BD can integrate modeling frameworks that operate at different spatial and temporal scales, by highlighting successful multiscale approaches that are amenable to further augmentation with BD simulations..

#### 6.3.1 Exchanging Microscopic Information with BD

##### 6.3.1.1 QM:(MM/MD)

As a prototypical example of multiscale modeling, the QM/MM approach couples ab initio, quantum chemical models of a molecule’s electronic structure with classical particle-based molecular dynamics simulations. When applied to substrate/enzyme modeling, it can provide ground states and transition states at stationary points along a reaction pathway and their corresponding zero-point energies [268]. The approach is based on using a user-defined molecular boundary that serves as an interface between the catalytic site where QM is applied and the classical region, across which the atoms are coupled (see Figure 9). Technically, this is done by iteratively optimizing a wavefunction in the quantum domain: first, a trial function is computed for a set of nuclear coordinates, and then the forces acting on the atoms are used to advance the atomic positions in both the classical molecular mechanics/dynamics and QM domains until figures of merit like electronic energies and forces converge. Popular tools that use this method include ONIOM [269] and the generalized hybrid orbital (GHO) method [270]. While useful for predicting reaction pathways and providing estimates of reaction rates [271], QM/MM simulations are generally limited to timescales up to tens of picoseconds, which are only a fraction of the millisecond and longer time scales required for large conformational motions in many enzymes.

BD simulations are generally not directly interfaced with ab initio calculations, though force field parameters and partial charges used for the simulations have long been derived from QM [272]. As a step toward BD integration, one could leverage the strategy used for polarizable continuum models (PCM) that interface a QM-treated solute with a continuum solvent [273]. PCM calculations rely on defining a solute/solvent interface as a boundary for continuum solvation modeling, based on the electron density predicted from ab initio calculations. Similarly, BD simulations performed in the implicit solvent surrounding a target could provide substrate/target binding poses as starting points for the QM treatments. Those poses could in turn be used to generate updated electron densities as inputs for the electrostatic potential used for BD. The advantage of BD in this context is its rapid sampling of transient-encounter complexes, whose conformations can be refined through more detailed MM and QM protocols. Convergence of the two models could be based on when the BD-predicted substrate poses adequately saturate the target surface that is used for the QM based electrostatic potentials.

##### 6.3.1.2 MD:BD

Conventional all-atom molecular dynamics (MD) simulations naturally provide access to much longer timescales relative to QM and QM/MM approaches, with microsecond length simulations now routine [246]. Although MD simulations have been used in tandem with Brownian dynamics (BD) in numerous applications, they have conventionally been linked in an ad hoc manner, usually by identifying starting conformations from MD to be used with rigid-body BD simulations [56, 274]. The recently proposed SEEKR approach moves beyond this convention by using a systematic solution that directly couples BD to MD. [217]. Analogous to the QM/MM example, it is based on defining a boundary surface dividing the two simulation domains representing bound and fully-dissociated states. Near the bound configuration, ligand-enzyme binding is treated using all-atom, explicit solvent MD; beyond this region BD is used. A key limitation of the SEEKR approach is that the MD/BD boundary is defined a priori, often based on geometric intuition, which may not generalize to arbitrary binding mechanism. However, an adaptive boundary approach might overcome this limitation, by continuously evaluating the ruggedness of the PMF along the reaction coordinate; a smooth potential energy surface would be expected where non-bond interactions are weak and appropriate for BD simulations, whereas roughness would signify strong intermolecular interactions better suited to MD. This boundary could be varied iteratively until a figure of merit, such as the *k*_*on*_ in Eq. 18.32 reflecting the combined MD/BD contributions, achieves convergence between iterations.

##### 6.3.1.3 CGMD:BD

Coarse-grained (CG) particle simulation approaches that use reduced representations of atom groups can sample temporal and spatial scales beyond the reach of conventional all-atom simulations. Common strategies include the Martini [275] force field for molecular dynamics and COFFDROP [276] for Brownian dynamics, In these models, groups of atoms are represented by single beads, and force fields are parameterized to recapture key atomistic-level interactions, such as those governing secondary structure. COFFDROP in particular has been used to study molecular association, with applications to IDPs including amyloid *β* and *α* synuclein [277]. Most applications of these CG approaches tend to be unidirectional, as they use atomistic-resolution data strictly as model inputs to capture coarse-grained properties like radii of gyration [277] or aggregation behavior [276].

Efforts to back-map or project CG-predicted molecular positions to their atomistic equivalents have been proposed. However, to our knowledge, prominent methods such as the Bayesian inference approach from Liu et al. [278] have not been applied to simulations of bi-molecular association. In principle, back-mapping strategies applied to COFFDROP/BD-based simulations, like the IDP aggregation events studied by Ahn et al. [277], could enable the validation of specific interactions at the atomistic level and be used to iteratively refine CG simulation parameters. Furthermore, partitioning the simulation domain into regions used for atomistic/CG refinement and others for CG-based Brownian dynamics could be amenable to the adaptive boundary approach we described in the previous section. These could be interfaced by a sharp spatial boundary, at which CGMD yields candidate substrate/target poses, whose transient encounters are modeled via BD. Fulfilling this integration would bridge the gap between the efficient sampling of CG dynamics and the structural fidelity required to describe protein-protein association. Such a development could be especially useful for proteins with extended, intrinsically disordered regions that are prone to undergoing disordered to ordered transitions during binding [279].

#### 6.3.2 Exchanging Macroscopic Information with BD

##### 6.3.2.1 (MD/BD):ODE

The examples discussed so far largely entail mapping between different resolutions of particle-based models. However, extending this coupling paradigm to real physiological systems requires interfacing microscopic particle descriptions with macroscopic phenomena described using continuum representations like ordinary differential equations (ODEs). The coupling of molecular-resolution models with ODE-based systems biology models has gained significant momentum in recent years, as exemplified by work from Silva et al. [280]. In that study, the authors bridged simulation paradigms by constructing a potential energy surface for a potassium channel using Monte Carlo sampling. To connect this thermodynamic landscape to time dependent physiology, they utilized a form of the Smoluchowski equation to compute transition rates along the PES. These rates were subsequently incorporated into a Markov model of channel gating that was integrated into a systems model of the cardiac action potential model; in effect, the strategy successfully bridged atomistic level protein function with whole cell electrophysiology. More recent applications have similarly combined MD with systems level ODEs [16, 265, 274]. For instance, Teitgen et al. [274] combined Brownian dynamics and MD to demonstrate how deoxy-ATP, an ATP analog, impacts myosin/actin association; this was in turn combined with an ODE model to predict cell contractility. Relatedly, our study of ABLIM1 [16] demonstrated how MD and CG simulations of an intrinsically disordered protein can be integrated with an ODE-based model of sarcomere contraction using polymer theory. Importantly, these multiscale pipelines replace the reliance on phenomenological parameter fitting common in systems biology with constants derived from simulation; by relying on molecular simulations to determine system-level parameters, quantitative prediction of how specific molecular perturbations, such as missense mutations or errant PTMs, propagate to alter organ-level function is possible.

A significant limitation of most ODE-based multiscale approaches linking protein-level and systems-level representations is that the scale coupling is often qualitative. For example, a faster *k*_*on*_ predicted for a given ligand/target association event may be reflected by an arbitrary increase in the corresponding rate coefficient for the ODE model, without any physical rationale for the modified value. A more robust and general strategy could be to replace the heuristic rate constants with those derived from coupled MD/BD simulations. To illustrate this, one could use PMF profiles for transient encounter complexes from MD simulations to compute intrinsic rate constants (such as *k*_off_ and *k*_int_ via Kramers’ theory), whereas BD or SEEKR could be used for estimating the diffusion-limited *k*_on_ to the transient encounter complex. This strategy was applied in a study of adenylyl cyclase, for which BD simulations were used to calculate corresponding association rates for ATP and G-proteins; those rates where then directly used in an ODE models of cAMP production [281]. The van Keulen et al. study [281] thus provides a concrete road map for the bottom-up scaling of molecular simulation-derived figures of merit, namely rate constants, to an overarching systems-level ODE model.

Less common, but equally important, are top-down constraints, e.g. macroscopic quantities derived from ODE-based systems models, that can be used to tune molecular simulations protocols. To illustrate this, suppose an ODE-based model of integrin clustering during cell adhesion [282] predicts a macroscopic rate of aggregation, *dA/dt*, at a given tension; that rate could be used as a constraint for BD models of integrin assembly [283]. BD is especially well suited as the intermediate domain for accommodating such constraints, as it can reflect diverse mesoscale phenomena including diffusion, molecular collisions, and binding probabilities, that can be inferred from macroscale figures of merit like tension, while maintaining key microscopic (atomistic) physicochemical details such as molecular self-assembly; further; the BD simulation parameters can be tuned to align its mesoscopic predictions from up-scaled microscale data with macroscopic figures of merit. This tuning generalizes existing practice, where BD/MD encounter criteria, such as steric collision thresholds or reaction probability assignments [122], are adjusted to reproduce experimentally measured rates. Analogously, the reaction probability or energy thresholds within a BD simulation could be tuned until the BD-predicted integrin association behavior matches the macroscopic *dA/dt* inferred from ODE-based tension model. Furthermore, the BD simulations could impose constraints on the association rates inferred at the macroscale, for instance, by precluding rates that exceed diffusion-limited values. By utilizing both top-down and bottom-up parameter values and constraints, self-consistent formulations of multiscale models are possible.

##### 6.3.2.2 BD:Continuum

Linking particle-based simulations (BD or MD) with continuum representations remains a challenge, though recent applications have made considerable progress toward bridging the cap. Conventional reaction-diffusion models can accommodate binding or unbinding rates derived from molecular simulations as source–sink terms or mixed boundary conditions, as was recently done by Van Keulen et al. [281] for a state-based adenyl cyclase model. More systematic bridging approaches using homogenization theory (Figure 9) with molecular-scale structural data have also been used to obtain effective macroscopic parameters like diffusion coefficients [207] or reaction rates [284] suitable for continuum modeling. We have successfully applied this framework to estimate *D*_eff_ and derive rate constants from MD for use in continuum transport problems in abiological materials [204], which can be applied to analogous processes in biological media. Additionally, strategies such as parameter lumping can consolidate rate equations for processes that equilibrate rapidly relative to the timescale of the continuum system [264], which can help align catalytic mechanisms with atomistic models [265], while reducing the problem stiffness and ill-conditioning [285] that frequently plague deterministic simulations. In parallel, physics-derived interaction potentials representing diverse phenomena, including long-range electrostatics or crowding, can be incorporated directly as mean-field potentials in expressions such as the drift term ∇*U* in the Smoluchowski equation.

BD can also serve as a dynamic interface between molecular and continuum models. Ion-channel conduction simulations are leading examples of this application, whereby the channel pore is represented with MD-derived PMFs inclusive of electrostatic effects, along which ion movements are simulated using BD/Langevin dynamics [286, 287]. Appropriate boundary conditions are chosen where the pore intersects with a continuum reservoir of ions to ensure physiological ion gradients are maintained, as summarized in [288]. More advanced hybrid BD/continuum schemes [289, 290] support bidirectional exchange by passing particle positions and continuum concentrations through a shared buffer region, ensuring conservation of both mass and momentum. However, many existing MD–BD–continuum multi-scale approaches rely strictly on static boundaries. Allowing BD–continuum interfaces to evolve dynamically by adjusting boundary placement or permeability to maintain flux continuity would greatly enhance bidirectional information flow, ensure self-consistent figures of merit, and potentially improve overall computational efficiency.

##### 6.3.2.3 The Feedback Loop: Closing the Circuit with Differentiable Multiscale Modeling

As suggested by these examples, a self-consistent multiscale modeling framework focused on the BD interface region could provide a unified strategy for linking particle-based and continuum models. The key advantage would be in establishing a true feedback loop that enables information to flow seamlessly in both directions. This would enable the backpropagation of macroscopic observables (e.g., tissue-level dynamics) down to fine-grained molecular parameters, and then propagating those updated microscopic, physicochemical details back to higher-level representations. This can be achieved by self-consistently updating shared properties (figures of merit) and dependent variables across scales, such as by enforcing continuity of chemical potential or the strict conservation of particle flux [291]. Self-consistency can be imposed by enforcing the optimization of error functions representing figure of merit mismatch, which renders multiscale modeling as a differentiable framework. An error function might take the form *ϵ* ≡ (*λ*(*ψ*) − *λ*_exp_(*ψ*))^2^, where *ψ* is an optimizable boundary coordinate and *λ*(*ψ*) is a measured property. Parameters such as the BD–continuum boundary location, energy thresholds, or effective diffusion coefficients could be optimized via gradient-based methods to minimize the error between predicted and target figures of merit. This approach is likely to work best for smoothly varying error functions, which may be unlikely given the inherent noisiness of simulation data. Techniques such as dimensionality reduction (e.g., identifying reaction coordinates that smoothly represent a process) could be helpful to mitigate the noise, although recent optimization strategies for machine learning applications, such as stochastic gradient descent (reviewed in [292]), may provide more robust solutions. With a robust error function and optimization strategy, a multiscale, differentiable model could automatically adjust parameters at the molecular scale, to reflect an emergent property like a drug’s residence time at an allosteric site, in order to minimize the difference between the simulated and experimentally-observed cellular-level response. As a whole, such a framework could provide a systematic, data-driven means to ‘close the circuit’ of multiscale model integration between ODE, PDE, BD, and MD representations.

### 6.4 AI in Enzyme Kinetics

Multiscale modeling is computationally intensive, this burden is further exacerbated by the combinatorics of having numerous degrees of freedom at molecular interfaces. Consequently, scaling these approaches to even cell-level representations via brute-force simulation alone is unlikely to succeed. In complement, machine learning (ML) offers a promising path to scalability by serving as rapid surrogate models for approximating expensive physics-based calculations when sufficient training data are available. In this section, we describe the state of the art in ML for kinetics prediction and propose potential directions for coupling ML with Brownian Dynamics (BD) to improve predictions of enzymatic kinetics.

#### 6.4.1 Prior Art

ML has revolutionized the prediction of protein structure, as exemplified by breakthroughs from RoseTTAFold [106] and AlphaFold [300] in generating accurate structural models from sequence alone. This success extends to protein design with diffusion-based models like RFdiffusion [301] and to functional interpretation using advanced protein language models, such as ESM-2 [302]. These advances provide a powerful foundation to multi-scale models, like those relying on BD to predict binding kinetics, as high-accuracy structural models could predict reasonable starting configurations needed for more detailed physics-based simulations. Surprisingly, the direct application of ML to predict the fundamental *k*_*on*_ and *k*_*off*_ rate constants of binding remains largely undeveloped especially in comparison to their well-established use for predicting biophysical properties like thermostability or binding affinity are well-established [303].

A survey of ML studies on binding kinetics from 2020 to 2025 yields just a few studies that focus almost exclusively on predicting the *k*_*off*_ rate for protein-inhibitor binding (see Table 1). The paucity of kinetic ML models is unsurprising, given the limited volume of experimental kinetic data available. Some of the most prominent datasets include 1) The Kinetic Data of Bio-molecular Interaction (*KDBI*) database, which contains experimental kinetic data for protein–protein, protein–ligand, and protein–nucleic acid interactions [304], with the latest update in 2025 incorporating RNA-protein binding kinetics [305]; 2) The *PDBbind-koff-2020* dataset, containing 680 *k*_off_ data points for protein–ligand complexes [296]; 3) A dataset of *k*_on_ and *k*_off_ values for 104 inhibitors targeting HSP90 [211]. 4) A dataset of *k*_on_ and *k*_off_ measurements for 270 inhibitors against 40 kinase targets [91] and 5) The Kinetics for Drug Discovery (*K4DD*) dataset, a resource developed through industry-academia collaboration to advance understanding of drug residence time and its relationship to in vivo efficacy [150]. These sets’ bias toward *k*_*off*_ prediction is consistent with drug residence time (1/*k*_*off*_) being recognized as a strong predictor of vivo drug efficacy [306].

**Table 1:**
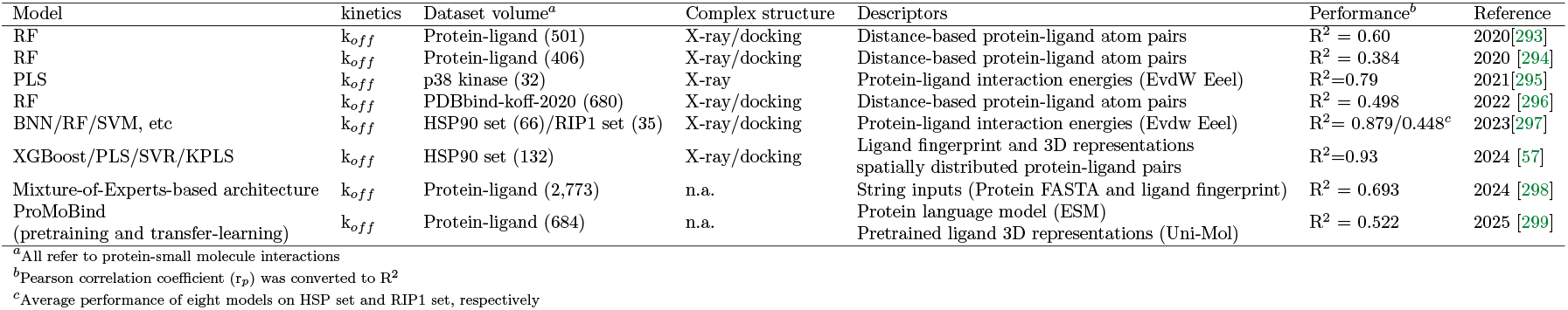
ML studies reported on binding kinetics from year 2020 to 2025. All ML models are focused on protein-small molecule dissociation rates predictions.

#### 6.4.2 Opportunities for Coupling BD and ML for Enzymatic Kinetics Predictions

The limited volume of experimental kinetic data is a major bottleneck for developing ML models to predict *k*_*on*_. This is partly because kinetics prediction is fundamentally more complex than predicting equilibrium properties, in part because a given rate constant reflects a distinct, path-dependent property rather than a simple state function. Here, BD has the potential to extend kinetic datasets used for ML model training. Specifically, BD simulations could in principle be used to generate extensive ligand/protein binding trajectories, which can then be calibrated against experimental rate data (Figure 10).

**Figure 10:**
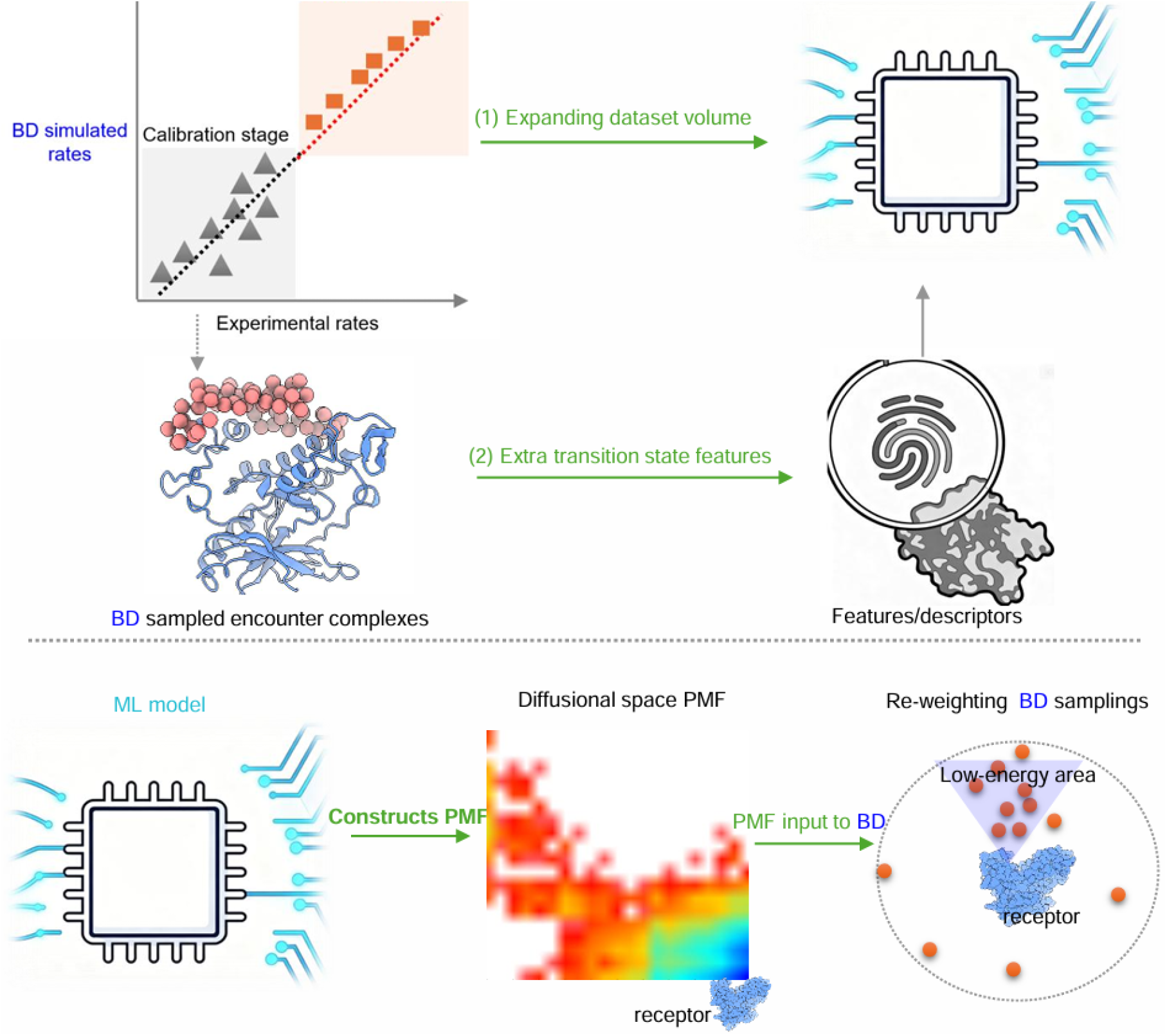
Using BD and ML for Kinetic Predictions. (a) BD informs ML training. After calibration against experimental data, BD can generate large-scale kinetic datasets for a target receptor, providing the extensive data needed for robust ML model training. Furthermore, BD-sampled encounter complexes naturally encode the path-dependent features essential for kinetic prediction, which are often elusive in static, bound-state complex structures. (b) ML enhances BD sampling. ML models can be used to generate potential of mean force (PMF) maps, such as those within the diffusional space surrounding a receptor. These ML-derived PMFs can serve as direct inputs for BD simulations, significantly accelerating sampling efficiency. This approach enables BD to model more realistic, atomically detailed biological binding events, thereby alleviating the inherent sampling burden required for accurate kinetic predictions.

The selection of appropriate features or descriptors is critical for the strong performance of ML models in predicting kinetics. Theoretically, binding kinetics (*k*_*on*_ or *k*_*off*_) are determined by both the transition state and the unbound/bound states. Therefore, ML features designed to predict these rates must be informed by features derived from both states. Most current ML studies on *k*_*off*_ prediction use features based heavily on bound-state complexes, such as ligand-atom pair distances [293, 296] or interaction energies [297]. Inherent in these strategies is an overreliance on bound-state feature to learn *k*_*off*_trends, when there are reports that *k*_*on*_values actually correlate more strongly with than *k*_*off*_values, at least for kinome-inhibitor complexes [91]. This raises the significant question of whether bound-state descriptors alone are truly sufficient. A major challenge is that descriptors often under-represent the role of protein dynamics and additional path-dependence. Although such features can be extracted from MD trajectories sampling binding/unbinding events, this is a computationally expensive strategy unsuitable for high-throughput tasks. BD can reduce this limitation by efficiently simulating thousands of substrate-enzyme encounter complexes. Meanwhile, outputs from BD, such as electrostatic potential maps or encounter complex statistics, could serve as valuable input features for ML model training (Figure 10).

In turn, there are burgeoning opportunities for ML to directly enhance BD simulations. ML methods can be used to directly improve BD-derived rate estimates or to enhance the underlying simulation data driving them. One critical area is the determination of PMFs characterizing substrate-target association pathways, which serve as essential inputs for BD. ML methods are increasingly integrated with PMF techniques to improve the accuracy of those derived from MD simulations [307], and could in principle be used to estimate the diffusion coefficient along a PMF surface. These refined PMFs can help prioritize exploration of under-sampled regions of diffusional space, such as by optimizing reaction coordinates for sampling or reconstructing PMFs in poorly-sampled regions using autoencoders [308] . Each of these contributions would be expected to greatly accelerate and improve the baseline accuracy of BD simulations (Figure 10).

#### 6.4.3 Challenges in ML+BD Approaches

As stated earlier, the development of robust ML models for binding kinetics is significantly hindered by the scarcity of large, high-quality, and standardized experimental datasets for kinetic rate constants (*k*_*on*_ and *k*_*off*_). Moreover, existing kinetic data are measured under different experimental conditions, which complicates the curation of consistent and reliable training sets. This scarcity of high-quality and consistent data exacerbates issues regarding model validation, and in particular, information leakage. This leakage arises when models are trained and tested on nonindependent datasets. Without rigorous splitting of the training data, models may simply memorize sequence patterns that are absent in unseen targets rather than learning generalizable principles [309]. Consequently, high-quality datasets are essential for implementing homology-aware splitting, where training and testing sets are systematically divided to ensure no close evolutionary relationship exists between them. This ensures the model ascertains transferable physicochemical rules rather than dataset-specific biases. Therefore, careful curation of input data sets and continuous retraining as more experimental data are accrued will be necessary to mitigate this risk.

Beyond data availability, predicting *k*_*on*_ is intrinsically difficult due to the complexity of the underlying physics. These difficulties include modeling implicit solvent effects and capturing long-range spatial and temporal correlations. Addressing these physical complexities often benefits from the use of advanced network architectures, such as 3D Convolutional Neural Networks (CNNs), which are capable of identifying patterns in complex spatial data. Such CNN approaches typically partition the protein interaction space into voxels containing physicochemical properties like electrostatic potentials and hydrophobic character; in turn, this discretization allows the network to identify long-range correlations and geometric patterns driving association [310].

### 6.5 Applications

We next outline potential applications of multiscale modeling utilizing BD as a central interface.

#### 6.5.1 Drug Design and Systems Pharmacology

In modern drug discovery, binding kinetics are recognized as critical determinants of a drug’s potency and duration of action [311]. For instance, it is increasingly recognized that the residence time of the drug (see Eq. 18.30) when bound to its target is a critical determinant of its overall pharmacological efficacy in a dynamic, open physiological system [66], hence motivating sustained target inhibition as a key consideration in drug development [312] For example, a long residence time (slow *k*_*off*_ rates) can be more important for in vivo efficacy than binding affinity in many kinase inhibitors [91], such as those targeting the PI3K pathway [311].

Once the binding behavior of a candidate molecule is well-characterized, multiscale modeling could be helpful in optimizing its systems-level efficacy. Simulations that account for the complex in vivo environment may help define critical therapeutic windows, thereby placing explicit constraints on the necessary *k*_*on*_ and *k*_*off*_ values for efficacious treatment. Because many drug-target association events are reaction-limited rather than diffusion-limited, the binding process is often governed by conformational selection, induced-fit mechanisms, or rebinding rates [313]. Consequently, computational models may be beneficial in such cases for identifying compounds that target specific, transient functional states, such as intermediate states of ion channels, as opposed to simply optimizing affinity for a single static structure. For instance, the Clancy group has modeled ion channel gating kinetics to identify drugs that specifically target intermediate states of the ryanodine receptor (RyR) to correct arrhythmias [314]. Here, BD is particularly well-suited to resolve the kinetics of diffusion and binding to such transient or cryptic sites, enabling the design of pathway-specific drugs that finely tune residence times and therapeutic efficacy [315]. Building on this, a more sophisticated drug design strategy could involve stabilizing specific functional conformations through allosteric trapping. For example, we recently used Bayesian inference to infer calcium uptake rates by the SERCA pump [260]; this framework could be combined with BD simulations to model how certain endogenous peptide modulators prefer the pump’s E1 state while others target its E2 state, thereby directly impacting Ca^2+^ pumping rates. Overall, a differentiable, multiscale approach in this context could go far beyond optimizing a single parameter like affinity by allowing for the simultaneous co-optimization of multiple, interconnected properties under appropriate biological constraints.

#### 6.5.2 Peptide Design for Dynamic Processes

A BD-centric framework for integrating molecular-level alterations with cellular-level outcomes could also be invaluable for probing and modulating signaling pathways. To illustrate this, we refer to a modified HEK 293 cell line with transient expression of the SERCA calcium pump, which has been used to characterize the pump’s dynamic regulation by membrane-resident peptides like phospholamban (PLB) [316, 317]. For this system, experimental techniques, like FRET, can be used to assess the binding kinetics of regulatory proteins (e.g., PLB to SERCA), after which the kinetics could then be used to parameterize ODE-based models of calcium transients that are shaped by SERCA pumping activity. Brownian dynamics here offers a powerful extension by computationally predicting peptide/pump binding kinetics, for instance, by representing the PLB-SERCA interaction as a two-dimensional diffusion process within the membrane.

#### 6.5.3 Extension to Other Biological Systems

In biological systems, reaction–diffusion processes at the molecular scale often have direct mathematical analogs at longer time and larger spatial scales, such as those involving higher-order entities like macromolecular complexes or even cells. For instance, reaction diffusion models have been used to study viral capsid [318] and actin filament assembly [319], even though the ‘particles’ implicitly reflected in the models for these cases are massive, macromolecular assemblies .Similarly, since some types of isolated cells can exhibit Brownian-like motion, at the population level, collective cell migration during processes like wound healing are often described by Keller–Segel equations, which closely resemble the Smoluchowski formalism for interacting Brownian particles [320]. This analogous behavior to molecular diffusion suggests that the fundamental concepts of Brownian dynamics may translate to macromolecular assemblies and beyond, providing new opportunities to apply diffusion-based BD modeling across multiple scales of biological organization.

#### 6.5.4 A Community Effort for Model Validation and Experimental Alignment

Lastly, computational models demand rigorous validation and constraint by experimental data in order to obtain reliable predictions of biological phenomena. In this regard, there is a critical need for experimental and computational datasets that are aligned to common conditions, such as using the same crowder types and densities for both research paradigms [321]. With new, standardized data in hand, statistical frameworks such as Bayesian inference, maximum entropy, and maximum likelihood estimation can enable the integration of diverse datasets, ranging from time-resolved X-ray crystallography and NMR to FRET and stopped-flow kinetics [322, 323]. Ultimately, these frameworks provide the rigor necessary to parameterize computational models used for multiscale simulations of ligand/target binding in vivo. Given that every step of a multiscale modeling strategy demands expertly curated experimental datasets for model design and validation, a community effort for experimentalists and multiscale modelers to co-design new assays and benchmarks could be invaluable to advancing this field, much as the CASP community [324] catalyzed the eventual solving of the perennial protein folding problem.

## 7 Conclusion

In this review, we have discussed fundamental aspects of biomolecular binding kinetics, from the classical principles of mass action to stochastic phenomena. We have emphasized that the kinetic rate constants (*k*_*on*_ and *k*_*off*_) are not merely fixed parameters, but dynamic determinants of cellular function, pharmacological behavior, and therapeutic efficacy. Central to this discussion is Brownian dynamics (BD), which we identify not simply as a simulation tool for ligand/target binding, but as a potential mesoscopic bridge linking atomistic molecular dynamics with macroscopic continuum models.

We have further examined how the idealized two-stage binding process comprising the diffusional encounter followed by molecular recognition is profoundly modulated by the cellular milieu.

Factors such as macromolecular crowding, dynamic phase separation, and electrostatic steering all have the capacity to reshape the energy landscapes underlying binding events. Although hybrid BD–MD approaches have emerged for accurately capturing these multiscale pathways, forward simulation alone is unlikely to be sufficient to address the full complexity of the cellular environment.

Because the in vivo environment encompasses an overwhelming number of interacting parameters, the next frontier in kinetic modeling lies in moving beyond isolated simulations toward well-coupled multiscale models. We propose focusing on the development of differentiable frameworks featuring closed feedback loops, in which macroscopic continuum representations are used to constrain microscopic systems modeled via molecular simulations, with BD serving an essential intermediary role at the mesoscale. Such a unified approach would allow biological systems to be simulated as deeply coupled and self-consistent across modeling scales, opening new avenues for both understanding molecular mechanisms and rationally engineering complex biological processes in vivo.

## Acknowledgments

We would like to thank Audrey Deyawe Kongmeneck for her rendering of a GPCR bound to beta arrestin. Research reported in this publication was supported by the Maximizing Investigators’ Research Award (MIRA) (R35) from the National Institute of General Medical Sciences (NIGMS) of the National Institutes of Health (NIH) under grant number R35GM124977, and the National Natural Science Foundation of China (32501103). Writefull, Grammarly, Microsoft Copilot, and other computer-aided language tools were used for grammar and style editing.

## References

[1] Joel W. Schwartz et al. “Substrate Binding Stoichiometry and Kinetics of the Norepinephrine Transporter *”. Journal of Biological Chemistry 280.19 (2005), pp. 19177–19184. ISSN: 0021-9258.

[2] S. S. Rosenfeld and E. W. Taylor. “Kinetic studies of calcium binding to regulatory complexes from skeletal muscle.” Journal of Biological Chemistry 260.1 (1985), pp. 252–261. ISSN: 0021-9258.

[3] Hadi Rahmaninejad et al. “Co-localization and confinement of ecto-nucleotidases modulate extracellular adenosine nucleotide distributions”. PLOS Computational Biology 16.6 (2020), pp. 1–31.

[4] Achim K. Moesta, Xian-Yang Li, and Mark J. Smyth. “Targeting CD39 in cancer”. Nature Reviews Immunology 20.12 (2020), pp. 739–755. ISSN: 1474-1741.

[5] H. X. Zhou. “Brownian dynamics study of the influences of electrostatic interaction and diffusion on protein-protein association kinetics”. Biophysical Journal 64.6 (1993), pp. 1711–1726. ISSN: 00063495.

[6] Scott H. Northrup, Stuart A. Allison, and J. Andrew McCammon. “Brownian dynamics simulation of diffusion-influenced bimolecular reactions”. The Journal of Chemical Physics 80.4 (1984), pp. 1517–1524. ISSN: 0021-9606.

[7] Xiaodong Pang and Huan-Xiang Zhou. “Rate Constants and Mechanisms of Protein-Ligand Binding”. Annual Review of Biophysics 46.Volume 46, 2017 (2017), pp. 105–130. ISSN: 1936-1238.

[8] M. Vijayakumar et al. “Electrostatic enhancement of diffusion-controlled protein-protein association: comparison of theory and experiment on barnase and barstar”. Journal of Molecular Biology 278 (5 1998), pp. 1015–1024. ISSN: 0022-2836.

[9] H X Zhou. “Enhancement of protein-protein association rate by interaction potential: accuracy of prediction based on local Boltzmann factor”. Biophysical Journal 73 (5 1997), pp. 2441–2445.

[10] M. v. Smoluchowski. Zeitschrift für Physikalische Chemie 92U.1 (1918), pp. 129–168.

[11] P H von Hippel and O G Berg. “Facilitated Target Location in Biological Systems”. Journal of Biological Chemistry 264.2 (1989), pp. 675–678. ISSN: 0021-9258.

[12] Petri Kaurola et al. “Distribution and dynamics of quinones in the lipid bilayer mimicking the inner membrane of mitochondria”. Biochimica et Biophysica Acta (BBA) - Biomembranes 1858.9 (2016), pp. 2116–2122. ISSN: 0005-2736.

[13] Murilo Hoias Teixeira and Guilherme Menegon Arantes. “Effects of lipid composition on membrane distribution and permeability of natural quinones”. RSC Adv. 9 (29 2019), pp. 16892–16899.

[14] Eugene E Ley, Christopher E Goodyer, and Annette L Bunge. “Mathematical models of diffusion through membranes from spatially distributed sources”. Journal of Membrane Science 283.1 (2006), pp. 399–410. ISSN: 0376-7388.

[15] David Van Valen, Mikko Haataja, and Rob Phillips. “Biochemistry on a Leash: The Roles of Tether Length and Geometry in Signal Integration Proteins”. Biophysical Journal 96 (4 2009), pp. 1275–1292.

[16] Bin Sun et al. “GSK3-driven phosphorylation of ABLIM1 regulates its interactions with titin cardiac muscle”. Journal of General Physiology 157.5 (2025), e202413737. ISSN: 0022-1295.

[17] H.C. Berg and E.M. Purcell. “Physics of chemoreception”. Biophysical Journal 20.2 (1977), pp. 193–219.

[18] Andrea Strasser, Hans-Joachim Wittmann, and Roland Seifert. “Binding Kinetics and Pathways of Ligands to GPCRs”. Trends in Pharmacological Sciences 38.8 (2017), pp. 717–732. ISSN: 0165-6147.

[19] Dima Kozakov et al. “Encounter complexes and dimensionality reduction in protein–protein association”. eLife 3 (2014), e01370.

[20] David D. Boehr, Ruth Nussinov, and Peter E. Wright. “The role of dynamic conformational ensembles in biomolecular recognition”. Nature Chemical Biology 5.11 (2009), pp. 789–796. ISSN: 1552-4469.

[21] Shozo IIDA and James D. Potter. “Calcium Binding to Calmodulin. Cooperativity of the Calcium-Binding Sites”. The Journal of Biochemistry 99.6 (1986), pp. 1765–1772. ISSN: 0021-924X.

[22] Stephen W. White et al. “THE STRUCTURAL BIOLOGY OF TYPE II FATTY ACID BIOSYNTHESIS”. Annual Review of Biochemistry 74.Volume 74, 2005 (2005), pp. 791–831. ISSN: 1545-4509.

[23] Svetlana B. Tikunova et al. “Effect of Calcium-Sensitizing Mutations on Calcium Binding and Exchange with Troponin C in Increasingly Complex Biochemical Systems”. Biochemistry 49.9 (2010). PMID: 20128626, pp. 1975–1984.

[24] Byeong J. Chun et al. “Purinoreceptors and ectonucleotidases control ATP-induced calcium waveforms and calcium-dependent responses in microglia: Roles of P2 receptors and CD39 in ATP-stimulated microglia”. Frontiers in Physiology Volume 13 - 2022 (2023). ISSN: 1664-042X.

[25] Julius Rebek. “Model Studies in Molecular Recognition”. Science 235.4795 (1987), pp. 1478–1484.

[26] Claudio N. Cavasotto, Natalia S. Adler, and Maria G. Aucar. “Quantum Chemical Approaches in Structure-Based Virtual Screening and Lead Optimization”. Frontiers in Chemistry (2018).

[27] Manuela Maurer, Niels Hansen, and Chris Oostenbrink. “Comparison of free-energy methods using a tripeptide-water model system”. Journal of Computational Chemistry 39.26 (2018), pp. 2226–2242.

[28] Sergio Decherchi and Andrea Cavalli. “Thermodynamics and Kinetics of Drug-Target Binding by Molecular Simulation”. Chemical Reviews 120.23 (2020). PMID: 33006893, pp. 12788–12833.

[29] Ercheng Wang et al. “End-Point Binding Free Energy Calculation with MM/PBSA and MM/GBSA: Strategies and Applications in Drug Design”. Chemical Reviews 119.16 (2019). PMID: 31244000, pp. 9478–9508.

[30] Anita de Ruiter and Chris Oostenbrink. “Advances in the calculation of binding free energies”. Current Opinion in Structural Biology 61 (2020), pp. 207–212. ISSN: 0959-440X.

[31] Pratyush Tiwary et al. “Kinetics of protein-ligand unbinding: Predicting pathways, rates, and rate-limiting steps”. Proceedings of the National Academy of Sciences of the United States of America 112.5 (2015), E386–E391.

[32] Abraham Muñiz-Chicharro et al. “Brownian dynamics simulations of biomolecular diffusional association processes”. WIREs Computational Molecular Science 13.3 (2023), e1649.

[33] Niel M. Henriksen, Andrew T. Fenley, and Michael K. Gilson. “Computational Calorimetry: High-Precision Calculation of Host–Guest Binding Thermodynamics”. Journal of Chemical Theory and Computation 11.9 (2015). PMID: 26523125, pp. 4377–4394.

[34] Vittorio Limongelli. “Ligand binding free energy and kinetics calculation in 2020”. WIREs Computational Molecular Science 10.4 (2020), e1455.

[35] Raechell et al. “On the Relationship Between Protein Stability, Thermostability, and Allosteric Signaling”. Journal of Molecular Biology (2025), p. 169537. ISSN: 0022-2836.

[36] David M. Thal et al. “Structural insights into G-protein-coupled receptor allostery”. Nature 559.7712 (2018), pp. 45–53. ISSN: 1476-4687.

[37] Melanie I. Stefan and Nicolas Le Novere. “Cooperative Binding”. PLOS Computational Biology 9.6 (2013), pp. 1–6.

[38] Austin T. Raper, Brian A. Maxwell, and Zucai Suo. “Dynamic Processing of a Common Oxidative DNA Lesion by the First Two Enzymes of the Base Excision Repair Pathway”. Journal of Molecular Biology 433.5 (2021), p. 166811. ISSN: 0022-2836.

[39] Brian P Ziemba and Joseph J Falke. “A PKC-MARCKS-PI3K regulatory module links Ca2+ and PIP3 signals at the leading edge of polarized macrophages”. en. PLoS One 13.5 (2018), e0196678.

[40] Jingwen Bai, Yaochen Li, and Guojun Zhang. “Cell cycle regulation and anticancer drug discovery”. Cancer Biology & Medicine 14.4 (2017), pp. 348–362. ISSN: 2095-3941.

[41] Pierre Colas. “Cyclin-dependent kinases and rare developmental disorders”. Orphanet Journal of Rare Diseases 15.1 (2020), p. 203. ISSN: 1750-1172.

[42] Charles R Scriver. “The PAH gene, phenylketonuria, and a paradigm shift”. Human Mutation 28.9 (2007), pp. 831–845.

[43] Henry G Zot et al. “Enhanced troponin I binding explains the functional changes produced by the hypertrophic cardiomyopathy mutation A8V of cardiac troponin C”. Arch. Biochem. Biophys. 601 (2016), pp. 97–104.

[44] Pierre Paoletti, Camilla Bellone, and Qiang Zhou. “NMDA receptor subunit diversity: impact on receptor properties, synaptic plasticity and disease”. Nature Reviews Neuroscience 14.6 (2013), pp. 383–400. ISSN: 1471-0048.

[45] Robert A. Copeland. “The drug–target residence time model: a 10-year retrospective”. Nature Reviews Drug Discovery 15.2 (2016), pp. 87–95. ISSN: 1474-1784.

[46] Hyung-June Woo and Benoît Roux. “Calculation of absolute protein-ligand binding free energy from computer simulations”. Proceedings of the National Academy of Sciences 102.19 (2005), pp. 6825–6830.

[47] Hernan G. Garcia et al. “A First Exposure to Statistical Mechanics for Life Scientists”. arXiv: Quantitative Methods (2007).

[48] Zheng Li and Themis Lazaridis. “The Effect of Water Displacement on Binding Thermodynamics: Concanavalin A”. The Journal of Physical Chemistry B 109.1 (2005). PMID: 16851059, pp. 662–670.

[49] Attila Szabo et al. “Reversible stochastically gated diffusion-influenced reactions”. The Journal of Chemical Physics 77.9 (1982), pp. 4484–4493. ISSN: 0021-9606.

[50] Huan-Xiang Zhou, Stanislaw T. Wlodek, and J. Andrew McCammon. “Conformation gating as a mechanism for enzyme specificity”. Proceedings of the National Academy of Sciences 95 (16 1998), pp. 9280–9283.

[51] G. Schreiber, G. Haran, and H.-X. Zhou. “Fundamental Aspects of Protein-Protein Association Kinetics”. Chemical Reviews 109.3 (2009). PMID: 19196002, pp. 839–860.

[52] Stephen J. Benkovic and Sharon Hammes-Schiffer. “A Perspective on Enzyme Catalysis”. Science 301.5637 (2003), p. 11961202.

[53] Tamara Frembgen-Kesner and Adrian H. Elcock. “Absolute Protein-Protein Association Rate Constants from Flexible, Coarse-Grained Brownian Dynamics Simulations: The Role of Intermolecular Hydrodynamic Interactions in Barnase-Barstar Association”. Biophysical Journal 99.9 (2010), pp. L75–L77. ISSN: 0006-3495.

[54] Michael Slutsky and Leonid A. Mirny. “Kinetics of Protein-DNA Interaction: Facilitated Target Location in Sequence-Dependent Potential”. Biophysical Journal 87.6 (2004), p. 40214035.

[55] R. R. Gabdoulline and R. C. Wade. “Simulation of the diffusional association of barnase and barstar”. Biophysical Journal 72.5 (1997), pp. 1917–1929. ISSN: 0006-3495.

[56] Bin Sun et al. “Electrostatic control of calcineurin’s intrinsically-disordered regulatory domain binding to calmodulin”. Biochimica et Biophysica Acta - General Subjects 1862.12 (2018), pp. 2651–2659. ISSN: 18728006.

[57] Chao Xu et al. “Accurate Characterization of Binding Kinetics and Allosteric Mechanisms for the HSP90 Chaperone Inhibitors Using AI-Augmented Integrative Biophysical Studies”. JACS Au 4.4 (2024), pp. 1632–1645.

[58] Te Liu et al. “Reconciling ASPP-p53 binding mode discrepancies through an ensemble binding framework that bridges crystallography and NMR data”. PLOS Computational Biology 20.2 (2024), pp. 1–20.

[59] Paul Robustelli, Stefano Piana, and David E Shaw. “Mechanism of coupled folding-upon-binding of an intrinsically disordered protein”. en. J. Am. Chem. Soc. 142.25 (2020), pp. 11092–11101.

[60] Milan Kumar Hazra and Yaakov Levy. “Affinity of disordered protein complexes is modulated by entropy–energy reinforcement”. Proceedings of the National Academy of Sciences 119.26 (2022), e2120456119.

[61] Josh Abramson et al. “Accurate structure prediction of biomolecular interactions with AlphaFold 3”. en. Nature 630.8016 (2024), pp. 493–500.

[62] Huan-Xiang Zhou and Paul A Bates. “Modeling protein association mechanisms and kinetics”. Current Opinion in Structural Biology 23.6 (2013), p. 887893.

[63] Riccardo Baron and J. Andrew McCammon. “Molecular Recognition and Ligand Association”. Annual Review of Physical Chemistry 64.1 (2013), p. 151175.

[64] Bruce J Berne, John D Weeks, and Ruhong Zhou. “Dewetting and hydrophobic interaction in physical and biological systems”. en. Annu. Rev. Phys. Chem. 60.1 (2009), pp. 85–103.

[65] P. N. Perera et al. “Observation of water dangling OH bonds around dissolved nonpolar groups”. Proceedings of the National Academy of Sciences 106.30 (2009), p. 1223012234.

[66] Farzin Sohraby and Ariane Nunes-Alves. “Advances in computational methods for ligand binding kinetics”. Trends in Biochemical Sciences 48.5 (2023), pp. 437–449. ISSN: 0968-0004.

[67] Inga Jarmoskaite et al. “How to measure and evaluate binding affinities”. eLife 9 (2020).

[68] Peter J. Tonge. “Quantifying the Interactions between Biomolecules: Guidelines for Assay Design and Data Analysis”. ACS Infectious Diseases 5.6 (2019), p. 796808.

[69] Margarida Bastos et al. “Isothermal titration calorimetry”. Nature Reviews Methods Primers 3.1 (2023).

[70] Lee A Freiburger, Karine Auclair, and Anthony K Mittermaier. “Elucidating protein binding mechanisms by variable-c ITC”. en. Chembiochem 10.18 (2009), pp. 2871–2873.

[71] Jeffrey P. Bonin et al. “Positive Cooperativity in Substrate Binding by Human Thymidylate Synthase”. Biophysical Journal 117.6 (2019), pp. 1074–1084. ISSN: 0006-3495.

[72] Hisham Mazal and Gilad Haran. “Single-molecule FRET methods to study the dynamics of proteins at work”. Current Opinion in Biomedical Engineering 12 (2019), pp. 8–17. ISSN: 2468-4511.

[73] Antoine M van Oijen. “Single-molecule approaches to characterizing kinetics of biomolecular interactions”. Current Opinion in Biotechnology 22.1 (2011). Analytical biotechnology, pp. 75–80. ISSN: 0958-1669.

[74] Bogumil Zelent et al. “Analysis of the co-operative interaction between the allosterically regulated proteins GK and GKRP using tryptophan fluorescence”. Biochemical Journal 459.3 (2014), pp. 551–564. ISSN: 0264-6021.

[75] Franz-Josef Meyer-Almes. “Kinetic binding assays for the analysis of protein–ligand interactions”. Drug Discovery Today: Technologies 17 (2015), pp. 1–8. ISSN: 1740-6749.

[76] Christian M. Heckmann and Francesca Paradisi. “Looking Back: A Short History of the Discovery of Enzymes and How They Became Powerful Chemical Tools”. ChemCatChem 12.24 (2020), pp. 6082–6102.

[77] Aki Fujioka et al. “Dynamics of the Ras/ERK MAPK cascade as monitored by fluorescent probes”. en. J. Biol. Chem. 281.13 (2006), pp. 8917–8926.

[78] Matthew W. Freyer and Edwin A. Lewis. Isothermal Titration Calorimetry: Experimental Design, Data Analysis, and Probing Macromolecule/Ligand Binding and Kinetic Interactions. Elsevier, 2008, pp. 79–113. ISBN: 9780123725202.

[79] Stephen R. Martin and Maria J. Schilstra. Interactions of a Signal Transduction Protein Investigated by Fluorescence Stopped-Flow Kinetics. Springer US, 2021, pp. 83–104. ISBN: 9781071611975.

[80] John F Eccleston and Jon P Hutchinson. Stopped-flow techniques. 2001, pp. 201–238. ISBN: 9781383049671.

[81] Quentin H. Gibson. Rapid mixing: Stopped flow. Elsevier, 1969, pp. 187–228. ISBN: 9780121818739.

[82] Tamanna Rob and Derek J. Wilson. “Time-Resolved Mass Spectrometry for Monitoring Millisecond Time-Scale Solution-Phase Processes”. European Journal of Mass Spectrometry 18.2 (2012), pp. 205–214.

[83] Claudio Poiesi et al. “Kinetic analysis of TNF-oligomer-monomer transition by surface plasmon resonance and immunochemical methods”. Cytokine 5.6 (1993), p. 539545.

[84] Robert V. Stahelin. “Surface plasmon resonance: a useful technique for cell biologists to characterize biomolecular interactions”. Molecular Biology of the Cell 24.7 (2013). Ed. by William Bement, pp. 883–886.

[85] Michael J. Roy et al. “SPR-Measured Dissociation Kinetics of PROTAC Ternary Complexes Influence Target Degradation Rate”. ACS Chemical Biology 14.3 (2019), p. 361368.

[86] Naman B. Shah and Thomas M. Duncan. “Bio-layer Interferometry for Measuring Kinetics of Protein-protein Interactions and Allosteric Ligand Effects”. Journal of Visualized Experiments 84 (2014).

[87] Myong-Hee Sung and James G. McNally. “Live cell imaging and systems biology”. WIREs Systems Biology and Medicine 3.2 (2010), pp. 167–182.

[88] Rajesh Babu Sekar and Ammasi Periasamy. “Fluorescence resonance energy transfer (FRET) microscopy imaging of live cell protein localizations”. The Journal of Cell Biology 160.5 (2003), pp. 629–633.

[89] Awadhesh Kumar Verma et al. “FRET Based Biosensor: Principle Applications Recent Advances and Challenges”. Diagnostics 13.8 (2023), p. 1375.

[90] Felix Schiele, Pelin Ayaz, and Amaury Fernández-Montalván. “A universal homogeneous assay for high-throughput determination of binding kinetics”. Analytical Biochemistry 468 (2015), pp. 42–49. ISSN: 0003-2697.

[91] Victoria Georgi et al. “Binding Kinetics Survey of the Drugged Kinome”. Journal of the American Chemical Society 140.46 (2018). PMID: 30362749, pp. 15774–15782.

[92] Mark E Bowen et al. “Single molecule observation of liposome-bilayer fusion thermally induced by soluble N-ethyl maleimide sensitive-factor attachment protein receptors (SNAREs)”. Biophysical journal 87.5 (2004), pp. 3569–3584.

[93] Yulong Li, George J. Augustine, and Keith Weninger. “Kinetics of Complexin Binding to the SNARE Complex: Correcting Single Molecule FRET Measurements for Hidden Events”. Biophysical Journal 93.6 (2007), pp. 2178–2187. ISSN: 0006-3495.

[94] Ming-Li Zhang et al. “Interplay Between Intracellular Transport Dynamics and Liquid–Liquid Phase Separation”. Advanced Science 11.19 (2024), p. 2308338.

[95] Xin Jin et al. “Membraneless organelles formed by liquid-liquid phase separation increase bacterial fitness”. Science Advances 7.43 (2021), eabh2929.

[96] Gorka Muñoz-Gil et al. “Stochastic particle unbinding modulates growth dynamics and size of transcription factor condensates in living cells”. Proceedings of the National Academy of Sciences 119.31 (2022), e2200667119.

[97] Marko Vendelin and Rikke Birkedal. “Anisotropic diffusion of fluorescently labeled ATP in rat cardiomyocytes determined by raster image correlation spectroscopy”. American Journal of Physiology-Cell Physiology 295.5 (2008), pp. C1302–C1315.

[98] Ian R. Kleckner and Mark P. Foster. “An introduction to NMR-based approaches for measuring protein dynamics”. Biochimica et Biophysica Acta (BBA) - Proteins and Proteomics 1814.8 (2011), pp. 942–968.

[99] Jeffrey A. Purslow et al. “NMR Methods for Structural Characterization of Protein-Protein Complexes”. Frontiers in Molecular Biosciences 7 (2020).

[100] Roberto N De Guzman et al. “Structural Basis for Cooperative Transcription Factor Binding to the CBP Coactivator”. Journal of Molecular Biology 355.5 (2006), pp. 1005–1013. ISSN: 0022-2836.

[101] Riley B. Peacock and Elizabeth A. Komives. “Hydrogen/Deuterium Exchange and Nuclear Magnetic Resonance Spectroscopy Reveal Dynamic Allostery on Multiple Time Scales in the Serine Protease Thrombin”. Biochemistry 60.46 (2021), p. 34413448.

[102] Jean-Philippe Demers and Anthony Mittermaier. “Binding Mechanism of an SH3 Domain Studied by NMR and ITC”. Journal of the American Chemical Society 131.12 (2009). PMID: 19267471, pp. 4355–4367.

[103] Nikos S. Hatzakis. “Single molecule insights on conformational selection and induced fit mechanism”. Biophysical Chemistry 186 (2014). Special issue : conformational selection, pp. 46–54. ISSN: 0301-4622.

[104] Fabian Paul and Thomas R. Weikl. “How to Distinguish Conformational Selection and Induced Fit Based on Chemical Relaxation Rates”. PLOS Computational Biology 12.9 (2016), pp. 1–17.

[105] Ekaterina Morgunova and Jussi Taipale. “Structural perspective of cooperative transcription factor binding”. Current Opinion in Structural Biology 47 (2017), pp. 1–8. ISSN: 0959-440X.

[106] Minkyung Baek and David Baker. “Accurate prediction of protein structures and interactions using a three-track neural network”. Science 373.6557 (2021), pp. 871–876.

[107] V A Jisna and P B Jayaraj. “Protein structure prediction: Conventional and deep learning perspectives”. en. Protein J. 40.4 (2021), pp. 522–544.

[108] Justin T Seffernick and Steffen Lindert. “Hybrid methods for combined experimental and computational determination of protein structure”. en. J. Chem. Phys. 153.24 (2020), p. 240901.

[109] M. P. Allen and D. J. Tildesley. Computer Simulation of Liquids. Oxford: Oxford University Press, 1989.

[110] Berend Smit and Theo L M Maesen. “Molecular Simulations of Zeolites: Adsorption, Diffusion, and Shape Selectivity”. Chemical Reviews 108 (10 2008), pp. 4125–4184.

[111] Simon Berneche and Benoit Roux. “A microscopic view of ion conduction through the K+ channel.” Proceedings of the National Academy of Sciences 100 (15 2003), pp. 8644–8648.

[112] Donald L. Ermak and J. A. McCammon. “Brownian dynamics with hydrodynamic interactions”. The Journal of Chemical Physics 69.4 (1978), pp. 1352–1360. ISSN: 0021-9606.

[113] Jens Rotne and Stephen Prager. “Variational Treatment of Hydrodynamic Interaction in Polymers”. The Journal of Chemical Physics 50.11 (1969), pp. 4831–4837. ISSN: 0021-9606.

[114] P. Debye. “Reaction Rates in Ionic Solutions”. Transactions of The Electrochemical Society 82.1 (1942), p. 265.

[115] CH Bamford, CFH Tipper, and RG Compton. Diffusion-Limited Reactions. Vol. 25. Comprehensive Chemical Kinetics. Elsevier.

[116] Hannes Risken. “The Fokker-Planck Equation: Methods of Solution and Applications”. Springer Series in Synergetics (1996).

[117] Jacob N Israelachvili. Intermolecular and surface forces. Academic press, 2011.

[118] Lin Li et al. “DelPhi: a comprehensive suite for DelPhi software and associated resources”. BMC Biophysics 5.1 (2012), p. 9. ISSN: 2046-1682.

[119] Elizabeth Jurrus et al. “Improvements to the APBS biomolecular solvation software suite”. Protein science 27.1 (2018), pp. 112–128.

[120] Yuhua Song et al. “Finite Element Solution of the Steady-State Smoluchowski Equation for Rate Constant Calculations”. Biophysical Journal 86.4 (2004), pp. 2017–2029.

[121] Ramzi Alsallaq and Huan-Xiang Zhou. “Electrostatic rate enhancement and transient complex of protein-protein association”. Proteins-Structure Function And Bioinformatics 71.1 (2008), pp. 320–335.

[122] Gary A. Huber and J. Andrew McCammon. “Browndye: A software package for Brownian dynamics”. Computer Physics Communications 181.11 (2010), pp. 1896–1905. ISSN: 0010-4655.

[123] Gary A Huber and J Andrew McCammon. “Brownian dynamics simulations of biological molecules”. en. Trends Chem. 1.8 (2019), pp. 727–738.

[124] S H Northrup and H P Erickson. “Kinetics of protein-protein association explained by Brownian dynamics computer simulation.” Proceedings of the National Academy of Sciences 89.8 (1992), pp. 3338–3342.

[125] Razif R. Gabdoulline and Rebecca C. Wade. “Protein-protein association: investigation of factors influencing association rates by Brownian dynamics simulations1 1Edited by B. Honig”. Journal of Molecular Biology 306.5 (2001), pp. 1139–1155. ISSN: 0022-2836.

[126] Jeffry D Madura et al. “Electrostatics and diffusion of molecules in solution: simulations with the University of Houston Brownian Dynamics program”. Computer Physics Communications 91.1 (1995), pp. 57–95. ISSN: 0010-4655.

[127] Michael Martinez et al. “SDA 7: A modular and parallel implementation of the simulation of diffusional association software”. Journal of Computational Chemistry 36.21 (2015), pp. 1631–1645.

[128] Maciej Dlugosz, Pawel Zieliński, and Joanna Trylska. “Brownian dynamics simulations on CPU and GPU with BD BOX”. Journal of Computational Chemistry 32.12 (2011), pp. 2734–2744.

[129] Moritz Hoffmann, Christoph Fröhner, and Frank Noé. “ReaDDy 2: Fast and flexible software framework for interacting-particle reaction dynamics”. PLOS Computational Biology 15.2 (2019), pp. 1–26.

[130] Timothy Cholko et al. “GeomBD3: Brownian Dynamics Simulation Software for Biological and Engineered Systems”. Journal of Chemical Information and Modeling 62.10 (2022), pp. 2257–2263.

[131] Lane W Votapka et al. “SEEKR: Simulation Enabled Estimation of Kinetic Rates, A computational tool to estimate molecular kinetics and its application to trypsin-benzamidine binding”. en. J. Phys. Chem. B 121.15 (2017), pp. 3597–3606.

[132] G. A. Huber and S. Kim. “Weighted-ensemble Brownian dynamics simulations for protein association reactions”. Biophysical Journal 70.1 (1996), pp. 97–110. ISSN: 0006-3495.

[133] Gang Zou, Robert D. Skeel, and Shankar Subramaniam. “Biased Brownian Dynamics for Rate Constant Calculation”. Biophysical Journal 79.2 (2000), pp. 638–645. ISSN: 0006-3495.

[134] BrianT. Castle and DavidJ. Odde. “Brownian Dynamics of Subunit Addition-Loss Kinetics and Thermodynamics in Linear Polymer Self-Assembly”. Biophysical Journal 105.11 (2013), p. 25282540.

[135] Henry Eyring. “The Theory of Absolute Reaction Rates”. The Journal of Chemical Physics 3.2 (1935), pp. 107–115.

[136] V.I. Melnikov. “The Kramers problem: Fifty years of development”. Physics Reports 209.1 (1991), pp. 1–71. ISSN: 0370-1573.

[137] Huan-Xiang Zhou. “Rate theories for biologists”. Quarterly Reviews of Biophysics 43.2 (2010), p. 219293.

[138] Alexander M Berezhkovskii, Attila Szabo, and Huan-Xiang Zhou. “Diffusion-influenced ligand binding to buried sites in macromolecules and transmembrane channels”. en. J. Chem. Phys. 135.7 (2011), p. 075103.

[139] M Hinczewski et al. “How the diffusivity profile reduces the arbitrariness of protein folding free energies”. The Journal of chemical physics 132 (24 2010), p. 245103.

[140] Philipp Metzner, Christof Schtte, and Eric Vanden-Eijnden. “Transition Path Theory for Markov Jump Processes”. Multiscale Modeling & Simulation 7.3 (2009), p. 11921219.

[141] Ron Elber. “Milestoning: An Efficient Approach for Atomically Detailed Simulations of Kinetics in Biophysics”. Annual Review of Biophysics 49.1 (2020), p. 6985.

[142] Ernesto Suárez, Joshua L Adelman, and Daniel M Zuckerman. “Accurate estimation of protein folding and unfolding times: Beyond Markov state models”. en. J. Chem. Theory Comput. 12.8 (2016), pp. 3473–3481.

[143] Brooke E Husic and Vijay S Pande. “Markov State Models: From an Art to a Science”. Journal of the American Chemical Society 140.7 (2018), pp. 2386–2396.

[144] Hao Wu and Frank Noé. “Variational Approach for Learning Markov Processes from Time Series Data”. Journal of Nonlinear Science 30.1 (2020), pp. 23–66. ISSN: 1432-1467.

[145] Andreas Mardt et al. “VAMPnets for deep learning of molecular kinetics”. Nature Communications 9.1 (2018), p. 5. ISSN: 2041-1723.

[146] Guillermo Pérez-Hernández et al. “Identification of slow molecular order parameters for Markov model construction”. The Journal of Chemical Physics 139.1 (2013), p. 015102. ISSN: 0021-9606.

[147] Christian R Schwantes and Vijay S Pande. “Improvements in Markov State Model Construction Reveal Many Non-Native Interactions in the Folding of NTL9”. Journal of Chemical Theory and Computation 9.4 (2013), pp. 2000–2009.

[148] Feng Qin, Anthony Auerbach, and Frederick Sachs. “A Direct Optimization Approach to Hidden Markov Modeling for Single Channel Kinetics”. Biophysical Journal 79.4 (2000), p. 19151927.

[149] Gerhard Hummer and Attila Szabo. “Kinetics from Nonequilibrium Single-Molecule Pulling Experiments”. Biophysical Journal 85.1 (2003), pp. 5–15.

[150] Doris A. Schuetz et al. “Kinetics for Drug Discovery: an industry-driven effort to target drug residence time”. Drug Discovery Today 22.6 (2017), pp. 896–911. ISSN: 1359-6446.

[151] G. Vauquelin and A. Packeu. “Ligands, their receptors and plasma membranes”. Molecular and Cellular Endocrinology 311.12 (2009), p. 110.

[152] Gen Honda et al. “Slow diffusion and signal amplification on membranes regulated by phospholipase D” (2024).

[153] William Y. C. Huang, Steven G. Boxer, and James E. Ferrell. “Membrane localization accelerates association under conditions relevant to cellular signaling”. Proceedings of the National Academy of Sciences 121.10 (2024).

[154] Yu-Jo Chai et al. “Heterogeneous nanoscopic lipid diffusion in the live cell membrane and its dependency on cholesterol”. Biophysical Journal 121.16 (2022), pp. 3146–3161.

[155] Zhaoqian Su, Kalyani Dhusia, and Yinghao Wu. “Understanding the functional role of membrane confinements in TNF-mediated signaling by multiscale simulations”. Communications Biology 5.1 (2022), p. 228. ISSN: 2399-3642.

[156] Timothy Cholko and Chia-en A. Chang. “Modeling Effects of Surface Properties and Probe Density for Nanoscale Biosensor Design: A Case Study of DNA Hybridization near Surfaces”. The Journal of Physical Chemistry B 125.7 (2021), pp. 1746–1754.

[157] Johan Hake, Peter M Kekenes-Huskey, and Andrew D McCulloch. “Computational modeling of subcellular transport and signaling”. Current Opinion in Structural Biology 25 (2014), pp. 92–97.

[158] S Kashif Sadiq et al. “Multiscale Approach for Computing Gated Ligand Binding from Molecular Dynamics and Brownian Dynamics Simulations”. Journal of Chemical Theory and Computation 17.12 (2021), pp. 7912–7929.

[159] Tom J Petty et al. “An induced fit mechanism regulates p53 DNA binding kinetics to confer sequence specificity”. The EMBO Journal 30.11 (2011), pp. 2167–2176.

[160] Xiaorong Liu, Jianlin Chen, and Jianhan Chen. “Residual Structure Accelerates Binding of Intrinsically Disordered ACTR by Promoting Efficient Folding upon Encounter”. Journal of Molecular Biology 431.2 (2019), pp. 422–432. ISSN: 0022-2836.

[161] Joseph M. Rogers, Chi T. Wong, and Jane Clarke. “Coupled Folding and Binding of the Disordered Protein PUMA Does Not Require Particular Residual Structure”. Journal of the American Chemical Society 136.14 (2014). PMID: 24654952, pp. 5197–5200.

[162] R.C. Wade et al. “Gating of the active site of triose phosphate isomerase: Brownian dynamics simulations of flexible peptide loops in the enzyme”. Biophysical Journal 64.1 (1993), pp. 9–15. ISSN: 0006-3495.

[163] Rebecca C. Wade et al. “Simulation of enzyme–substrate encounter with gated active sites”. Nature Structural Biology 1.1 (1994), pp. 65–69. ISSN: 1545-9985.

[164] Nicholas Greives and Huan-Xiang Zhou. “BDflex: A method for efficient treatment of molecular flexibility in calculating protein-ligand binding rate constants from Brownian dynamics simulations”. The Journal of chemical physics 137.13 (2012), p. 135105.

[165] Robert B Best. “Computational and theoretical advances in studies of intrinsically disordered proteins”. Current Opinion in Structural Biology 42 (2017). Folding and binding • Proteins: Bridging theory and experiment, pp. 147–154. ISSN: 0959-440X.

[166] W. Wendell Smith, Po-Yi Ho, and Corey S. O’Hern. “Calibrated Langevin-dynamics simulations of intrinsically disordered proteins”. Phys. Rev. E 90 (4 2014), p. 042709.

[167] Tamara Frembgen-Kesner et al. “Parametrization of backbone flexibility in a coarse-grained Force Field for proteins (COFFDROP) derived from all-atom explicit-solvent molecular dynamics simulations of all possible two-residue peptides”. en. J. Chem. Theory Comput. 11.5 (2015), pp. 2341–2354.

[168] Ioana M Ilie, Wouter K den Otter, and Wim J Briels. “A coarse grained protein model with internal degrees of freedom. Application to *α*-synuclein aggregation”. en. J. Chem. Phys. 144.8 (2016), p. 085103.

[169] Steffen Mühle et al. “Loop formation and translational diffusion of intrinsically disordered proteins”. en. Phys. Rev. E. 100. 5-1 (2019), p. 052405.

[170] Jinan Wang et al. “Predicting Biomolecular Binding Kinetics: A Review”. Journal of Chemical Theory and Computation 19.8 (2023), pp. 2135–2148.

[171] Yinglong Miao, Apurba Bhattarai, and Jinan Wang. “Ligand Gaussian Accelerated Molecular Dynamics (LiGaMD): Characterization of Ligand Binding Thermodynamics and Kinetics”. Journal of Chemical Theory and Computation 16.9 (2020). PMID: 32692556, pp. 5526–5547.

[172] Ron O. Dror et al. “Pathway and mechanism of drug binding to G-protein-coupled receptors”. Proceedings of the National Academy of Sciences 108.32 (2011), pp. 13118–13123.

[173] Yibing Shan et al. “How Does a Drug Molecule Find Its Target Binding Site?” Journal of the American Chemical Society 133.24 (2011). PMID: 21545110, pp. 9181–9183.

[174] Daria B. Kokh et al. “Estimation of Drug-Target Residence Times by *τ*-Random Acceleration Molecular Dynamics Simulations”. Journal of Chemical Theory and Computation 14.7 (2018). PMID: 29768913, pp. 3859–3869.

[175] Dhiman Ray and Michele Parrinello. “Kinetics from Metadynamics: Principles, Applications, and Outlook”. Journal of Chemical Theory and Computation 19.17 (2023). PMID: 37585703, pp. 5649–5670.

[176] Samuel D Lotz and Alex Dickson. “Unbiased Molecular Dynamics of 11 min Timescale Drug Unbinding Reveals Transition State Stabilizing Interactions”. Journal of the American Chemical Society 140.2 (2018). PMID: 29303257, pp. 618–628.

[177] Neil J Bruce et al. “New approaches for computing ligand-receptor binding kinetics”. Current Opinion in Structural Biology 49 (2018), pp. 1–10. ISSN: 0959-440X.

[178] William Sinko et al. “ModBind, a Rapid Simulation-Based Predictor of Ligand Binding and Off-Rates”. Journal of Chemical Information and Modeling 65.1 (2025). PMID: 39681514, pp. 265–274.

[179] Joel Berry et al. “RNA transcription modulates phase transition-driven nuclear body assembly”. Proceedings of the National Academy of Sciences 112.38 (2015), E5237–E5245.

[180] Eric D. Siggia. “Late stages of spinodal decomposition in binary mixtures”. Phys. Rev. A 20 (2 1979), pp. 595–605.

[181] Yifeng Qi and Bin Zhang. “Chromatin network retards nucleoli coalescence”. Nature Communications 12.1 (2021), p. 6824. ISSN: 2041-1723.

[182] Jingpeng Zhang, Yanyi Huang, and Fan Bai. “Stochastic Monte Carlo Model for Simulating the Dynamic Liquid–Liquid Phase Separation in Bacterial Cells”. The Journal of Physical Chemistry B 127.18 (2023). PMID: 37130439, pp. 4145–4153.

[183] Paul E. Schavemaker, Arnold J. Boersma, and Bert Poolman. “How Important Is Protein Diffusion in Prokaryotes?” Frontiers in Molecular Biosciences 5 (2018).

[184] Huan-Xiang Zhou, Germán Rivas, and Allen P Minton. “Macromolecular crowding and confinement: biochemical, biophysical, and potential physiological consequences”. en. Annu. Rev. Biophys. 37.1 (2008), pp. 375–397.

[185] Germán Rivas and Allen P. Minton. “Macromolecular Crowding In Vitro, In Vivo, and In Between”. Trends in Biochemical Sciences 41.11 (2016), pp. 970–981.

[186] Begoña Monterroso et al. “Macromolecular Crowding, Phase Separation, and Homeostasis in the Orchestration of Bacterial Cellular Functions”. Chemical Reviews 124.4 (2024), pp. 1899–1949.

[187] Anthony A. Hyman, Christoph A. Weber, and Frank Jülicher. “Liquid-Liquid Phase Separation in Biology”. Annual Review of Cell and Developmental Biology 30.1 (2014), pp. 39–58.

[188] Bhawna Saini and Tushar Kanti Mukherjee. “Biomolecular Condensates Regulate Enzymatic Activity under a Crowded Milieu: Synchronization of Liquid–Liquid Phase Separation and Enzymatic Transformation”. The Journal of Physical Chemistry B 127.1 (2023). PMID: 36594499, pp. 180–193.

[189] Andrea Papale and David Holcman. “Chromatin phase separated nanoregions explored by polymer cross-linker models and reconstructed from single particle trajectories”. PLOS Computational Biology 20.1 (2024), pp. 1–21.

[190] Marcos Gil-Garcia et al. “Local environment in biomolecular condensates modulates enzymatic activity across length scales”. Nature Communications 15.1 (2024), p. 3322. ISSN: 2041-1723.

[191] Yi Zhang et al. “MORC3 forms nuclear condensates through phase separation”. IScience 17 (2019), pp. 182–189.

[192] Maria Hondele et al. “DEAD-box ATPases are global regulators of phase-separated organelles”. Nature 573.7772 (2019), pp. 144–148.

[193] J C Maxwell Garnett. “XII. Colours in metal glasses and in metallic films”. Philosophical Transactions of the Royal Society of London. Series A, Containing Papers of a Mathematical or Physical Character 203. 359-371 (1904), pp. 385–420.

[194] Grzegorz Wieczorek and Piotr Zielenkiewicz. “Influence of macromolecular crowding on protein-protein association rates–a Brownian dynamics study”. en. Biophys. J. 95.11 (2008), pp. 5030–5036.

[195] Igor M. Sokolov. “Models of anomalous diffusion in crowded environments”. Soft Matter 8 (35 2012), pp. 9043–9052.

[196] Mikita M. Misiura and Anatoly B. Kolomeisky. “Role of Intrinsically Disordered Regions in Acceleration of Protein–Protein Association”. The Journal of Physical Chemistry B 124.1 (2020). PMID: 31804089, pp. 20–27.

[197] Christopher C. Roberts and Chia-en A. Chang. “Analysis of Ligand–Receptor Association and Intermediate Transfer Rates in Multienzyme Nanostructures with All-Atom Brownian Dynamics Simulations”. The Journal of Physical Chemistry B 120.33 (2016), pp. 8518–8531.

[198] Isseki Yu et al. “Biomolecular interactions modulate macromolecular structure and dynamics in atomistic model of a bacterial cytoplasm”. eLife 5 (2016). Ed. by Yibing Shan, e19274. ISSN: 2050-084X.

[199] Sen Yang et al. “Self-construction of actin networks through phase separation–induced abLIM1 condensates”. Proceedings of the National Academy of Sciences 119.29 (2022), e2122420119.

[200] Utkarsh Kapoor, Young C. Kim, and Jeetain Mittal. “Coarse-Grained Models to Study Protein–DNA Interactions and Liquid–Liquid Phase Separation”. Journal of Chemical Theory and Computation 20.4 (2024), pp. 1717–1731.

[201] Lin-ge Li and Zhonghuai Hou. “Theoretical modelling of liquid–liquid phase separation: from particle-based to field-based simulation”. Biophysics Reports 8.2 (2022), pp. 55–67. ISSN: 2364-3439.

[202] J L Auriault and J Lewandowska. “Effective diffusion coefficient: from homogenization to experiment”. Transport in porous media 27 (2 1997), pp. 205–223.

[203] Jean Louis Auriault, Claude Boutin, and Christian Geindreau. “Homogenization of Coupled Phenomena in Heterogenous Media”. Homogenization of Coupled Phenomena in Heterogenous Media. Wiley-ISTE, 2010. ISBN: 9781848211612.

[204] Tom Pace et al. “Homogenization of Continuum-Scale Transport Properties from Molecular Dynamics Simulations: An Application to Aqueous-Phase Methane Diffusion in Silicate Channels”. The Journal of Physical Chemistry B 125.41 (2021), pp. 11520–11533.

[205] Peter M Kekenes-Huskey, Caitlin E Scott, and Selcuk Atalay. “Quantifying the Influence of the Crowded Cytoplasm on Small Molecule Diffusion”. The Journal of Physical Chemistry B (2016), acs.jpcb.6b03887.

[206] Peter M. Kekenes-Huskey et al. “Molecular and Subcellular-Scale Modeling of Nucleotide Diffusion in the Cardiac Myofilament Lattice”. Biophysical Journal 105.9 (2013), pp. 2130–2140. ISSN: 0006-3495.

[207] P R Shorten and J sneyd. “A Mathematical Analysis of Obstructed Diffusion within Skeletal Muscle”. Biophysical Journal 96 (12 2009), pp. 4764–4778.

[208] Christine Deisl, Jay H. Chung, and Donald W. Hilgemann. “Longitudinal diffusion barriers imposed by myofilaments and mitochondria in murine cardiac myocytes”. Journal of General Physiology 155.10 (2023), e202213329. ISSN: 0022-1295.

[209] B B Roux. “Statistical mechanical equilibrium theory of selective ion channels”. Biophysical Journal 77 (1 1999), pp. 139–153.

[210] P. M. Kekenes-Huskey, A. K. Gillette, and J. A. McCammon. “Predicting the influence of long-range molecular interactions on macroscopic-scale diffusion by homogenization of the Smoluchowski equation”. The Journal of Chemical Physics 140.17 (2014), p. 174106. ISSN: 0021-9606.

[211] Ariane Nunes-Alves, Daria B Kokh, and Rebecca C Wade. “Recent progress in molecular simulation methods for drug binding kinetics”. Current Opinion in Structural Biology 64 (2020), pp. 126–133. ISSN: 0959-440X.

[212] Gregory R Bowman, Xuhui Huang, and Vijay S Pande. “Using generalized ensemble simulations and Markov state models to identify conformational states”. Methods (San Diego, Calif) 49 (2 2009), pp. 197–201.

[213] J Andrew McCammon. “Gated Diffusion-controlled Reactions”. BMC Biophysics 4.1 (2011).

[214] Chia-En Chang et al. “Gated Binding of Ligands to HIV-1 Protease: Brownian Dynamics Simulations in a Coarse-Grained Model”. Biophysical Journal 90.11 (2006), pp. 3880–3885. ISSN: 0006-3495.

[215] Yu-ming M. Huang, Myungshim Kang, and Chia-en A. Chang. “Mechanistic Insights into Phosphopeptide-BRCT Domain Association: Preorganization, Flexibility, and Phosphate Recognition”. The Journal of Physical Chemistry B 116.34 (2012), pp. 10247–10258.

[216] Lane W Votapka and Rommie E Amaro. “Multiscale Estimation of Binding Kinetics Using Brownian Dynamics, Molecular Dynamics and Milestoning”. PLOS Computational Biology 11.10 (2015), pp. 1–24.

[217] Lane W Votapka et al. “SEEKR2: Versatile multiscale milestoning utilizing the OpenMM molecular dynamics engine”. Journal of chemical information and modeling 62.13 (2022), pp. 3253–3262.

[218] Marcellus Ubbink. “The courtship of proteins: Understanding the encounter complex”. FEBS Letters 583.7 (2009), pp. 1060–1066. ISSN: 0014-5793.

[219] Ali S Saglam and Lillian T Chong. “Protein–protein binding pathways and calculations of rate constants using fully-continuous, explicit-solvent simulations”. Chem. Sci. 10.8 (2019), pp. 2360–2372.

[220] J. Robert Lane et al. “A kinetic view of GPCR allostery and biased agonism”. Nature Chemical Biology 13.9 (2017), pp. 929–937. ISSN: 1552-4469.

[221] Brian T. DeVree et al. “Allosteric coupling from G protein to the agonist-binding pocket in GPCRs”. Nature 535.7610 (2016), pp. 182–186. ISSN: 1476-4687.

[222] Jonathan P Davis et al. “Effects of thin and thick filament proteins on calcium binding and exchange with cardiac troponin C”. en. Biophys. J. 92.9 (2007), pp. 3195–3206.

[223] Gozde Kar et al. “Allostery and population shift in drug discovery”. Current Opinion in Pharmacology 10.6 (2010), p. 715722.

[224] Jacque Monod, Jeffries Wyman, and Jean-Pierre Changeux. “On the nature of allosteric transitions: a plausible model”. Journal of molecular biology 12.1 (1965), pp. 88–118.

[225] Meritxell Canals et al. “A Monod-Wyman-Changeux Mechanism Can Explain G Proteincoupled Receptor (GPCR) Allosteric Modulation *¡sup¿ ¡/sup¿”. Journal of Biological Chemistry 287.1 (2012), pp. 650–659. ISSN: 0021-9258.

[226] S Bowerman and J Wereszczynski. “Detecting Allosteric Networks Using Molecular Dynamics Simulation”. Computational Approaches for Studying Enzyme Mechanism Part B. Ed. by Gregory A Voth. Vol. 578. Methods in Enzymology. Academic Press, 2016. Chap. 17, pp. 429–447.

[227] Brian C Goodwin. “Temporal organization in cells. A dynamic theory of cellular control processes.” (1963).

[228] Didier Gonze and Peter Ruoff. “The Goodwin Oscillator and its Legacy”. Acta Biotheoretica 69.4 (2021), pp. 857–874. ISSN: 1572-8358.

[229] Didier Gonze and Wassim Abou-Jaoudé. “The Goodwin Model: Behind the Hill Function”. PLoS ONE 8.8 (2013). Ed. by Nick Monk, e69573.

[230] Paul A. Srere. “COMPLEXES OF SEQUENTIAL METABOLIC ENZYMES”. Annual Review of Biochemistry 56.1 (1987), p. 89124.

[231] Yan Jiang et al. “Electrostatic Steering Enables Flow-Activated Von Willebrand Factor to Bind Platelet Glycoprotein, Revealed by Single-Molecule Stretching and Imaging”. Journal of Molecular Biology 431.7 (2019), pp. 1380–1396. ISSN: 0022-2836.

[232] Harkewal Singh et al. “Structures of the PutA peripheral membrane flavoenzyme reveal a dynamic substrate-channeling tunnel and the quinone-binding site”. Proceedings of the National Academy of Sciences 111.9 (2014), pp. 3389–3394.

[233] Vincent T Metzger et al. “Electrostatic channeling in P. falciparum DHFR-TS: Brownian dynamics and Smoluchowski modeling”. en. Biophys. J. 107.10 (2014), pp. 2394–2402.

[234] Marc Leibundgut et al. “Structural Basis for Substrate Delivery by Acyl Carrier Protein in the Yeast Fatty Acid Synthase”. Science (New York, N.Y.) 316 (2007), pp. 288–90.

[235] Lee J. Sweetlove and Alisdair R. Fernie. “The role of dynamic enzyme assemblies and substrate channelling in metabolic regulation”. Nature Communications 9.1 (2018), p. 2136. ISSN: 041-1723.

[236] Changsun Eun et al. “A model study of sequential enzyme reactions and electrostatic channeling”. The Journal of Chemical Physics 140.10 (2014), p. 105101. ISSN: 0021-9606.

[237] Hadi Rahmaninejad et al. “Crowding within synaptic junctions influences the degradation of nucleotides by CD39 and CD73 ectonucleotidases”. Biophysical Journal 121.2 (2022), pp. 309–318. ISSN: 0006-3495.

[238] Yu-Ming M Huang et al. “Brownian dynamic study of an enzyme metabolon in the TCA cycle: Substrate kinetics and channeling”. en. Protein Sci. 27.2 (2018), pp. 463–471.

[239] Yuanchao Liu et al. “Cascade Kinetics of an Artificial Metabolon by Molecular Dynamics and Kinetic Monte Carlo”. ACS Catalysis 8.8 (2018), pp. 7719–7726.

[240] Arthur F. Voter. “INTRODUCTION TO THE KINETIC MONTE CARLO METHOD”. Radiation Effects in Solids. Ed. by Kurt E. Sickafus, Eugene A. Kotomin, and Blas P. Uberuaga. Dordrecht: Springer Netherlands, 2007, pp. 1–23. ISBN: 978-1-4020-5295-8.

[241] Michael T. Klann, Alexei Lapin, and Matthias Reuss. “Agent-based simulation of reactions in the crowded and structured intracellular environment: Influence of mobility and location of the reactants”. BMC Systems Biology 5.1 (2011), p. 71. ISSN: 1752-0509.

[242] Maciej Dobrzyński et al. “Computational methods for diffusion-influenced biochemical reactions”. Bioinformatics 23.15 (2007), pp. 1969–1997.

[243] Jinglin Fu et al. “Interenzyme Substrate Diffusion for an Enzyme Cascade Organized on Spatially Addressable DNA Nanostructures”. Journal of the American Chemical Society 134.12 (2012). PMID: 22414276, pp. 5516–5519.

[244] Nuo Wang and J. Andrew McCammon. “Substrate channeling between the human dihydrofolate reductase and thymidylate synthase”. Protein Science 25.1 (2016), pp. 79–86.

[245] Jason Yang, Francesca-Zhoufan Li, and Frances H. Arnold. “Opportunities and Challenges for Machine Learning-Assisted Enzyme Engineering”. ACS Central Science 10.2 (2024), pp. 226–241.

[246] Scott A. Hollingsworth and Ron O. Dror. “Molecular Dynamics Simulation for All”. Neuron 99.6 (2018), p. 11291143.

[247] Fu Lin and Renxiao Wang. “Systematic Derivation of AMBER Force Field Parameters Applicable to Zinc-Containing Systems”. Journal of Chemical Theory and Computation (2010).

[248] Julien Michel, Julian Tirado-Rives, and William L. Jorgensen. “Prediction of the Water Content in Protein Binding Sites”. The Journal of Physical Chemistry B 113.40 (2009), p. 1333713346.

[249] Paul Katzberger and Sereina Riniker. “A general graph neural network based implicit solvation model for organic molecules in water”. Chem. Sci. 15 (28 2024), pp. 10794–10802.

[250] Alexander N. Volkov. “Structure and Function of Transient Encounters of Redox Proteins”. Accounts of Chemical Research 48.12 (2015), pp. 3036–3043.

[251] Benjamin P. Tu and Steven L. McKnight. “Metabolic cycles as an underlying basis of biological oscillations”. Nature Reviews Molecular Cell Biology 7.9 (2006), p. 696701.

[252] Bin Sun et al. “Simulation-Based Characterization of Electrolytes and Small Molecule Diffusion in Oriented Mesoporous Silica Thin Films”. Computational Materials, Chemistry, and Biochemistry: From Bold Initiatives to the Last Mile: In Honor of William A. Goddard’s Contributions to Science and Engineering. Ed. by Sadasivan Shankar et al. Cham: Springer International Publishing, 2021, pp. 521–558. ISBN: 978-3-030-18778-1.

[253] Monica X Li, Leo Spyracopoulos, and Brian D Sykes. “Binding of cardiac troponin-I147-163 induces a structural opening in human cardiac troponin-C”. Biochemistry 38 (26 1999), pp. 8289–8298.

[254] Bin Sun and Peter M. Kekenes-Huskey. Myofilament-associated proteins with intrinsic disorder (MAPIDs) and their resolution by computational modeling. 2023.

[255] Charlotte S. Srensen and Magnus Kjaergaard. “Effective concentrations enforced by intrinsically disordered linkers are governed by polymer physics”. Proceedings of the National Academy of Sciences 116.46 (2019), p. 2312423131.

[256] EstellaA. Newcombe et al. “How phosphorylation impacts intrinsically disordered proteins and their function”. Essays in Biochemistry 66.7 (2022). Ed. by Samrat Mukhopadhyay, p. 901913.

[257] Munehito Arai et al. “Conformational propensities of intrinsically disordered proteins influence the mechanism of binding and folding”. Proceedings of the National Academy of Sciences 112.31 (2015), p. 96149619.

[258] Thomas A. Leonard, Martin Loose, and Sascha Martens. “The membrane surface as a platform that organizes cellular and biochemical processes”. Developmental Cell 58.15 (2023), p. 13151332.

[259] Bas Teusink et al. “Can yeast glycolysis be understood in terms of in vitro kinetics of the constituent enzymes? Testing biochemistry”. European Journal of Biochemistry 267.17 (2000), pp. 5313–5329.

[260] Xuan Fang et al. “A Bayesian framework for systems model refinement and selection of calcium signaling”. Biophysical Journal 124.14 (2025), p. 23472361.

[261] Jianyong Sun, J. M. Garibaldi, and C. Hodgman. “Parameter Estimation Using Metaheuristics in Systems Biology: A Comprehensive Review”. IEEE/ACM Transactions on Computational Biology and Bioinformatics 9.1 (2012), p. 185202.

[262] Nathaniel J. Linden, Boris Kramer, and Padmini Rangamani. “Bayesian parameter estimation for dynamical models in systems biology”. PLOS Computational Biology 18.10 (2022). Ed. by Jeffrey J. Saucerman, e1010651.

[263] Kenneth Tran et al. “A Thermodynamic Model of the Cardiac Sarcoplasmic/Endoplasmic Ca2+ (SERCA) Pump”. Biophysical Journal 96 (5 2009), pp. 2029–2042.

[264] N P Smith and E J Crampin. “Development of models of active ion transport for whole-cell modelling: cardiac sodium potassium pump as a case study”. Progress in biophysics and molecular biology 85 (2-3 2004), pp. 387–405.

[265] Jose Guerra, Huan Rui, and Benot Roux. “Multistate Kinetic Model of the SodiumPotassium ATPase”. The Journal of Physical Chemistry B 129.38 (2025), p. 96099621.

[266] Yoram Rudy and Jonathan R Silva. “Computational biology in the study of cardiac ion channels and cell electrophysiology”. Quarterly Reviews of Biophysics 39 (01 2006), p. 57.

[267] T R Shannon et al. “A mathematical treatment of integrated Ca dynamics within the ventricular myocyte”. Biophysical Journal 87 (5 2004), pp. 3351–3371.

[268] Sérgio Filipe Sousa et al. “Application of quantum mechanics/molecular mechanics methods in the study of enzymatic reaction mechanisms”. WIREs Computational Molecular Science 7.2 (2017), e1281.

[269] Lung Wa Chung et al. “The ONIOM Method and Its Applications”. Chemical Reviews 115.12 (2015), pp. 5678–5796.

[270] Jiali Gao et al. “A Generalized Hybrid Orbital (GHO) Method for the Treatment of Boundary Atoms in Combined QM/MM Calculations”. The Journal of Physical Chemistry A 102.24 (1998), pp. 4714–4721.

[271] Mackenzie Taylor, Haibo Yu, and Junming Ho. “Predicting Solvent Effects on SN2 Reaction Rates: Comparison of QM/MM, Implicit, and MM Explicit Solvent Models”. The Journal of Physical Chemistry B 126.44 (2022), p. 90479058.

[272] Christopher I. Bayly et al. “A well-behaved electrostatic potential based method using charge restraints for deriving atomic charges: the RESP model”. The Journal of Physical Chemistry 97.40 (1993), pp. 10269–10280.

[273] John M. Herbert. “Dielectric continuum methods for quantum chemistry”. WIREs Computational Molecular Science 11.4 (2021), e1519.

[274] Abigail E. Teitgen et al. “Multiscale modeling shows how 2-deoxy-ATP rescues ventricular function in heart failure”. Proceedings of the National Academy of Sciences 121.35 (2024).

[275] Siewert J. Marrink et al. “The MARTINI Force Field: Coarse Grained Model for Biomolecular Simulations”. The Journal of Physical Chemistry B 111.27 (2007), p. 78127824.

[276] Casey T. Andrews and Adrian H. Elcock. “COFFDROP: A Coarse-Grained Nonbonded Force Field for Proteins Derived from All-Atom Explicit-Solvent Molecular Dynamics Simulations of Amino Acids”. Journal of Chemical Theory and Computation 10.11 (2014), p. 51785194.

[277] “Investigating Intrinsically Disordered Proteins With Brownian Dynamics”. Frontiers in Molecular Biosciences 9 (2022).

[278] Pu Liu et al. “A Bayesian statistics approach to multiscale coarse graining”. The Journal of Chemical Physics 129.21 (2008).

[279] H.Jane Dyson and Peter E Wright. “Coupling of folding and binding for unstructured proteins”. Current Opinion in Structural Biology 12.1 (2002), pp. 54–60. ISSN: 0959-440X.

[280] Jonathan R Silva et al. “A multiscale model linking ion-channel molecular dynamics and electrostatics to the cardiac action potential”. Proceedings of the National Academy of Sciences 106.27 (2009), pp. 11102–11106.

[281] Siri C. van Keulen et al. “Multiscale molecular simulations to investigate adenylyl cyclase-based signaling in the brain”. WIREs Computational Molecular Science 13.1 (2022).

[282] Bo Cheng et al. “Nanoscale integrin cluster dynamics controls cellular mechanosensing via FAKY397 phosphorylation”. Science Advances 6.10 (2020), eaax1909.

[283] Tamara C Bidone et al. “Multiscale model of integrin adhesion assembly”. en. PLoS Comput. Biol. 15.6 (2019), e1007077.

[284] Pranay Goel, James Sneyd, and Avner Friedman. “Homogenization of the cell cytoplasm: the calcium bidomain equations”. Multiscale Modeling & Simulation 5 (4 2006), pp. 1045–1062.

[285] Catherine Ta, Dongyong Wang, and Qing Nie. “An integration factor method for stochastic and stiff reaction–diffusion systems”. Journal of Computational Physics 295 (2015), pp. 505–522. ISSN: 0021-9991.

[286] Kyu Il Lee et al. “Brownian dynamics simulations of ion transport through the VDAC”. en. Biophys. J. 100.3 (2011), pp. 611–619.

[287] Tamsyn A Hilder and Shin-Ho Chung. “Conductance properties of the inwardly rectifying channel, Kir3.2: molecular and Brownian dynamics study”. en. Biochim. Biophys. Acta 1828.2 (2013), pp. 471–478.

[288] Vikram Krishnamurthy and Shin-Ho Chung. “Large-Scale Dynamical Models and Estimation for Permeation in Biological Membrane Ion Channels”. Proceedings of the IEEE 95.5 (2007), pp. 853–880.

[289] Patricia Bauler, Gary A Huber, and J Andrew McCammon. “Hybrid finite element and Brownian dynamics method for diffusion-controlled reactions”. The Journal of chemical physics 136 (16 2012), p. 164107.

[290] Gary A. Huber et al. “Hybrid finite element and Brownian dynamics method for charged particles”. The Journal of Chemical Physics 144.16 (2016).

[291] R. Delgado-Buscalioni and P. V. Coveney. “Continuum-particle hybrid coupling for mass, momentum, and energy transfers in unsteady fluid flow”. Physical Review E 67.4 (2003).

[292] Léon Bottou, Frank E. Curtis, and Jorge Nocedal. Optimization Methods for Large-Scale Machine Learning. 2018.

[293] “Baseline Model for Predicting Protein-Ligand Unbinding Kinetics through Machine Learning”. Journal of Chemical Information and Modeling 60.12 (2020), pp. 5946–5956.

[294] Minyi Su et al. “Machine-Learning Model for Predicting the Rate Constant of ProteinLigand Dissociation”. Acta Phys.-Chim. Sin. 36.1 (2020), p. 1907006.

[295] Ariane Nunes-Alves, Fabian Ormersbach, and Rebecca C Wade. “Prediction of the Drug-Target Binding Kinetics for Flexible Proteins by Comparative Binding Energy Analysis”. Journal of Chemical Information and Modeling 61.7 (2021), pp. 3708–3721.

[296] Huisi Liu et al. “Public Data Set of Protein-Ligand Dissociation Kinetic Constants for Quantitative Structure-Kinetics Relationship Studies”. ACS Omega 7.22 (2022), pp. 18985–18996.

[297] Yujing Zhao et al. “Machine learning methods for developments of binding kinetic models in predicting protein-ligand dissociation rate constants”. Smart Molecules 1.3 (2023), e20230012.

[298] Yujing Zhao et al. “Mixture-of-Experts Based Dissociation Kinetic Model for De Novo Design of HSP90 Inhibitors with Prolonged Residence Time”. Journal of Chemical Information and Modeling 64.22 (2024), pp. 8427–8439.

[299] Shuo Zhang et al. “Sequence-based Drug-Target Complex Pre-training Enhances Protein-Ligand Binding Process Predictions Tackling Crypticity”. bioRxiv (2025).

[300] John Jumper et al. “Highly accurate protein structure prediction with AlphaFold”. en. Nature 596.7873 (2021), pp. 583–589.

[301] Joseph L. Watson et al. “De novo design of protein structure and function with RFdiffusion”. Nature 620.7976 (2023), pp. 1089–1100. ISSN: 1476-4687.

[302] J. Capela et al. “Comparative Assessment of Protein Large Language Models for Enzyme Commission Number Prediction”. BMC Bioinformatics 26.68 (2025).

[303] Tobias Harren et al. “Modern machine-learning for binding affinity estimation of proteinligand complexes: Progress, opportunities, and challenges”. WIREs Computational Molecular Science 14.3 (2024), e1716.

[304] Pankaj Kumar et al. “Update of KDBI: Kinetic Data of Bio-molecular Interaction database”. en. Nucleic Acids Res. 37.Database issue (2009), pp. D636–41.

[305] Yunpeng He et al. “KDBI-RP: Kinetic Data of RNA-Protein Interactions Database”. Journal of Molecular Biology 437.21 (2025), p. 169357. ISSN: 0022-2836.

[306] Gaurav K Ganotra and Rebecca C Wade. “Prediction of Drug-Target Binding Kinetics by Comparative Binding Energy Analysis”. ACS Medicinal Chemistry Letters 9.11 (2018), pp. 1134–1139.

[307] Martina Bertazzo et al. “Machine Learning and Enhanced Sampling Simulations for Computing the Potential of Mean Force and Standard Binding Free Energy”. Journal of Chemical Theory and Computation 17.8 (2021), pp. 5287–5300.

[308] Zineb Belkacemi et al. “Chasing Collective Variables using Autoencoders and biased trajectories” (2021).

[309] Judith Bernett et al. “Guiding questions to avoid data leakage in biological machine learning applications”. Nature Methods 21.8 (2024), pp. 1444–1453.

[310] Evan N. Feinberg et al. “PotentialNet for Molecular Property Prediction”. ACS Central Science 4.11 (2018), p. 15201530.

[311] Robert A. Copeland, David L. Pompliano, and Thomas D. Meek. “Drug–target residence time and its implications for lead optimization”. Nature Reviews Drug Discovery 5.9 (2006), pp. 730–739. ISSN: 1474-1784.

[312] Peter J. Tummino and Robert A. Copeland. “Residence Time of Receptor-Ligand Complexes and Its Effect on Biological Function”. Biochemistry 47.20 (2008). PMID: 18412369, pp. 5481–5492.

[313] Adriaan P. Ijzerman and Dong Guo. “DrugTarget Association Kinetics in Drug Discovery”. Trends in Biochemical Sciences 44.10 (2019), p. 861871.

[314] Gonzalo Hernandez-Hernandez et al. “A computational model predicts sex-specific responses to calcium channel blockers in mammalian mesenteric vascular smooth muscle”. eLife 12 (2024).

[315] Mattia Bernetti et al. “Kinetics of Drug Binding and Residence Time”. Annual Review of Physical Chemistry 70.1 (2019), p. 143171.

[316] Cleary SR et al. “Inhibitory and stimulatory micropeptides preferentially bind to different conformations of the cardiac calcium pump”. J Biol Chem (2022).

[317] Nikolaienko R et al. “Cysteines 1078 and 2991 cross-linking plays a critical role in redox regulation of cardiac ryanodine receptor (RyR)”. Nature Communications (2023).

[318] Martin Hemberg, Sophia N. Yaliraki, and Mauricio Barahona. “Stochastic Kinetics of Viral Capsid Assembly Based on Detailed Protein Structures”. Biophysical Journal 90.9 (2006), p. 30293042.

[319] David Sept, Adrian H Elcock, and J.Andrew McCammon. “Computer simulations of actin polymerization can explain the barbed-pointed end asymmetry”. Journal of Molecular Biology 294.5 (1999), p. 11811189.

[320] Li Chen et al. “Mathematical models for cell migration: a non-local perspective”. Philosophical Transactions of the Royal Society B: Biological Sciences 375.1807 (2020), p. 20190379.

[321] Adrian H Elcock. “Models of macromolecular crowding effects and the need for quantitative comparisons with experiment”. Current Opinion in Structural Biology 20.2 (2010), p. 196206.

[322] Sandro Bottaro, Tone Bengtsen, and Kresten Lindorff-Larsen. Integrating Molecular Simulation and Experimental Data: A Bayesian/Maximum Entropy Reweighting Approach. Springer US, 2020, p. 219240. ISBN: 9781071602706.

[323] Colin D. Kinz-Thompson, Korak Kumar Ray, and Ruben L. Gonzalez. “Bayesian Inference: The Comprehensive Approach to Analyzing Single-Molecule Experiments”. Annual Review of Biophysics 50.1 (2021), pp. 191–208.

[324] John Moult. “A decade of CASP: progress, bottlenecks and prognosis in protein structure prediction”. Curr. Opin. Struct. Biol. 15.3 (2005), pp. 285–289.

